# Motion-Induced Blindness as a Noisy Excitable System

**DOI:** 10.1101/2022.11.12.516289

**Authors:** Mikhail Katkov, Alexander Cooperman, Noya Meital-Kfir, Dov Sagi

## Abstract

Perceptual disappearance of a salient target induced by a moving texture mask (MIB: Motion Induced Blindness) is a striking effect, currently poorly understood. Here, we investigated whether the mechanisms underlying MIB qualify as an excitable system. Excitable systems exhibit fast switches from one state to another (e.g., visible/invisible) induced by an above-threshold perturbation and stimulus-independent dynamics, followed by a refractory period. In the experiments, disappearance was induced by masks consisting of slowly rotating radial bars with a gap at the target location, leading to periodic perturbation of the visual field around the target (a bright parafoveal spot). When passed around the target location, masks frequently induced an abrupt target disappearance, pointing to locality. As expected from excitable systems, the disappearance time was not affected by additional bars crossing the target during invisibility, and there was little dependence on the mask configuration. After the target reappeared, it stayed for at least 0.5-2 seconds (the refractory period). Therefore, the mechanisms governing MIB represent an example of an excitable system, where the transition to the invisible state is induced by the mask, with the dynamics that follow determined mostly by the internal network properties.

## Introduction

Motion-Induced Blindness (MIB) is a striking phenomenon where a moving mask suppresses the perception of a physically present static target (Bonneh et al., 2001). MIB disappearances are an all-or-none phenomenon and can last up to several seconds, even with a high-contrast target located near fixation (Bonneh et al., 2014). Furthermore, invisibility can be perturbed by a similar visible cue flashed at the target neighborhood (Meital-Kfir & Sagi, 2018), indicating that the neuronal networks involved in invisibility preserve the sensitivity. Despite the detailed experimental exploration of MIB, the underlying mechanism remains unknown. Here, we consider the application of the theory of dynamical systems, which was found useful in describing bistable phenomena.

MIB, with the corresponding spontaneous transition between seen and not-seen states, is often treated as a bistable phenomenon (Bonneh et al., 2014; Devyatko et al., 2017; Hsu et al., 2006; Jaworska & Lages, 2014). MIB was found to be affected by stimulation parameters in a way similar to that of binocular rivalry; the two phenomena show a highly correlated pattern of perceptual transitions across individuals (Carter & Pettigrew, 2003). Others point to large individual differences in MIB measurements as a modeling challenge, arguing that “normative models of MIB may not be practical” (Sparrow et al., 2017).

One interesting approach to describe bistable phenomena involves modeling a stable percept as an attractor state, which is the persistent activity of a group of neurons (Braun & Mattia, 2010; Cao et al., 2021). In this model, the irregular switching between percepts is governed by intrinsic noise, which drives the network state between the two attractors. The mechanism underlying switching can be described as follows: there is a basin of attraction around attractor states, meaning that in the absence of noise, the system would return to the attractor state when perturbed, making switching impossible. However, the presence of noise introduces random perturbations that can move the system outside of the current basin of attraction into the basin of another attractor.

When an external periodic perturbation or signal is introduced, an intriguing phenomenon can occur. The external signal effectively alters the relative sizes of the basins of attraction for the two attractor states. A smaller basin of attraction increases the probability of escaping from it. When the size change is significant, there is a marked difference in switching probability during different phases of the signal. The level of noise plays a crucial role in determining the system’s behavior. For instance, with a small amount of noise, the perturbation caused by it is weak, resulting in a low probability of switching during all phases. Conversely, in the presence of high levels of noise, the system dynamics are predominately governed by noise, leading to similar switching behavior across all phases. However, when the noise level is intermediate, the switching probability varies substantially across different phases of the signal. Here, switching predominantly occurs during the favorable phase of the signal. This phenomenon, characterized by an increased frequency component in switching behavior corresponding to the stimulus frequency, is known as stochastic resonance (refer to the supplementary material for an illustration).

An important difference between MIB and other bistable phenomena, such as binocular rivalry, lies in the number of stable states. Whereas in bistable phenomena two percepts may equally last for prolonged periods, in MIB the invisibility state is unstable, exhibiting a transient behavior. In dynamical systems theory, such behavior characterizes excitable systems. For example, consider a spiking neuron. In the absence of a strong input, the neuron is in a resting state – small input fluctuations lead to small fluctuations in the membrane potential, tracking the input frequency. However, when the input exceeds a threshold level, the neuron responds with a large-scale excursion in phase space (action potential). Once an action-potential is initiated, the membrane potential changes substantially and is weakly dependent on the input. The temporal profile of the spike largely depends on the properties of the neuron, but not on the properties of the input. When the action-potential comes to an end, there is a measurable refractory period during which the neuron cannot emit spikes.

In a manner similar to bistable system, excitable systems can also exhibit stochastic resonance. In the absence of noise or external perturbation, the dynamics of a bistable system converge to one of the stable states, whereas an excitable system would converge to its resting state. In the presence of a sufficient amount of noise, a bistable system switches between stable states, whereas an excitable system repeatedly undergoes a large trajectory; each trajectory ends at its resting state. The detailed dynamics underlying these transitions, which depend on the strength of the noise and the properties of the system, can be explored using periodic external stimulation. A stimulation period that is close to the characteristic system time constant is expected to facilitate switching at a stimulation frequency. In contrast, when the stimulation frequency is too high, the system, while in transit toward its stable state, is insensitive to external stimulation and is not expected to be affected by the frequent incoming stimulation. When the stimulation frequency is too low, since intrinsic dynamics is faster than the driving frequency, several switches are possible during a single stimulation period that broadens the response in the frequency domain. Overall, this looks like a noise-assisted resonance – there is a specific noise-dependent frequency of stimulation that leads to optimal switching (Gammaitoni et al., 1998; Muratov et al., 2005).

Bistable models assume that perception corresponds to the proximity of the dynamical system state to one of the attractors. However, when considering perception in an excitable system, this assumption needs to be clarified. Regarding motion-induced blindness (MIB), the perceptual disappearance of a target does not align with any specific attractor state. We can draw a parallel to memory retrieval in an attractor neural network, where retrieval occurs when the system approaches one of its attractor states associated with a memory. Similarly, in MIB, when the system diverges sufficiently from the attractor state representing the target and there are no nearby attractor states near the system trajectory, the target perceptually disappears. Under this interpretation, within the limit of low noise, we can anticipate several characteristics: (1) After the system is excited, there will be a long excursion time determined by the system dynamics before it relaxes to a visible resting state. This results in a substantial, non-zero duration of invisibility; (2) There will be a refractory period, which represents the minimal visibility time required before the target can disappear again.; and (3) The switching frequency, measured as the signal-to-noise ratio at the stimulation frequency, will exhibit a non-monotonic dependency on the mask period. Regarding high noise levels, we still expect a substantial mean visibility period, and the dependence of the signal-to-noise ratio on the mask period may be less pronounced or even disappear together with the refractory period.

In our study, we employed two types of stimulation to investigate motion-induced blindness (MIB):

1. Static Mask: We used a fixed mask that did not change over time or an absent mask, which resulted in perceiving the Troxler effect. The Troxler effect refers to the perceptual disappearance of a static target when it is presented away from the viewer’s fixation point, as described by Troxler in 1804.
2. Periodic Stimulation: We employed periodic stimulation to study the dynamics of MIB while considering the mask as a driving force. This allowed us to explore how the presence of a moving mask influences the perceptual switches in MIB.

We found that approximately half of the observers experienced a few periods of invisibility shorter than 400 ms, indicating the presence of a minimum duration of invisibility (similar to the findings by Meital-Kfir et al., 2016). On the other hand, the remaining half of observers exhibited very short invisibility periods even under conditions without a moving mask. This suggests that a significant portion of the switches observed were not directly related to the presence of the moving mask. Furthermore, for most participants, the visible periods were not shorter than 1-2 seconds, indicating a relatively longer duration of visibility.

In our analysis of the switching records, we examined the signal-to-noise ratio (SNR) and observed a non-monotonic relationship between the SNR and the stimulation frequency. This suggests that the switching behavior in MIB is influenced by the frequency of the applied stimulation. To model the distribution of switching times, we found that a combination of two random variables provided a good fit. The first variable followed a Gamma distribution, which captured the statistical properties of the internal dynamics underlying the switches. The second variable followed a Gaussian distribution and represented the minimal delay due to neuronal dynamics and motor response jitter when reporting a disappearance. Note that the Gamma distribution alone was insufficient to accurately describe the observed switching times.

## Results

To efficiently track the dependence of perceptual transitions on a mask structure, we employed masks consisting of discrete, well-defined parts. The mask (Figure 1) consisted of bars rotating around a fixation point (with breaks inserted to avoid physical interference with the target); the number of bars and their speed was varied to control the mask angular frequency (see the Methods). The important timescale is the time interval between consecutive events where a bar passing the target shows indistinguishable displayed images. Therefore, we report the period of the mask as the minimal time between identical image appearances on the screen.

**Figure 1.**
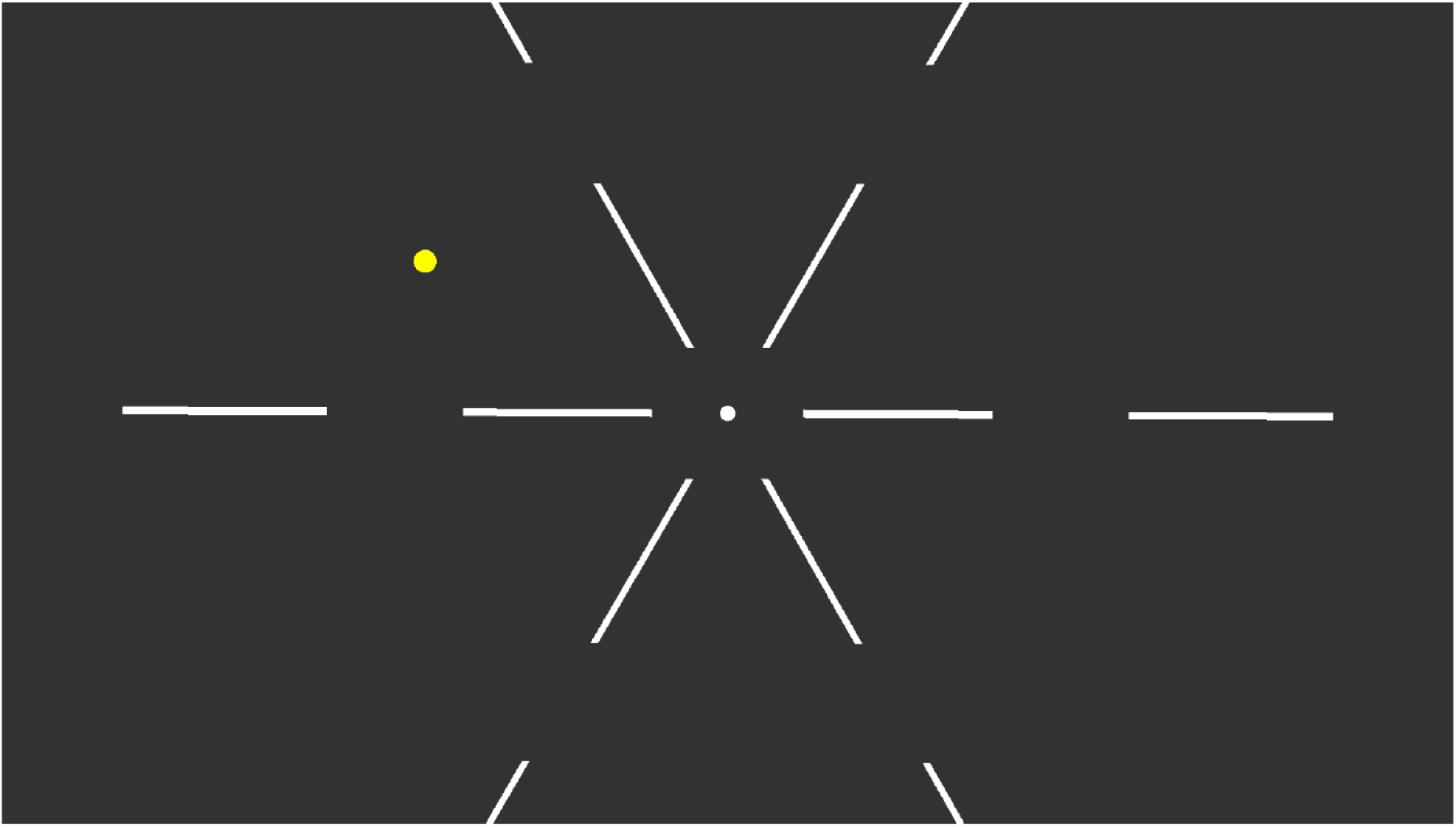
An example of stimuli used in the experiments. A screenshot of stimulus shown on a 24” display that was viewed from a distance of 120 cm. In the experiment the mask consisted of white bars, rotating around the fixation point, thus leading to periodic stimulation; the time period depended on the rotation speed and the number of bars.

*Figure 2* depicts the dependency of the perceptual state on the phase of the mask for two observers. Clearly, the disappearance report (the black lines) lacks a structure for short periods (1 sec) of mask rotation, but it was phase locked for longer periods (4 sec). The results for all observers are shown in the supplementary materials. For the longer rotation period (4 sec), the target was reported as visible during some phases of the stimulation cycle on practically every mask cycle, whereas during other phases, visibility reports were recorded on only 65-80% of the cycles. This indicates that the moving bars are effective inducers of MIB. In contrast, with the shorter mask period (1 sec), the invisibility reports were uniformly distributed during the stimulation cycle, indicating that the system does not have sufficient time for relaxation from the invisible state. The simple mathematical properties of this mask enable a systematic study of the specific mechanisms governing disappearances. Moreover, at slow rotation speeds, this mask can be used to study interactions across the boundary of awareness, since for slow presentation periods the time of invisibility is highly predictable.

**Figure 2.**
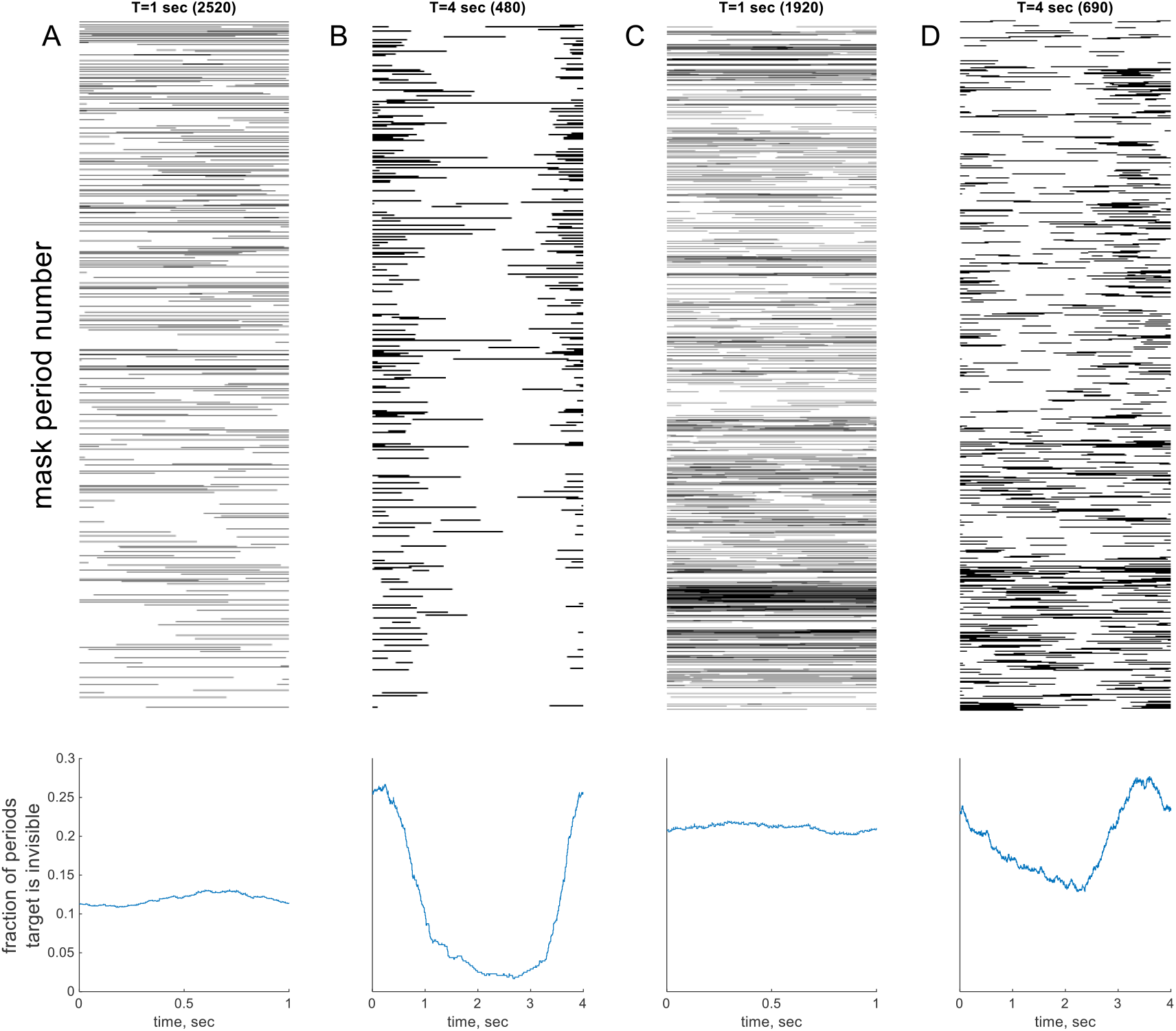
Examples of disappearance patterns from 2 participants and 2 stimulation frequencies. Top row: A raster plot of the invisibility reports. The black regions represent moments when the target was reported absent. Consecutive periods of mask rotation are stacked vertically. Bottom row: The disappearance rates of a target as a function of time within one stimulation cycle. (A) and (B) are the results from one human observer for mask periods of 1 sec (A) and 4 sec (B). (C) and (D) the same as (A) and (B) obtained from another human observer. One can observe that the disappearance rate toward the end of the cycle for both observers approaches 25%, indicating the high effectiveness of the mask to induce MIB. In contrast, in the middle of the cycle for slow mask periods (B, D) there are reduced disappearance rates. For example, in (B) the target is practically always visible in the middle of the cycle.

To determine whether a refractory period is present, we examined the distribution of visible periods (Figure 3, and Supplementary material). One can see that all visible periods in this example are longer than 800 miliseconds, except one. This is not explained by limits on the reporting speed, since there are many invisible periods with shorter durations. Within the context of the current theory, this minimal duration is due to the system’s refractory period, that is, the system needs to recover from one invisibility epoch before transitioning to a new invisibility epoch. That is, when the target becomes visible, it will stay visible for some minimum time. The time interval distributions for visible and invisible periods for all observers are depicted in Supplementary Figure S3 1-6. Of the six observers, four show evidence of a refractory period when excluding a few very brief visibility events that could potentially be attributed to accidental key presses or releases during the reporting of target visibility, or rare short events influenced by the presence of noise. For observers O4 and O6, as well as observer O2, when the mask has a period of 1 second, there is a notable occurrence of numerous short visibility events. This behavior aligns with what would be expected in an excitable system with a significant level of noise, as supported by the simulations presented in the Supplementary material. However, we acknowledge that it is possible that a different mechanism may be responsible for perception in these observers.

**Figure 3.**
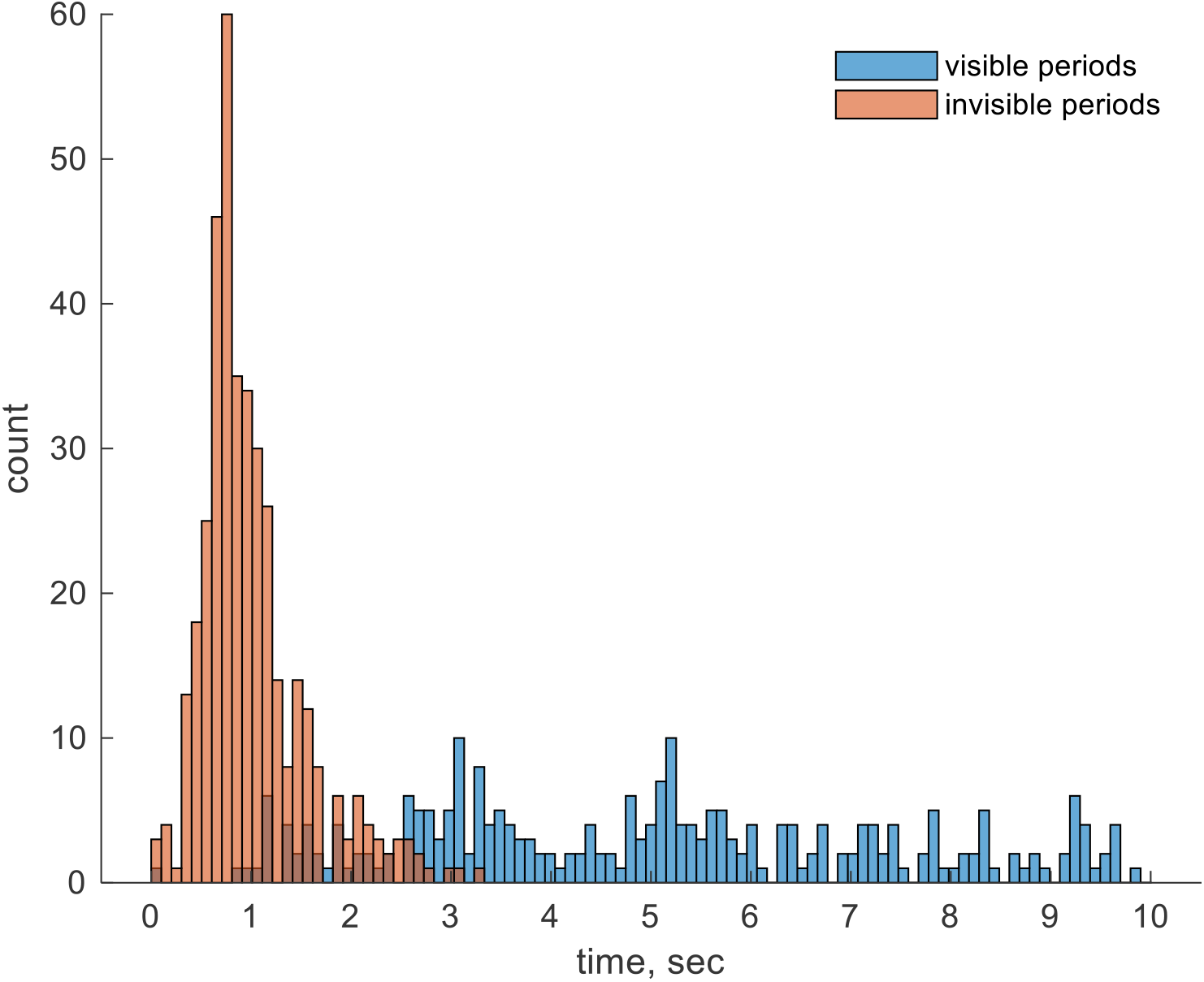
Example of the distributions of visible and invisible periods for one observer and one condition.

Previous studies have explored the distribution of bistable dominance time, which refers to the duration of a single percept before switching to another percept. Some of these studies have attempted to model this distribution using a Gamma distribution (Brascamp, van Ee, Pestman, & van der Berg, 2005; Leopold & Logothetis, 1996). However, it has been reported that the data are not always well described by a Gamma distribution.

Similarly, in our own experiments, we also did not obtain a good fit using a Gamma distribution. In our experimental setup, in addition to the underlying decision process, there is also a motor reporting component, which is often modeled as a random variable with a Gaussian distribution. Taking this into consideration, we hypothesized that the combination of two random variables, one with a Gamma distribution representing the decision process and the other with a Gaussian distribution representing the motor reporting, may provide a better fit to the data.

In 140 out of 148 conditions, the distribution of the invisible periods is well described by the sum of gamma distributed and normally distributed random variables, assuming a 5% criterion for the Kolmogorov-Smirnov test. The normally distributed random variable represents the noisy reaction time for pressing and releasing the keyboard space bar, and the average transition time through the excited state back to the resting state (see Table 1, and the supplementary material). One can also observe that the Gaussian distribution parameters are relatively consistent for the same observer in all experimental conditions, probably indicating prototypical dynamics. The gamma distribution parameters are more variable, even within the same observer, possibly indicating the differential effect of the mask. For instance, one can observe that the values of κ (the shape parameter, see Eq.1 in the Methods) are more consistent in conditions with moving masks than in the static conditions (e.g. Troxler, see the condition details in the Methods). We found that the estimated distributions cannot be rejected using Kolmogorov-Smirnoff statistics, as detailed in the Supplementary material. Overall, considering all the experiments, the fraction of the fitted distributions that can be rejected is within the rejection criterion p-value, indicating that the mixture model provides an excellent description of our data.

**Table 1.**
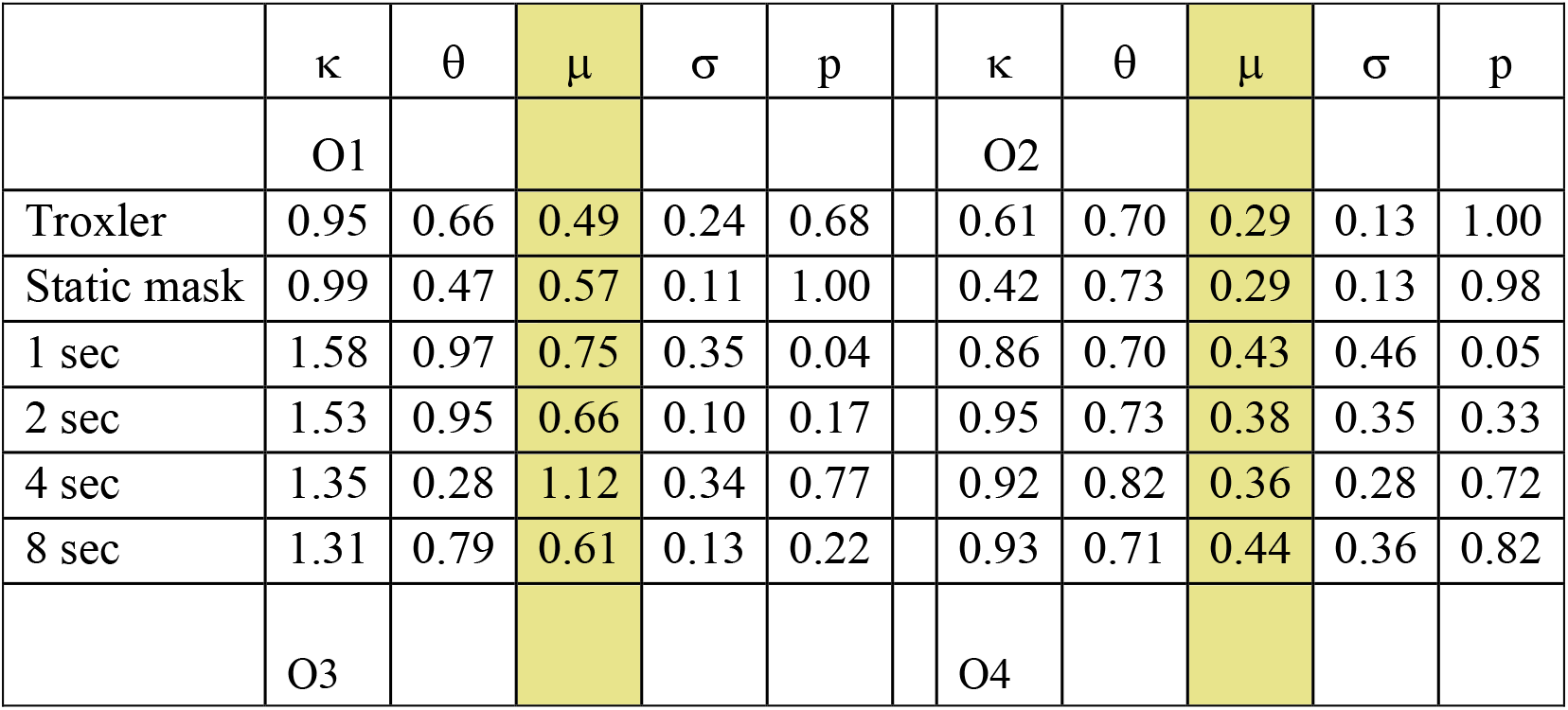
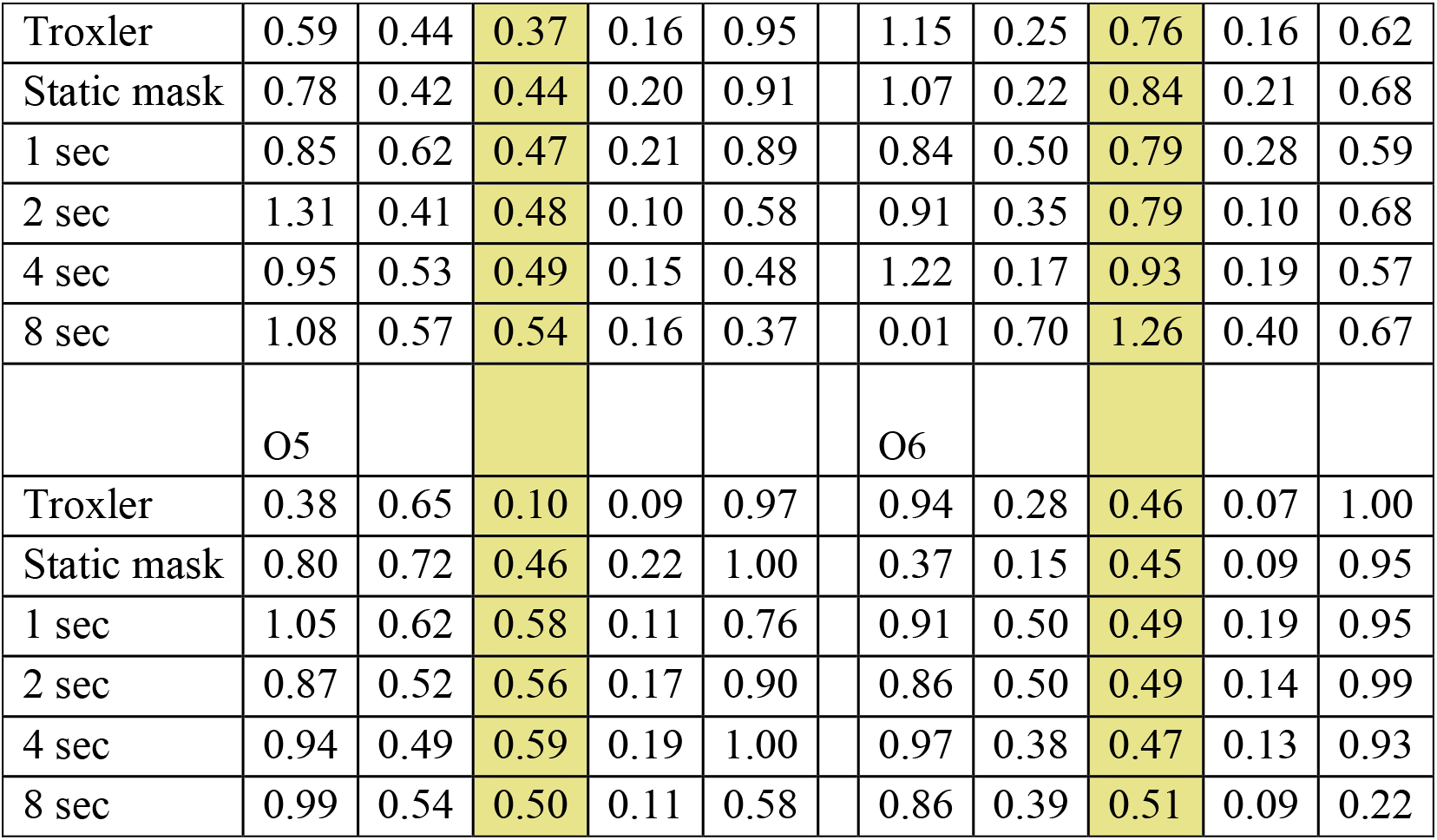
The distribution parameters for the disappearance times. The distributions were assumed to be the sum of two random variables, one normally distributed and one gamma distributed. The symbols *κ* and *θ* represent the shape and scale parameters of the gamma distribution; *μ* and *σ* are the mean and SD of the normal distribution, respectively. p represents the p-value of the Kolmogorov-Smirnoff goodness of fit. All six observers are shown (3 rows X 2 columns). The highlighted columns show the constant delay before the target reappears, representing a long excursion time (∼0.5 sec) in an excitable system until it returns to a stable state. One can see that these times are relatively consistent for the same observer in all experimental conditions, probably indicating prototypical dynamics. The symbol *σ* probably indicates the jitter in the response. The parameters of the gamma distribution are more variable, even within the same observer, possibly indicating the differential effect of the mask. For instance, one can see that the *κ* values are more consistent in conditions with a moving mask than in the Troxler or static mask conditions (see the condition details in the Methods).

We have presented evidence supporting the existence of a refractory period following the invisibility state, which is clearly observed in half observers (Supplemental Figures S3 1-6). Additionally, we have observed a delay in the dynamics before the system returns to the resting state. Next, we show the presence of noise-assisted resonance. Figure 4 depicts the signal-to-noise ratio (SNR, see the Methods for details) for each observer under different rotating mask periods. As one can see, for most observers there is a non-monotonic dependence of SNR on the mask frequency. One observer shows monotonic dependence, possibly indicating that the resonance is at a higher frequency.

**Figure 4.**
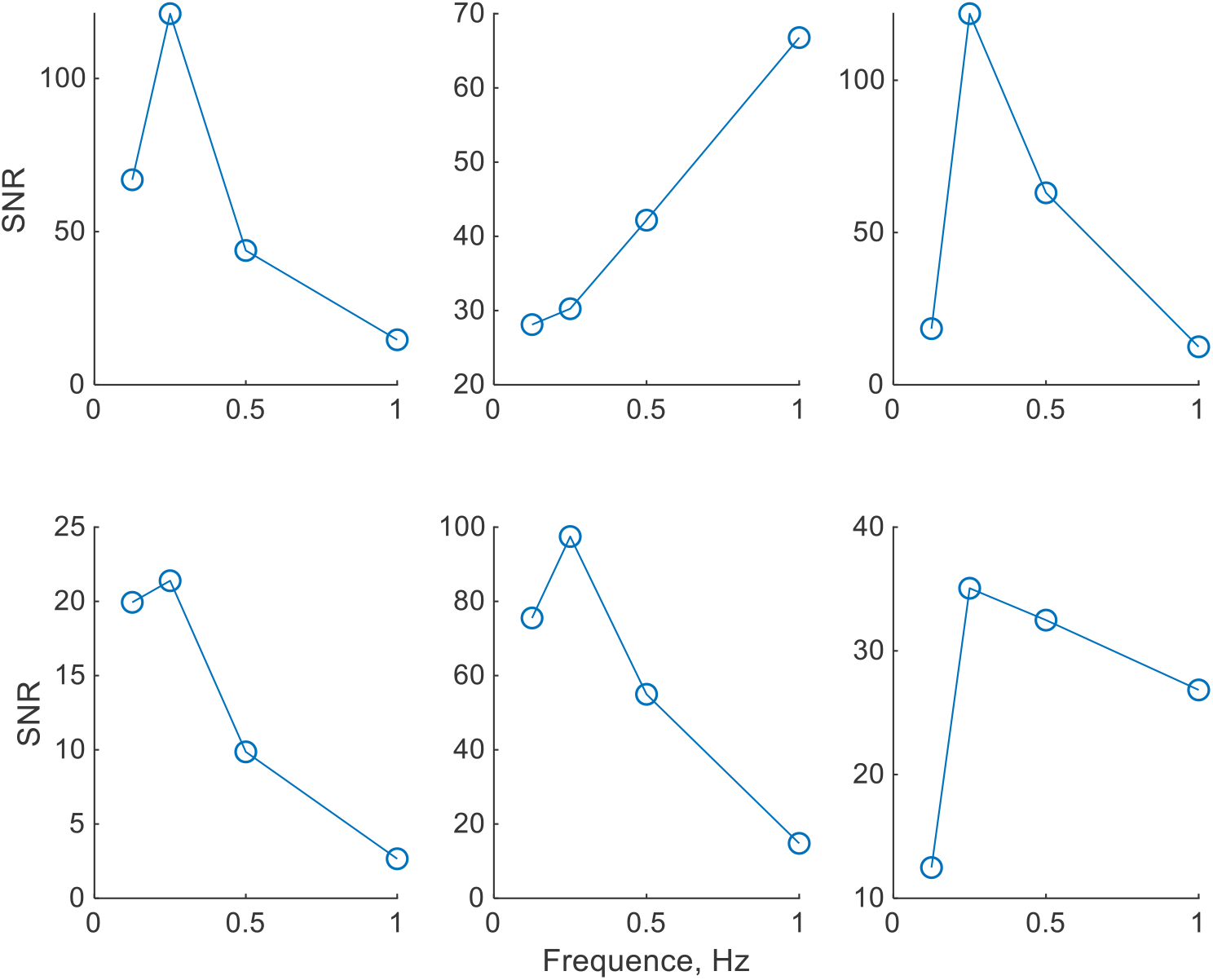
Frequency dependence of the signal-to-noise ratio (SNR). Each plot shows the SNR for a single observer for different mask periods (see details in the Methods).

The phase-locking analysis shows that the effectiveness of the mask is 20-30%. For slow speeds, the disappearance events are concentrated around a specific cycle phase. Additionally, the system spends a characteristic time in the invisible state. By jointly considering these three observations, we can conclude that a bar passing in the vicinity of the target induces a target to disappear, but not always. When the rotation speed is fast, so that consecutive bars cross the target within a single disappearance epoch, a new disappearance event will not take place. Therefore, we would expect the statistics of the inter-disappearance events (the time between two consecutive disappearances) to correspond to the statistics of spontaneous disappearances under the limit of high rotation speed. If the rotation speed is too slow, there will be longer periods of visibility (resting states) between the induced invisible events, allowing for infrequent spontaneous events of disappearance (as in the Troxler effect) when the mask is in a favorable phase. In both cases, the distribution of the inter-disappearance times would be smeared across many timescales and would weakly depend on the stimulation frequency. For intermediate rotation speeds, where the rotation period is near the characteristic timescale of the system, upon a state switch triggered by a favorable stimulus event, the intrinsic dynamics bring the system to a resting state, thus to a response that is time locked to a specific phase and frequency. Since the mask does not induce a disappearance in every period, one would expect the inter-disappearance times to be predominantly multiples of the rotation period.

Figure 5 (top row) shows the inter-disappearance times for one observer. One can see that for mask periods (T) of 2, 4, and 8 seconds, there are peaks of the distributions of the inter-disappearance times corresponding to times nT, where n is an integer number, i.e., it frequently requires several mask periods before the target disappears. One can also observe that close to the resonance frequency (T = 4sec, Figure 5), the second and third times the bar passes the target (peaks at 8 and 12 sec) are more successful in inducing disappearance than the first time is. This is expected if the rotation period is slightly faster than the characteristic timescale of the system. Here, when the system returns to a resting state, the mask has already passed the most favorable phase for switching.

**Figure 5.**
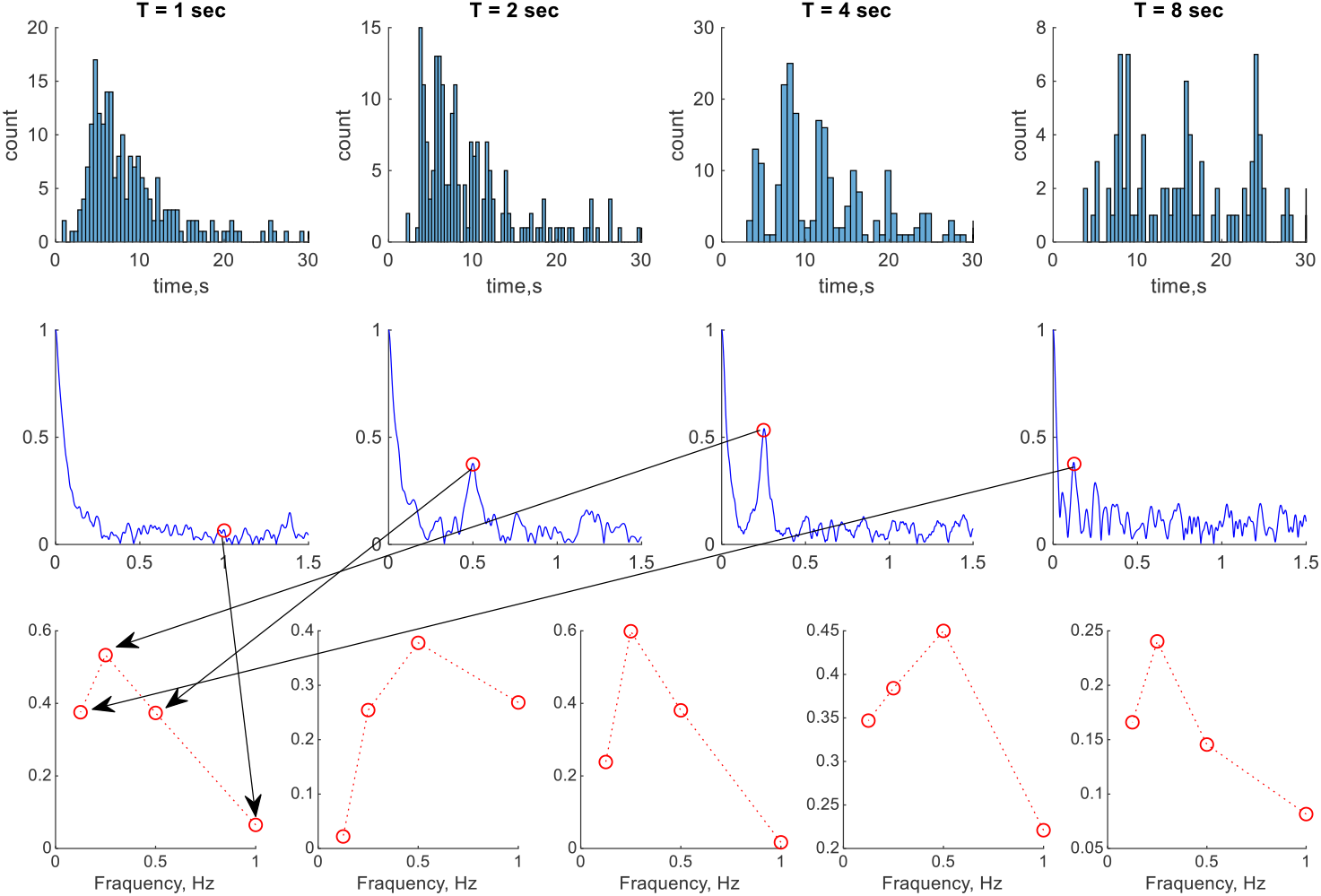
Inter-disappearance times. Top row: histograms of the inter-disappearance times for one observer during different mask periods (indicated in the subplot titles). Middle row: empirical characteristic functions of the distributions shown in the top row. Bottom row: Amplitudes from each characteristic function at the frequency corresponding to the frequency of the mask are shown. Each plot corresponds to the experiments performed by a single observer, with the left one matching the data depicted in the top and middle rows.

Another way to see the periodicity of the function is to take its Fourier transform. A single peaked amplitude spectrum will indicate a single dominant frequency. The characteristic function (Billingsley, 1995) of the distribution is practically the Fourier transform of its probability density function (pdf). Since the pdf shape is not known, one may compute an empirical characteristic function (Cramér, 1946), and if the pdf is dominated by a periodic component, then a peak in the absolute value of the empirical characteristic function is expected. The middle row in Figure 5 shows the absolute value of the empirical characteristic function for samples whose histograms are shown in the top row. The peaks at the stimulation frequencies are clearly seen for mask periods of 2, 4, and 8 seconds (red circles) – confirming that the disappearance events are predominantly spaced by the integer number of periods (nT), with many different values for the integer n. Interestingly, the amplitude of the mode corresponding to stimulation frequency has a non-monotonic behavior) (Figure 5, bottom row for all observers). For example, for the observer shown in the top and middle rows, the peak amplitude for T= 4 sec is larger than for T= 2 sec or T=8 sec.

Distributions and empirical characteristic functions for all observers are presented in the supplementary materials. The data for one observer (O4, examples of raster plots are shown in Figure 2C,D) show no peak in the characteristic functions for any stimulation frequency. These results indicate the domination of noise in the behavior but may also reflect the low efficiency of bars as inducers for this observer. One of the indications for the first case is the similarity of the inter-disappearance distributions between moving bars, static bars, and Troxler conditions (Supplementary Figures Figure S2 1-6).

## Discussion

The series of experiments performed in this work indicate that rotating bars are effective inducers of target disappearance in MIB. Moreover, the effect is phase locked to the mask for long periods (Figure 2). We observed a refractory period for most observers (Figure 3, S3 1-6). For two observers (O4 and O6), phase locking was less pronounced (Figure S.1. 4, Figure S.1. 6), suggesting inefficient stimulation. Signal-to-noise analysis of the temporal sequences of responses showed resonance at frequencies around 0.25Hz, corresponding to a mask period of 4 sec (Figure 4). Analyzing the empirical characteristic functions, we observed peaks at frequencies corresponding to integer multiples of mask periods (Figure 5). In other words, several mask periods are frequently required to induce a disappearance, indicating that moving bars are weak disappearance inducers and that the amount of intrinsic noise is relatively small. Additionally, in all experiments the distribution of invisibility times is well modeled by the sum of two random variables distributed according to gamma and the normal distributions. The Gamma distribution was frequently considered for describing switching dynamics in bistable systems. We added the Gaussian distribution to absorb temporal jitter when executing a motor command to press or release a keyboard key and the mean time required for a long excursion of the system until it returns to a resting state. In many different experiments the mean time in a Gaussian distribution predominantly varied between 1 and 2 seconds, with extreme values ranging from 0.5 sec to 2.5 sec. In the supplementary Materials we present the simulation of an excitable system that shows similar behavior. Therefore, we concluded that the mechanisms responsible for MIB may operate in the regime of an excitable system.

One of the models used in the past to study bistable perception describes perceptual switches by the random motion of a “particle” in a double-well potential. In this model, the decision variable corresponds to a one-dimensional coordinate of a “particle” inside a potential with two local minima (two wells), separated by a barrier. In the absence of noise, the “particle” “falls” to the position of the nearest local minimum and stays there forever. In the presence of noise, however, the “particle” has a chance to gain enough energy to overcome the barrier and move to another well and it will subsequently fall to another local minimum. Translating this to perception, it is usually assumed that when a “particle” is close to one minimum, one image is perceived, and when the “particle” is close to another minimum, another image is perceived. This one-dimensional model was introduced to study binocular rivalry by Kim et al., 2006. Nevertheless, it is hard to extend this formalism to describe the quasi-stable perception in MIB or the Troxler effect (Troxler, 1804). Here we suggest that the theory of excitable systems is a suitable framework for describing MIB. Although the practical implementation in a biological network can be quite complicated, the simplest formal model describing excitable systems is the FitzHugh-Nagumo model (FitzHugh, 1961; Nagumo et al., 1962; Sherwood, 2014). It assumes that the neuronal dynamics can be effectively described by two one-dimensional variables with substantially different timescales of evolution. The variable having fast dynamics can be interpreted as a decision variable in the double-well model. The variable with slow dynamics is assumed to represent processes such as adaptation (Caetta et al., 2007; Gorea & Caetta, 2009), filling-in (Hsu et al., 2006), motion streaks (Wallis & Arnold, 2009), depth ordering, and surface completion (Graf et al., 2002). Within this proposal, there is a perceptual criterion on the “fast” decision variable, so that a target is reported invisible when this criterion is crossed but is reported visible when the criterion crossing yielding a decision is in the opposite direction.

By changing parameters in the same noisy FitzHugh-Nagumo model, one can describe other bistable phenomena such as binocular rivalry. This description can be general enough to describe all stable, bistable and quasi-stable perceptual phenomena. Furthermore, stable fixed points of the dynamical systems, forming attractors, constitute the basis for the memory models in the attractor neural networks (ANN). In these networks, the memory items are assumed to be recalled, or perceived, when the pattern of network activity is close enough to one of the stored patterns of activity (memory). Technically, when the overlap (correlation) between the activity and the stored memory exceeds some threshold value, the model is assumed to recall this stored memory. In other words, the overlap measure can be considered a decision variable. A memory item is then recalled (perceived as this specific item) when the dynamics of the network activity are near the memorized pattern (the fixed point of the dynamics), for example, the resting state in MIB or stable percepts in bistable phenomena. Thus, a possible interpretation of MIB follows, so that both the static target and the moving mask are separately consistent with some attractor states in the brain corresponding to their generated percepts, whereas both of them together are inconsistent with any attractor. Here, the mask plays the role of a force driving the network state away from an otherwise stable attractor that corresponds to the static target perception. It appears that there are no other stable attractors near the target (meaning that there is no stable illusory percept), and that the network dynamics relax (after a long excursion) into the stable state, yielding a visible target. During state transition, there is no stored pattern along the path of the dynamic that can be interpreted as target presence; therefore, the target stays perceptually invisible.

By considering the dynamics of the nonlinear systems near the bifurcation point in the context of the Attractor Neural Network model of memory, we speculate that the conscious state consists of activating a specific attractor associated with a specific memory. Once the network is driven away from the attractor, the brain becomes unaware of the physical stimulus until the network dynamics return to the memory attractor associated with any stimulus. Despite the brain being unaware of the physical stimulus, the stimulus drives the dynamics of the network, allowing an interaction across the boundary of awareness (Meital-Kfir et al., 2016; Meital-Kfir & Sagi, 2018). As in the initial stage of testing these speculations, we established here that MIB operates in a regime similar to that of an excitable system.

## Acknowledgment

This work was supported by the Basic Research Foundation administered by the Israel Academy of Sciences and Humanities (grant No. 6501560).

## Methods

### Human Observers

This study was approved by the Weizmann Institute of Science Ethics Committee and the Helsinki Committee. Ten human observers with normal or corrected-to-normal vision participated in the experiment. Before experimentation, all observers provided their informed consent under the approved Declaration of Helsinki.

Six observers with normal or corrected-to-normal vision participated in all experiments.

### Stimuli

This study included 6 experimental conditions. Stimuli consisted of a static target (yellow dot, 0.5 deg in diameter) placed at 6 deg in the upper-left visual field and a fixation point (white dot 0.25 deg in diameter). These elements were present in all conditions. In the Troxler condition, no additional elements were present on the screen. In the other (number) conditions, there was either a static or a rotating mask. The mask consisted of lines placed along six rays originating near the fixation point and form a 60° angle between each pair of rays. Each ray was composed of two distinguishable lines to avoid local interactions with the target during mask rotation. A similar “protection zone” was also kept between the mask and the fixation point. The inner line started at 1.25 deg of visual angle and ended at 4.6 deg, whereas the outer line started at 7 deg and ended at 10.5 deg. In the rotating mask conditions, the rays were rotated clockwise with constant angular velocity. The angular velocity was chosen so that the periods of motion (i.e., the time between identical images on the screen) were 1, 2, 4 or 8 sec.

### Procedure

After signing a consent form, the observers performed several daily sessions (minimum 6, maximum-12). In each daily session, the observers performed three blocks of trials, 20 minutes each. A mandatory 15-minute break separated the blocks. Each trial was self-initiated by the observer and lasted 120 sec. During the trial, the observers were instructed to fixate on a central fixation point and report when the target is perceptually invisible by pressing and holding the space bar on a computer keyboard until the target becomes visible.

### Empirical characteristic function

The characteristic function is formally defined as

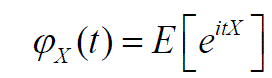

where E[.] is the expected value, 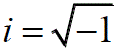 and X are random variables. Empirical distribution functions are computed by substituting samples from X

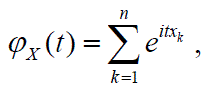

where *X*_*k*_ is the *k* –th data point out of *n*. Absolute values of *φ_X_* (*t*) are shown on graphs. *φ_X_* (0) = 1 by definition, and if X has a periodic density function, there is only one non-zero value of *φ_X_* (*t*) corresponding to the period of density function.

### Fitting procedure

We fitted the disappearance distributions as the sum of two random variables, one having a normal distribution and the other one having a Gamma distribution.

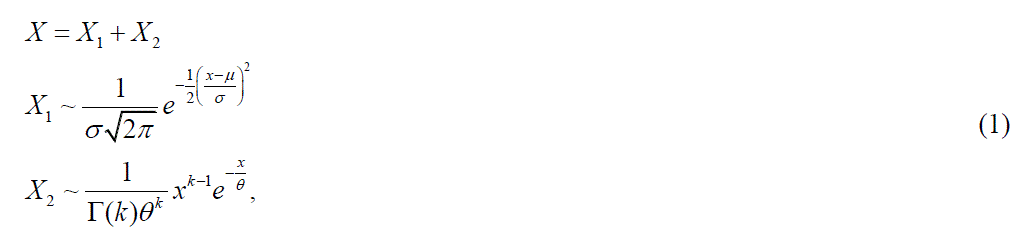

where Γ(*k*) is a gamma function. The Gamma distribution was previously used to fit bistable phenomena (Carter & Pettigrew, 2003; Devyatko et al., 2017). Additionally, we assumed that there is a minimal delay required for the deterministic trajectory to return to a stable fixed point when the system is excited (the mean of the Normal distribution), and that some jitter exists in the motor responses, which we are attempting to capture by varying the variance of normal distribution. Although it is possible to compute the resulting distribution formally, estimations using the resulting formula are unstable; instead, we used many tuples of samples (10^5^ for each set of parameters) where the first elements of a tuple consisted of samples from the normal distribution, and the second element of a tuple consisted of samples from the normal distribution. The values of tuples were added to have a sample from the theoretical distribution, which in turn, were used to form an empirical cumulative distribution, and finally, the intermediate values of the theoretical cumulative distribution function (CDF) were estimated by linear interpolation. Using interpolated CDF, we estimated the p-value of the Kolmogorov–Smirnoff goodness of fit. The obtained p-values were used as a cost function in an optimization procedure that consisted of 2 consecutive runs of Genetic Algorithm (GA (Deb, 1999), as implemented in Matlab ® optimization toolbox), followed by non-linear least-squares optimization. GA was scanned over different parameter values, and following a smooth method, the local minimum near the best set of parameters was found by GA. Repeating this procedure with the best value found in the first round as seed values for a second round of GA increased the chances of finding better solutions. This method does not guarantee that no good fit to the data exists in cases where our procedure failed to find one.

### Signal-to-Noise Ratio analysis

To compute the Signal-to-Noise Ratio (SNR), we created a signal representing the perceptual states as a function of time. The visible target was assigned the value ‘1’ and the invisible target was assigned the value ‘0’ discretized at 1ms precision. All trials were concatenated, sequentially forming a single signal for every observer. SNR was defined as the ratio between peak amplitude at a stimulation frequency within a narrow band (∼2-10mHz) of an expected stimulation frequency and an average amplitude in a wider frequency window (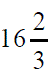 times wider). More specifically, a Fast Fourier Transform was computed on the generated signal and a narrow band was selected as 6 frequency bins around the expected frequency, whereas the wide band contained 100 frequency bins.

The details of exploratory experiment are provided in the Supplementary materials.

### Declaration of generative AI and AI-assisted technologies in the writing process

During the preparation of this work the author(s) used chatGPT in order to improve grammarand text consistency. After using this tool/service, the author(s) reviewed and edited the content as needed and take(s) full responsibility for the content of the publication.

## Supplementary materials

### Disappearance patterns for individual observers

Below are figures of the disappearance patterns for all 6 observers presented in the format of Figure 1. Each column represents one rotation period – 1, 2, 4, and 8 seconds from left to right.

**Figure S.1.1.**
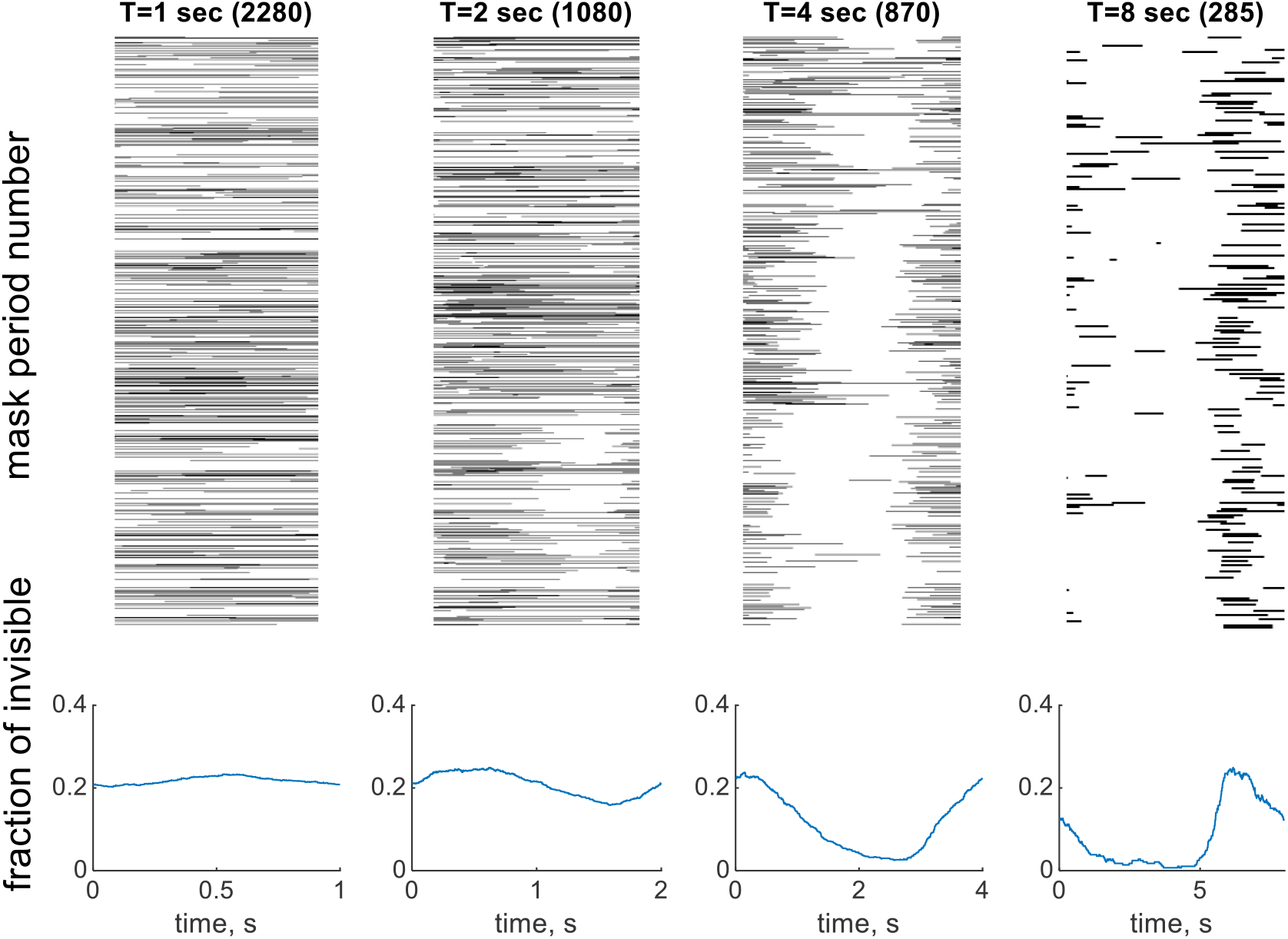
*Disappearance patterns for observer O1*.

**Figure S.1.2.**
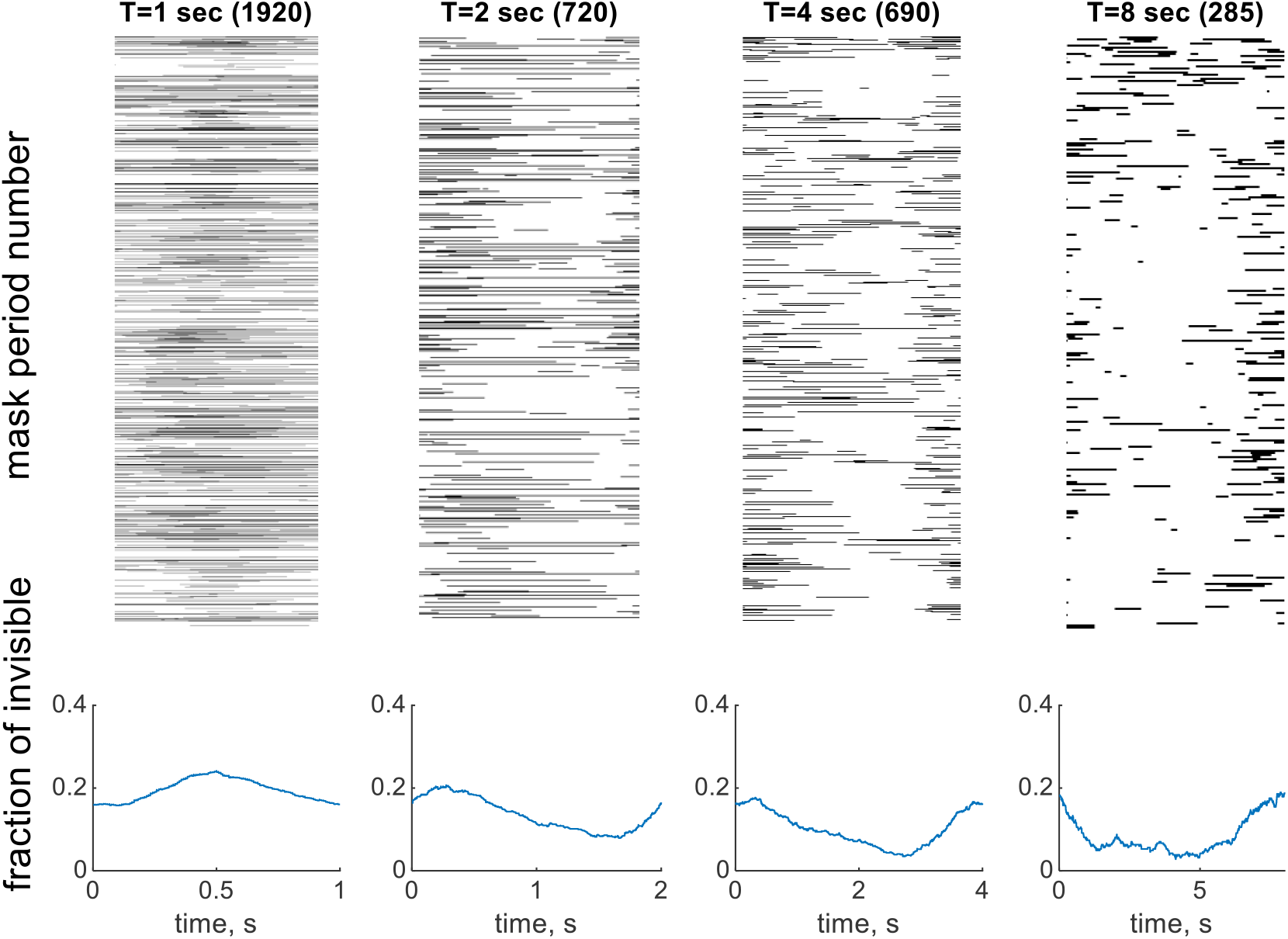
*Disappearance patterns for observer O2*.

**Figure S.1.3.**
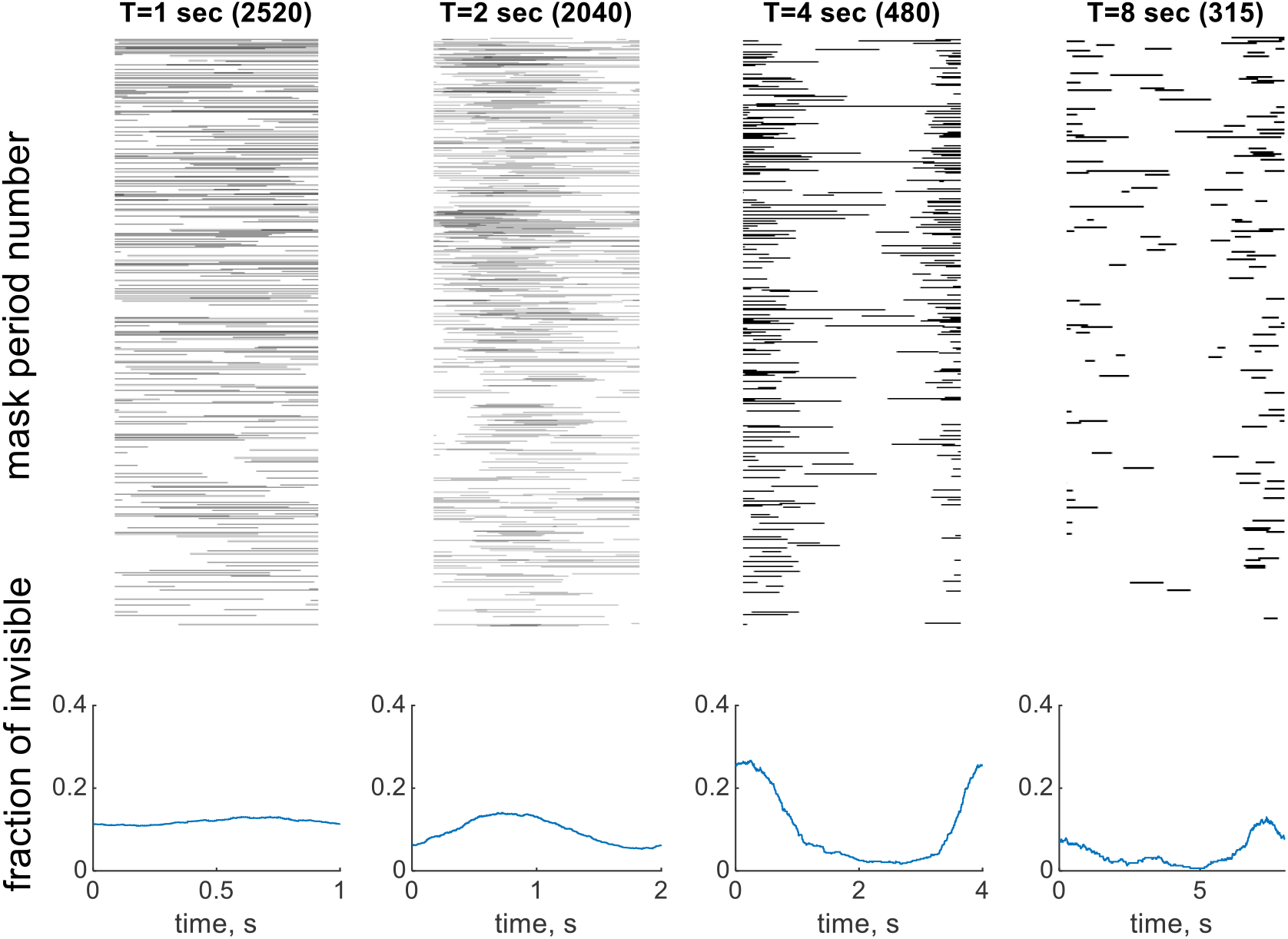
*Disappearance patterns for observer O3*.

**Figure S.1.4.**
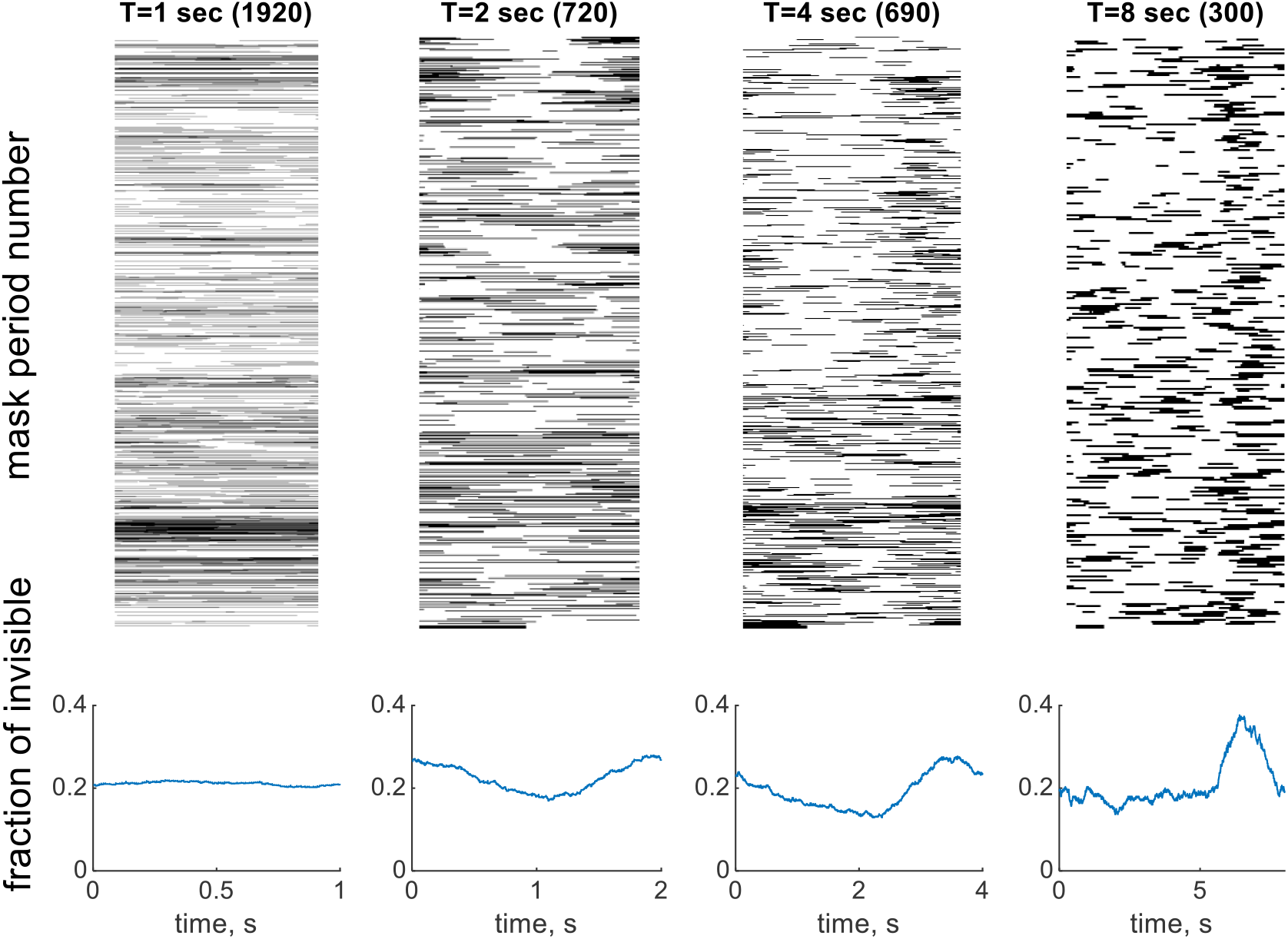
*Disappearance patterns for observer O4*.

**Figure S.1.5.**
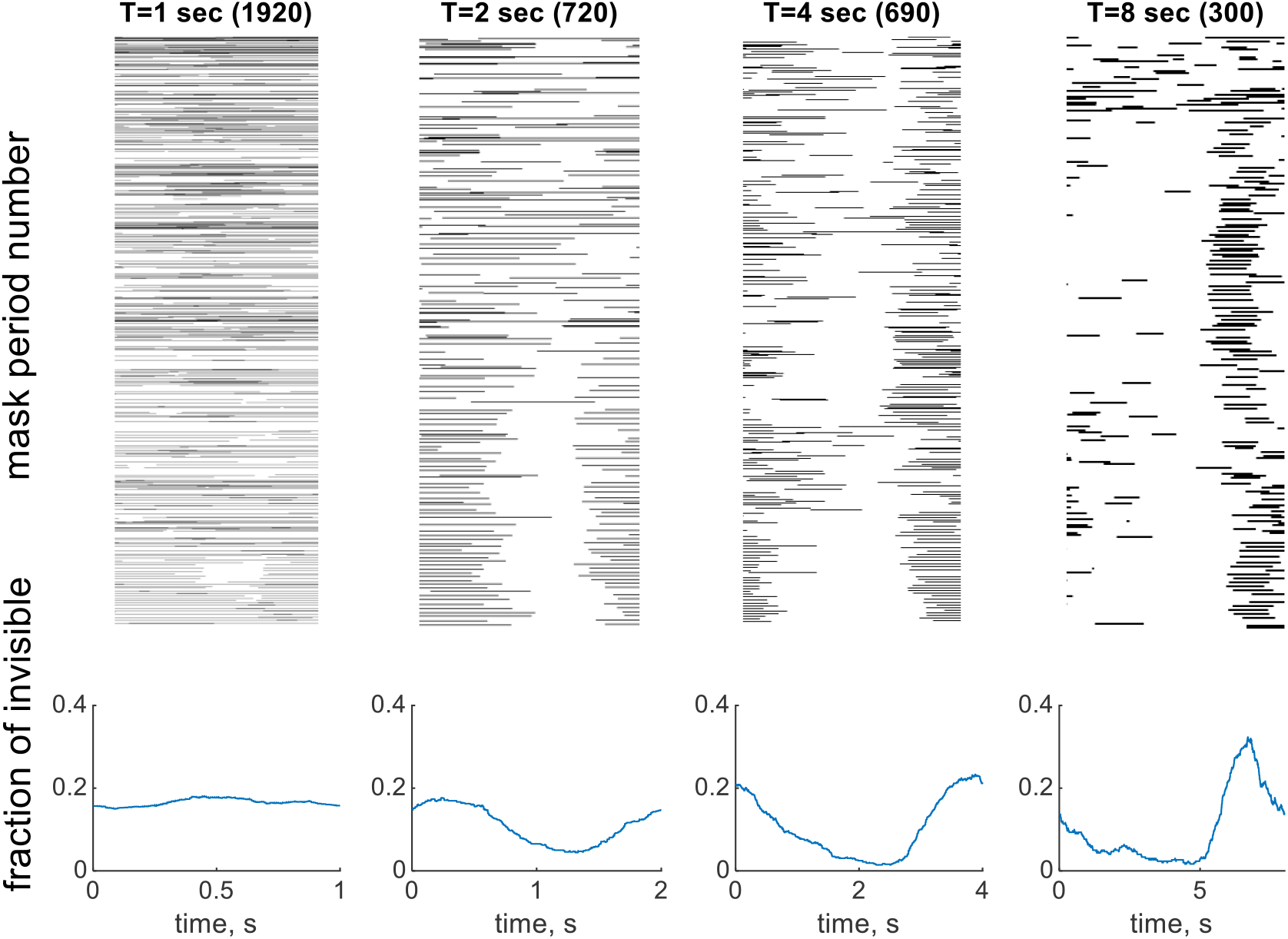
*Disappearance patterns for observer O5*.

**Figure S.1.6.**
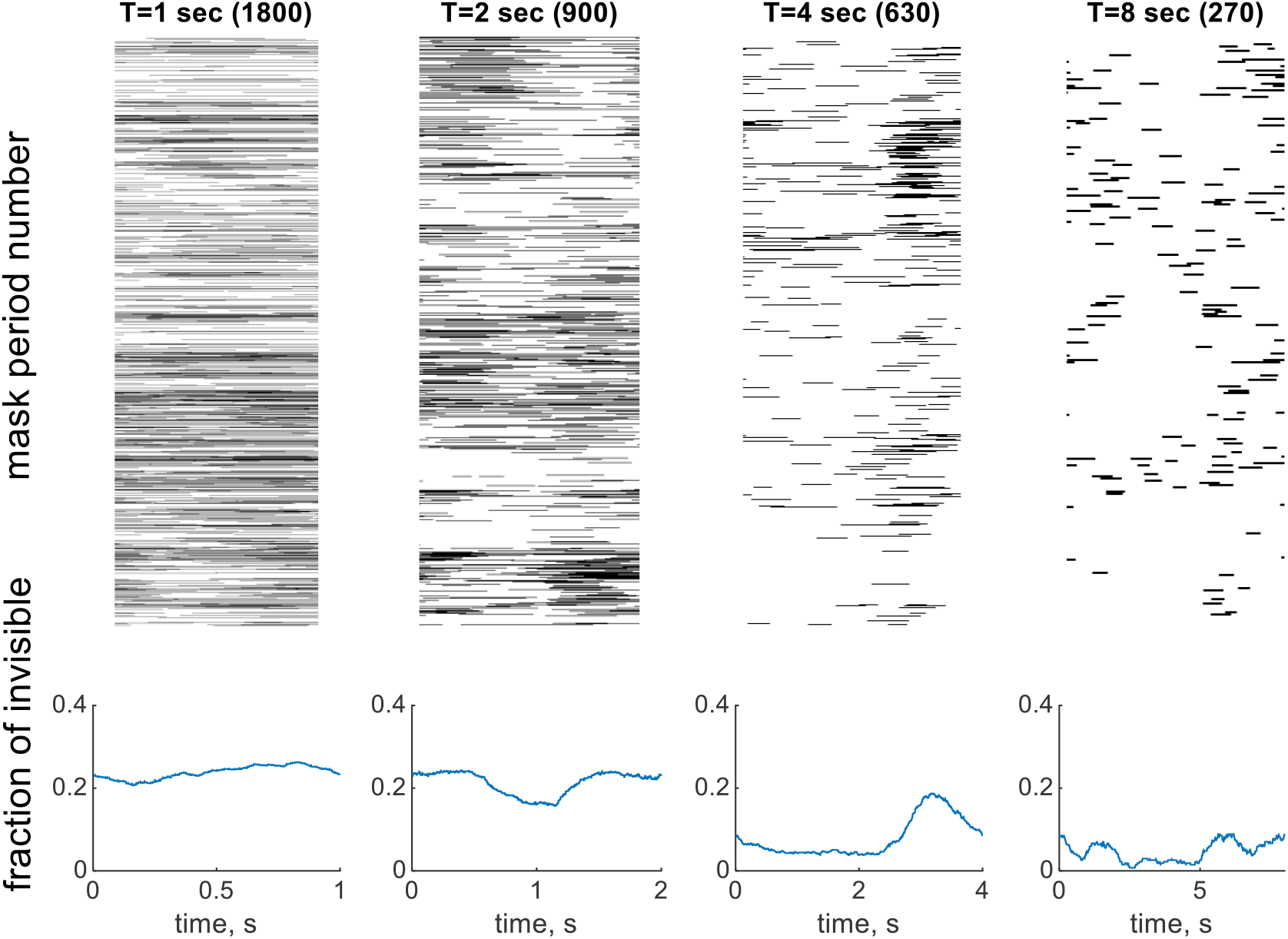
*Disappearance patterns for observer O6*.

### Inter-disappearance times for all observers

Below are the inter-disappearance times for all 6 observers. Each figure shows the results obtained in different conditions (row). The left column shows the distribution of the inter-disappearance times, the right column, the empirical characteristic function. The characteristic function for strictly periodic distributions would have a peak at the frequency corresponding to a period. The conditions are (from bottom to top) Troxler, a static mask, and a rotating mask with periods of 1, 2, 4, and 8 seconds. The distributions are not shown if there were fewer than 5 disappearance events.

**Figure S2 1.**
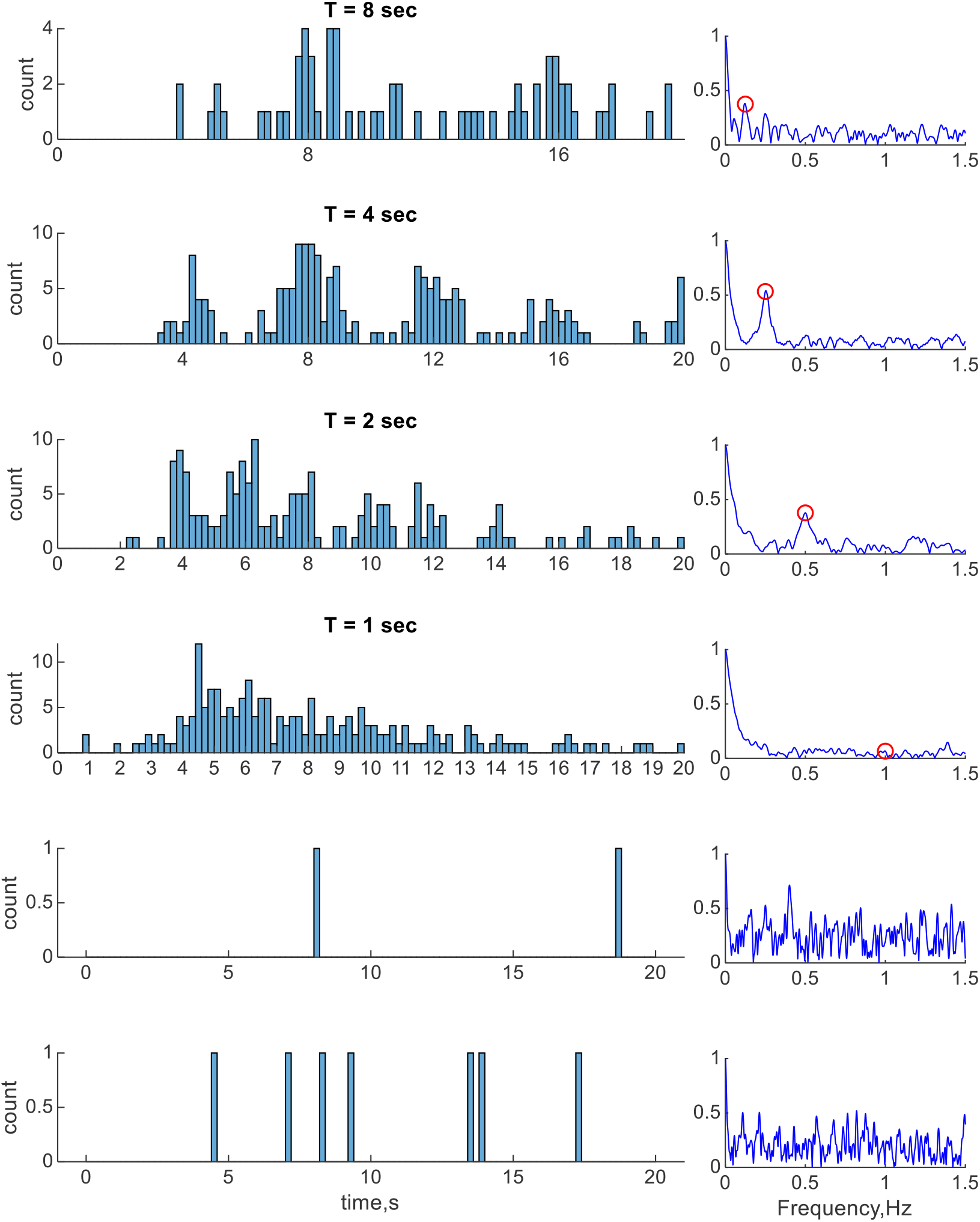
*Inter-disappearance times for observer O1. The bottom row shows the results for the static mask condition, and one row above it shows the results for the Troxler effect (no mask present)*.

**Figure S2 2.**
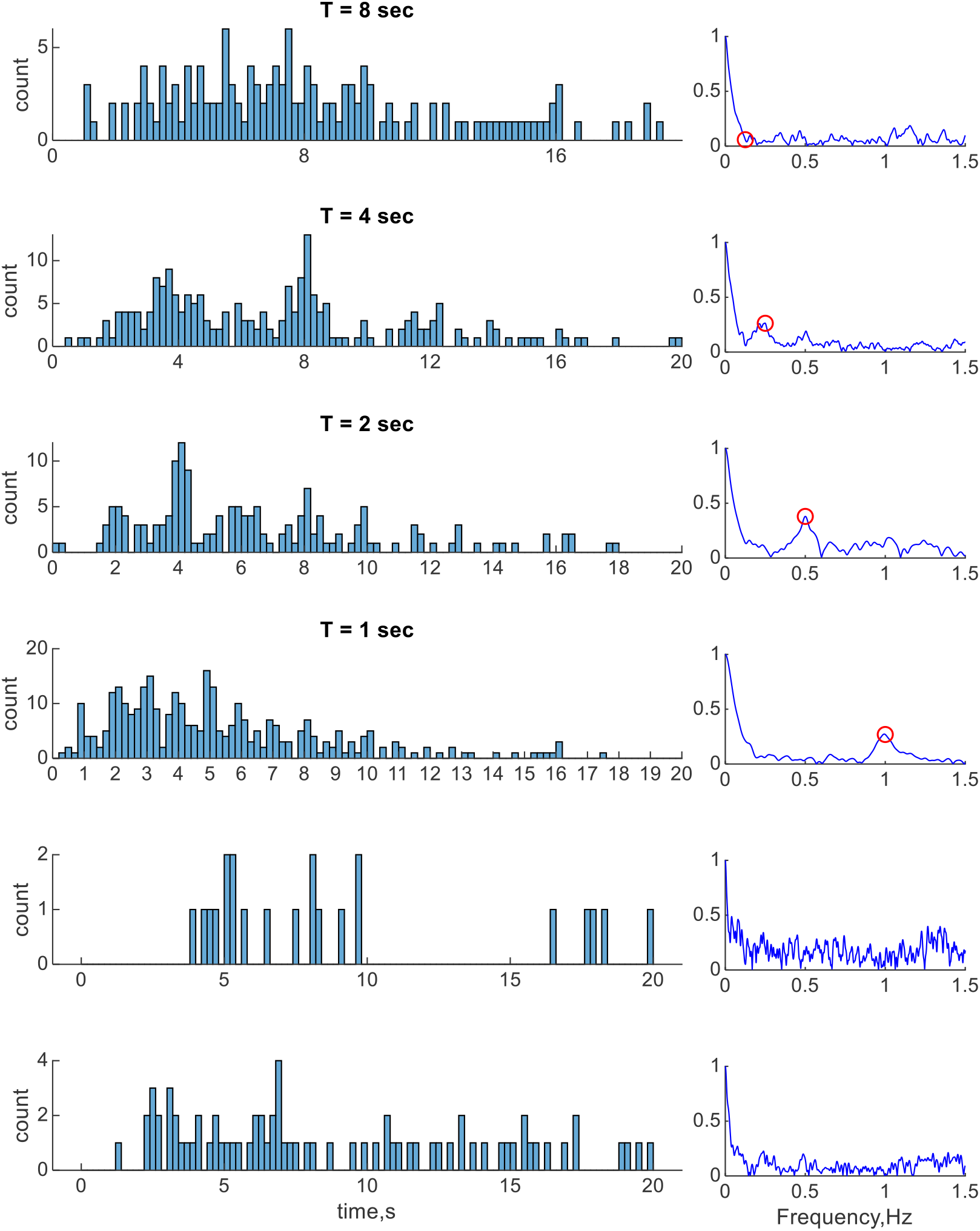
*Inter-disappearance times for observer O2. The bottom row shows the results for the static mask condition, and one row above it shows the results for the Troxler effect (no mask present)*.

**Figure S2 3.**
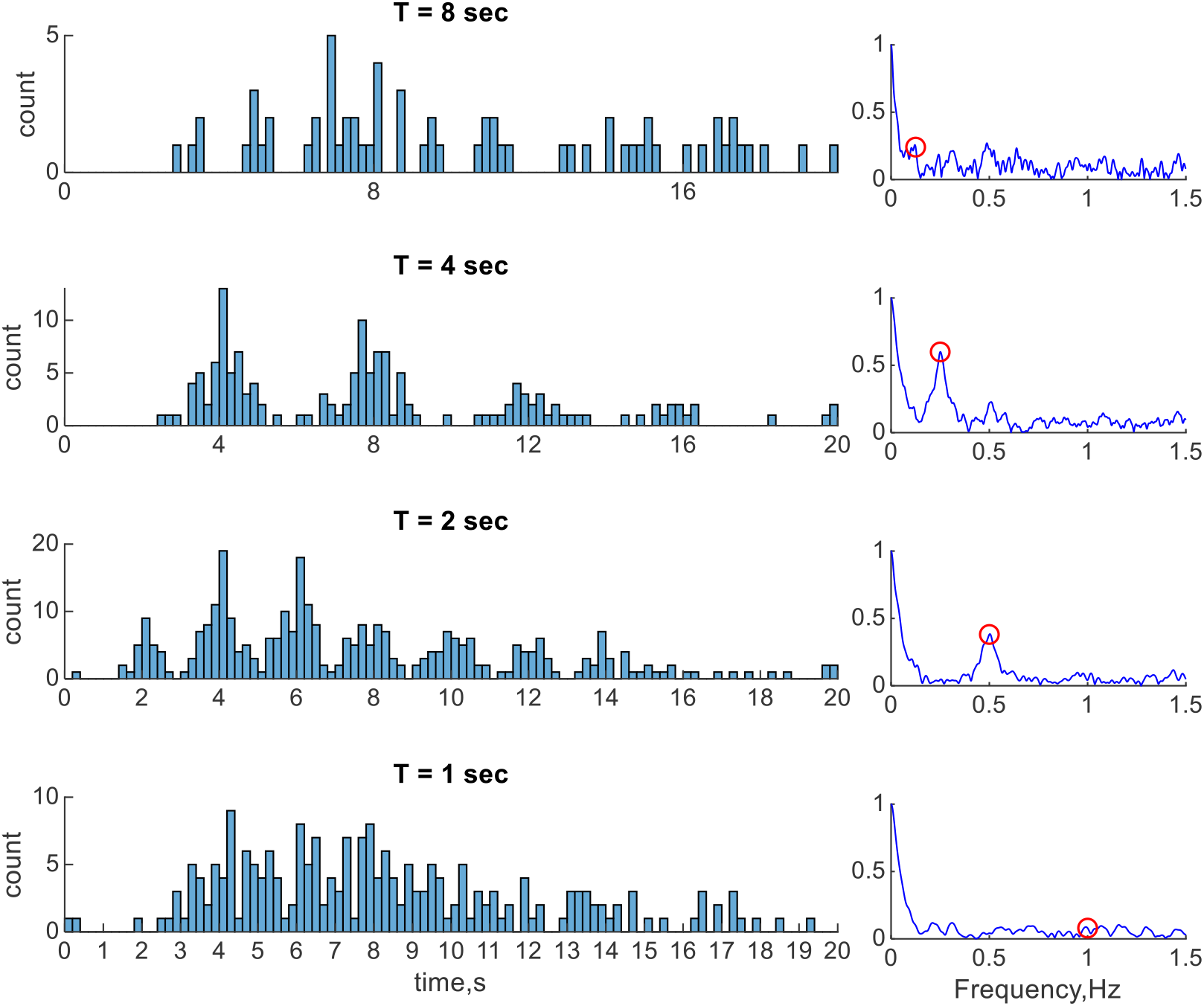
*Inter-disappearance times for observer O3*

**Figure S2 4.**
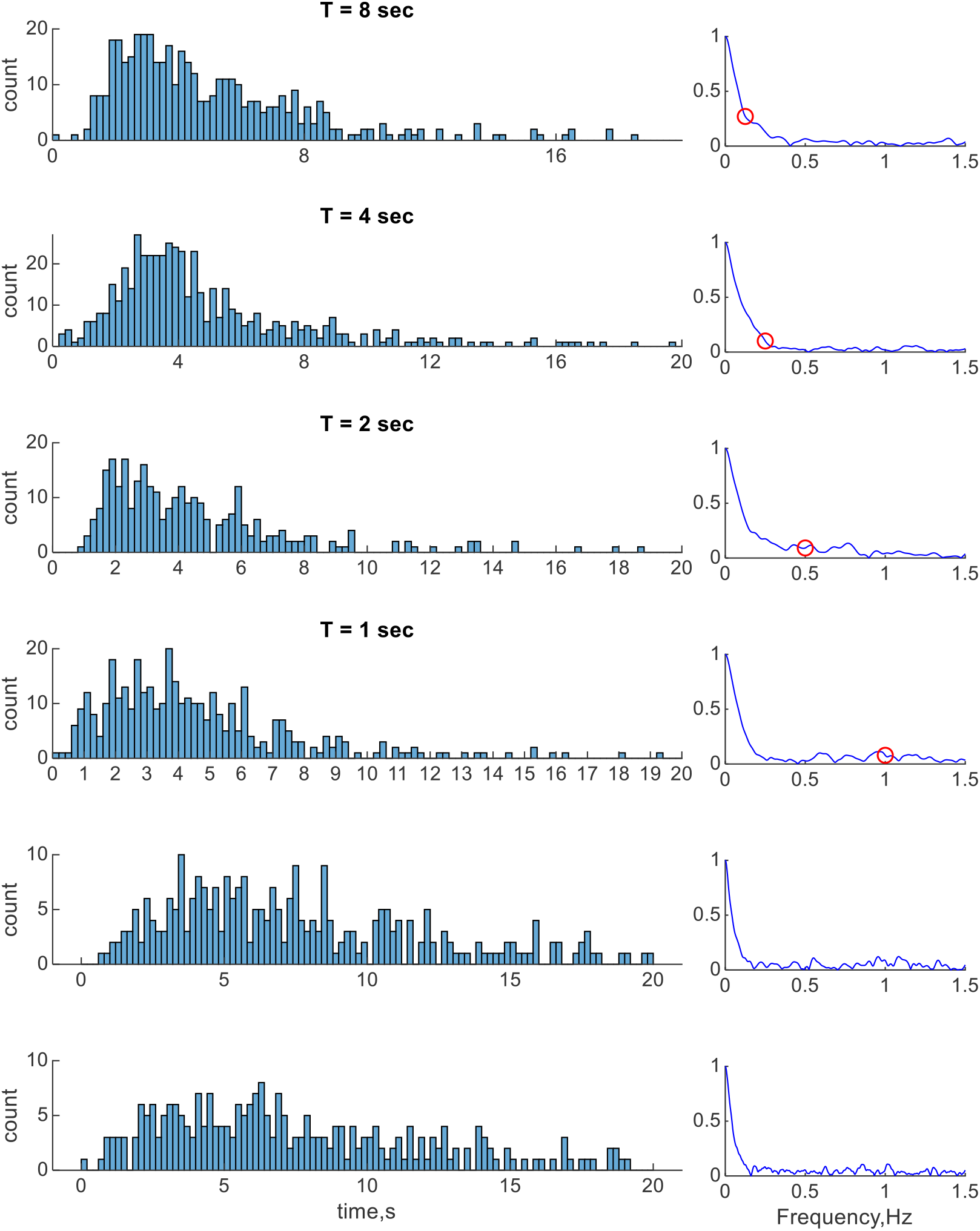
*Inter-disappearance times for observer O4. The bottom row shows the results for the static mask condition, and one row above it shows the results for the Troxler effect (no mask present)*.

**Figure S2 5.**
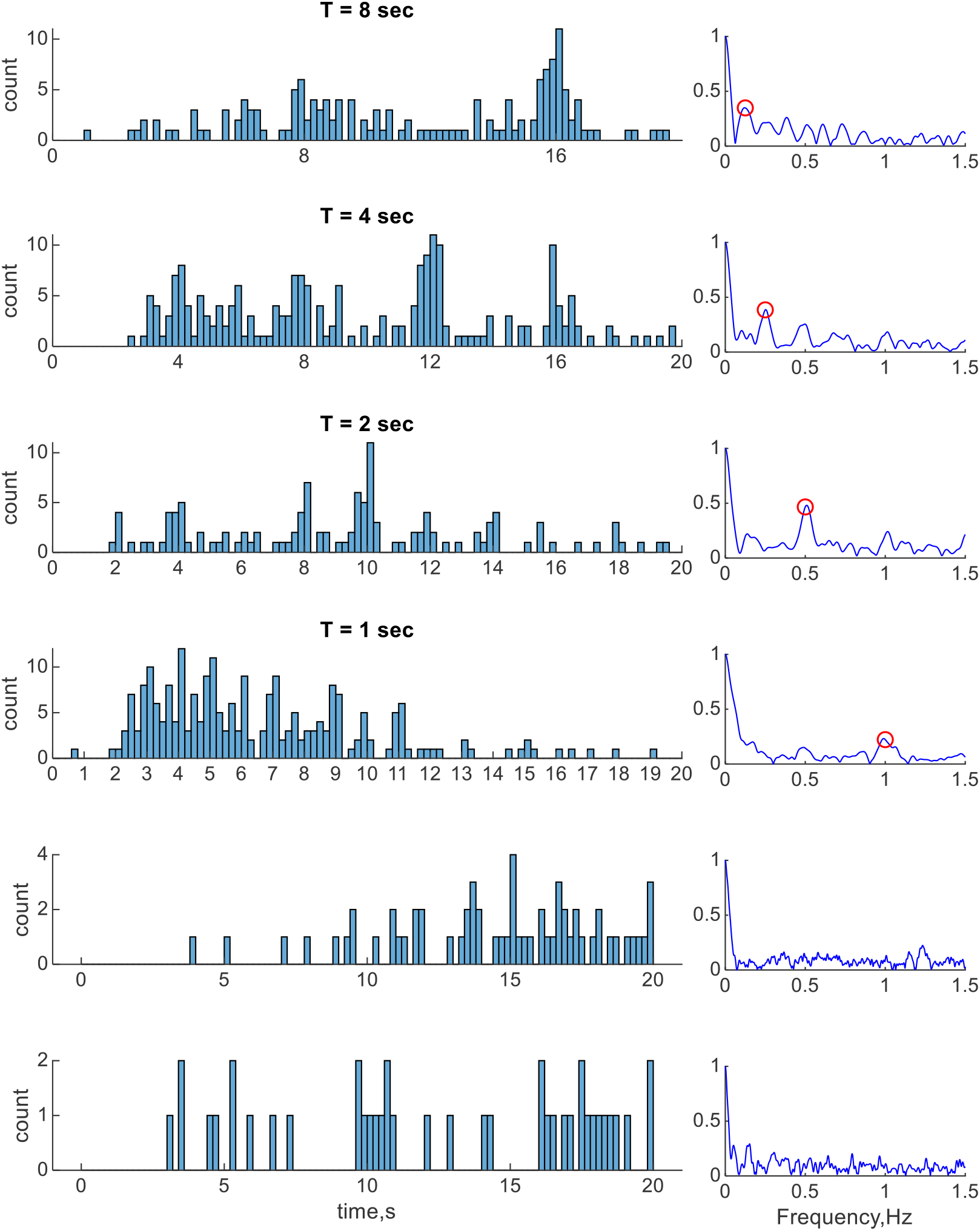
*Inter-disappearance times for observer O5. The bottom row shows the results for the static mask condition, and one row above it shows the results for the Troxler effect (no mask present)*.

**Figure S2 6.**
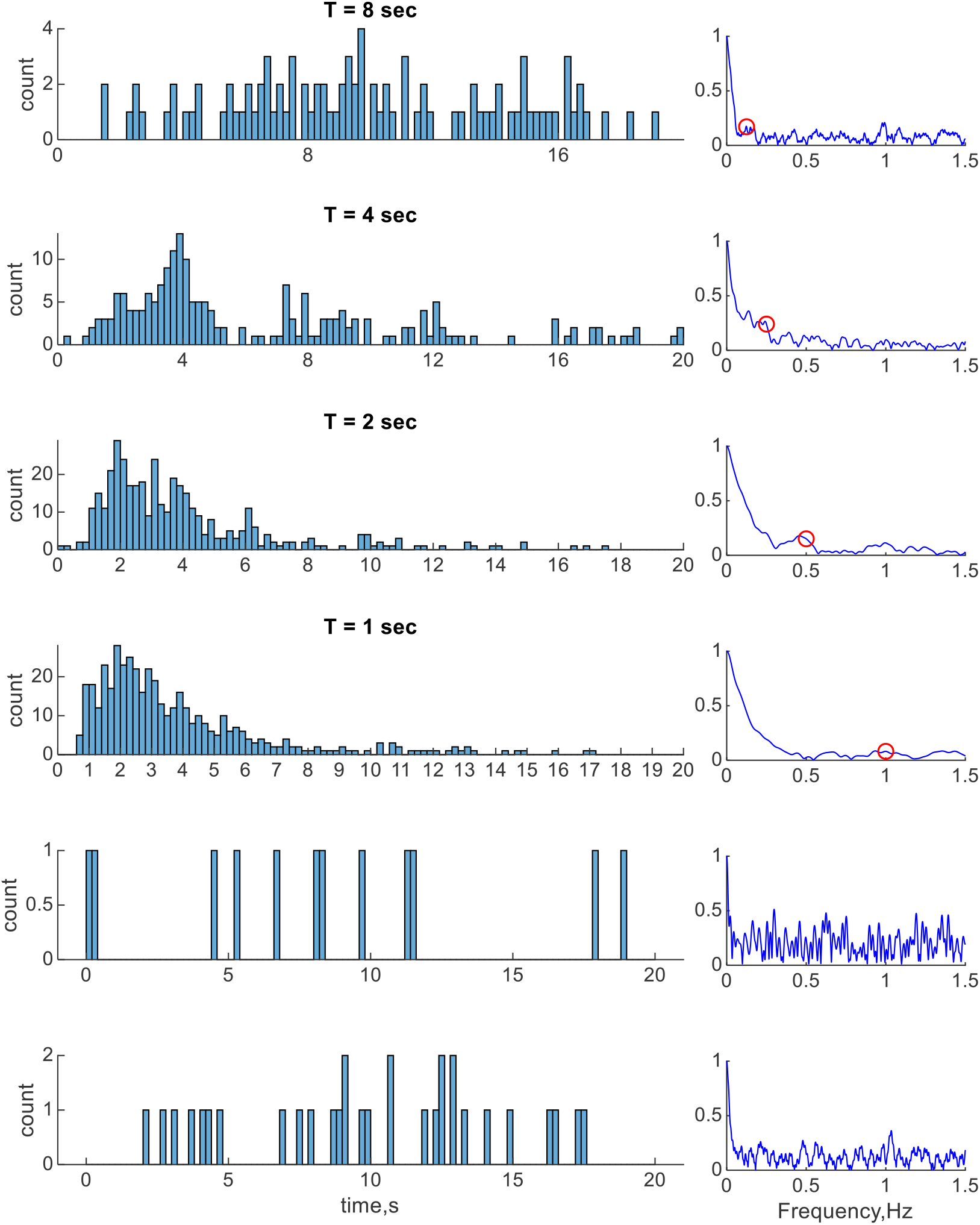
*Inter-disappearance times for observer O6. The bottom row shows the results for the static mask condition, and one row above it shows the results for the Troxler effect (no mask present)*.

### Distributions of visible and invisible periods for all observers

**Figure S3 1.**
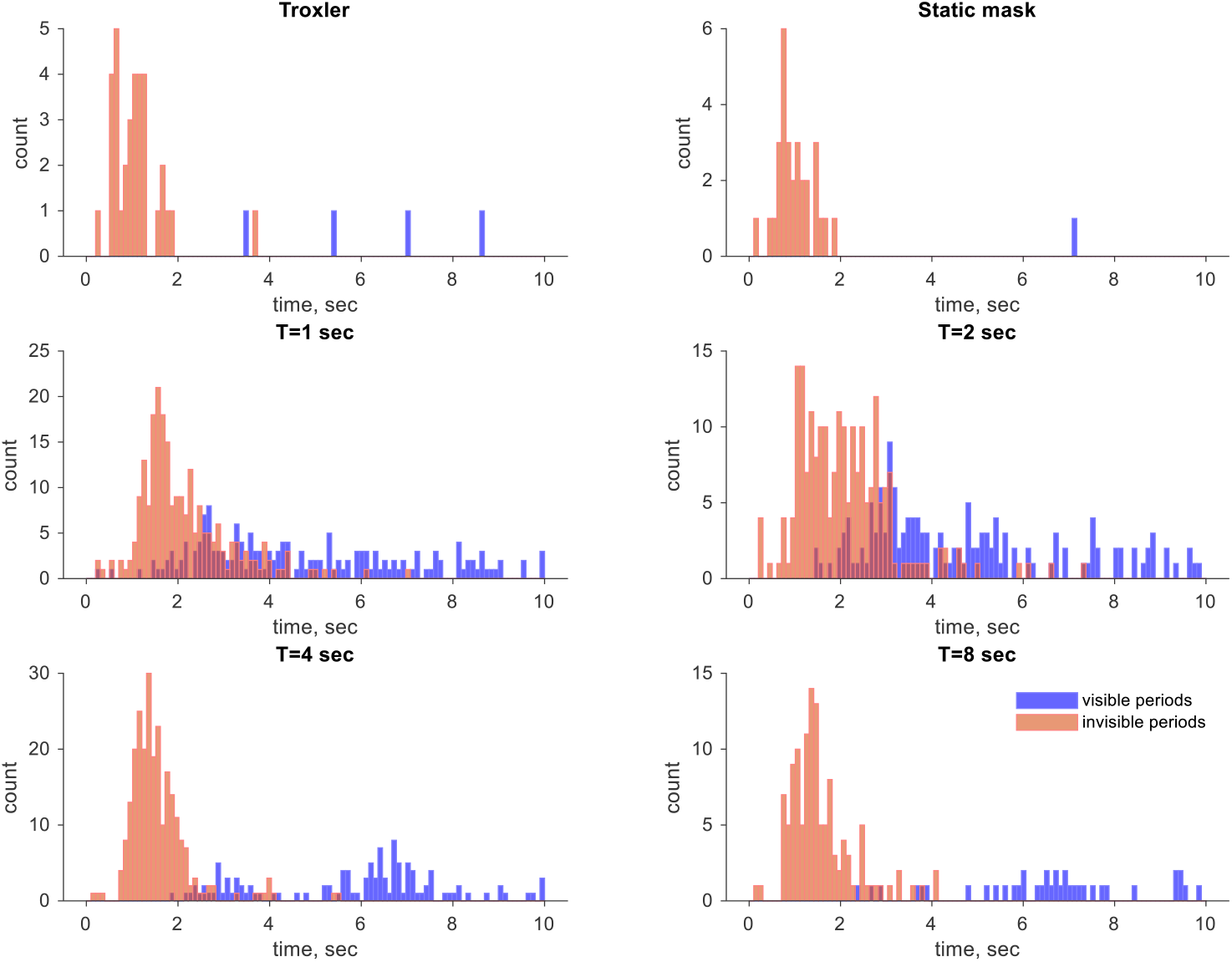
*Distributions of visible and invisible periods for observer O1*

**Figure S3 2.**
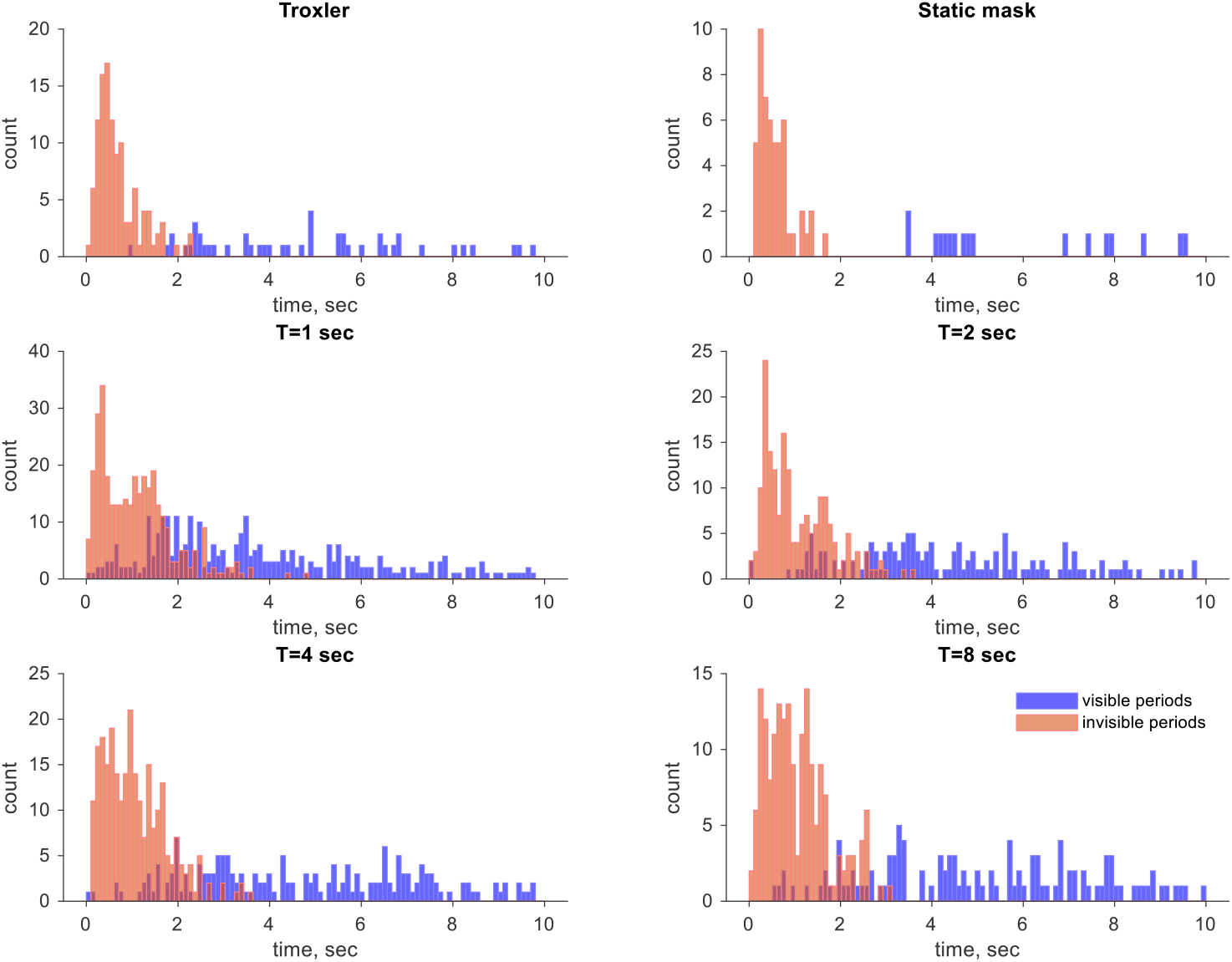
*Distributions of visible and invisible periods for observer O2*

**Figure S3 3.**
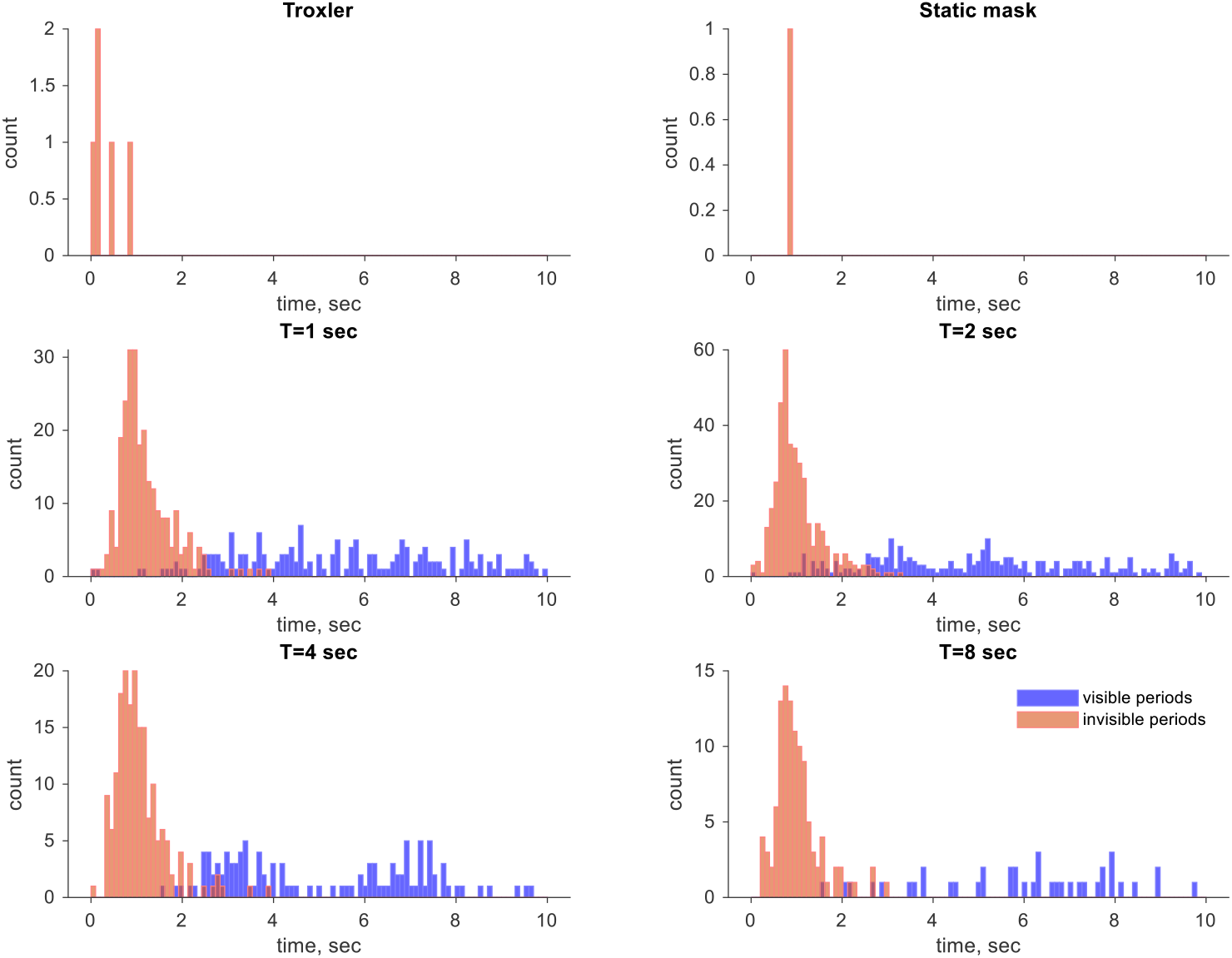
*Distributions of visible and invisible periods for observer O3*

**Figure S3 4.**
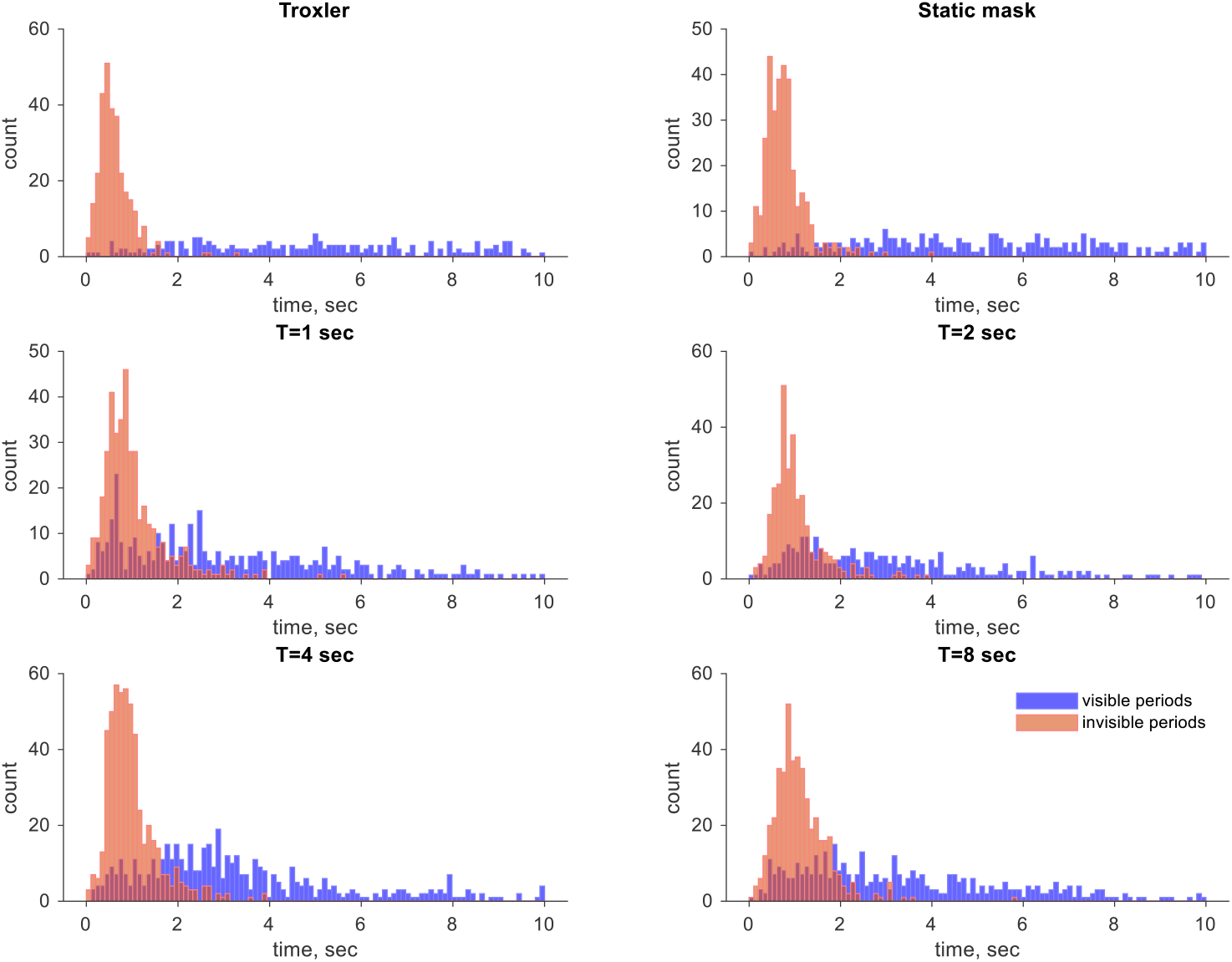
*Distributions of visible and invisible periods for observer O4*

**Figure S3 5.**
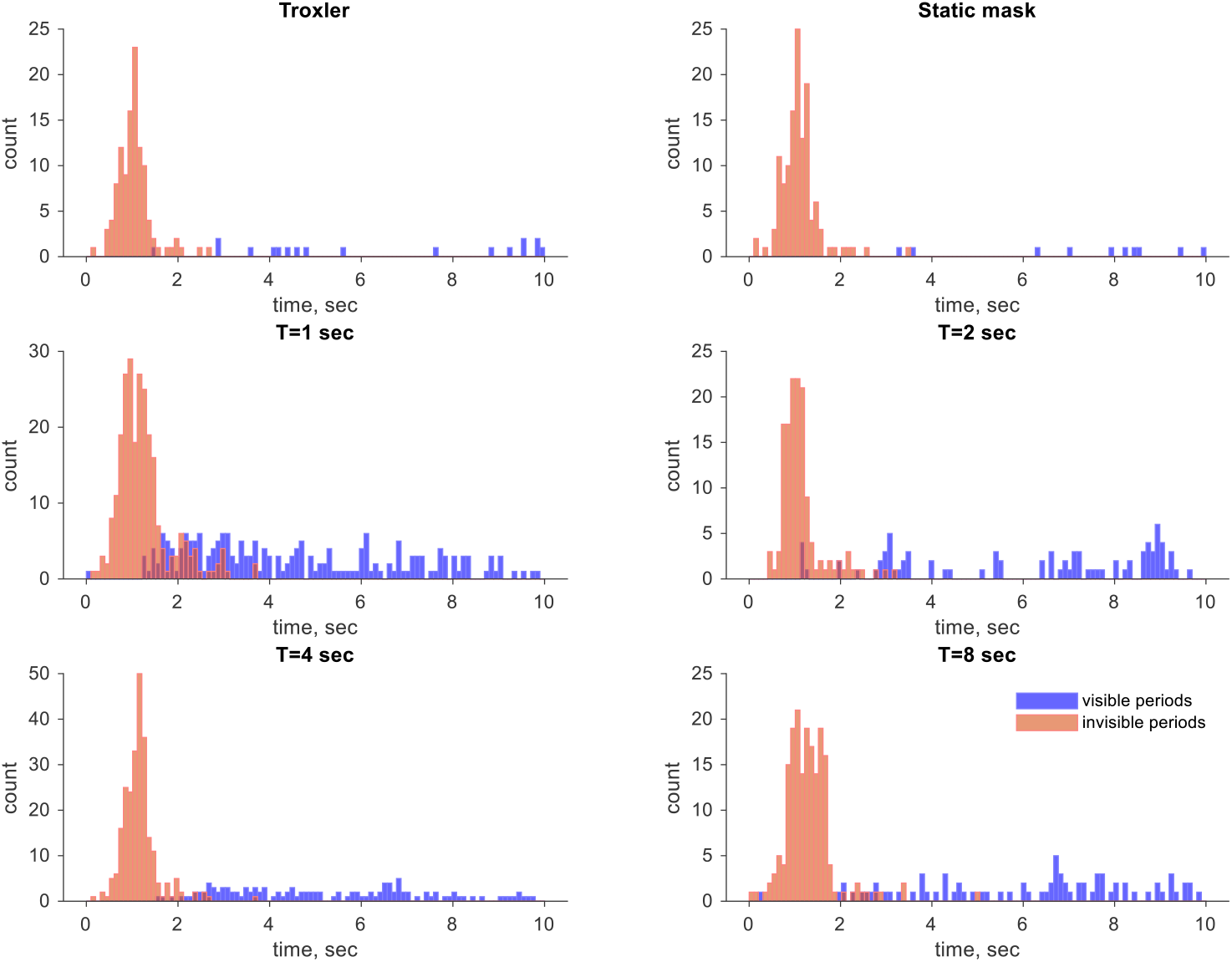
*Distributions of visible and invisible periods for observer O5*

**Figure S3 6.**
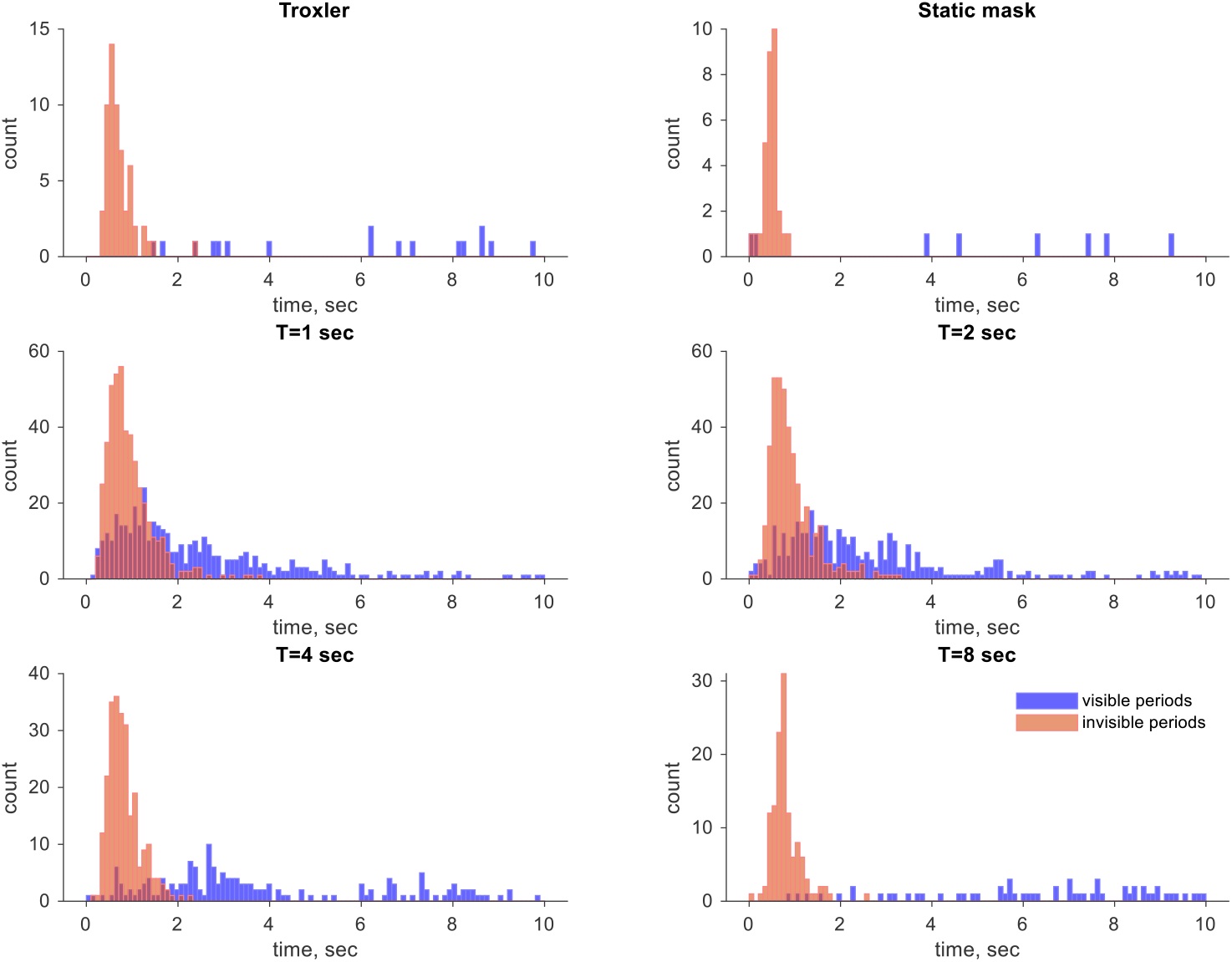
*Distributions of visible and invisible periods for observer O6*

### The distribution parameters in the exploratory experiments

There were many experiments under different experimental conditions. The table below presents the parameters of the Gaussian and the Gamma distributions fitted to the data. Only 7 p-values (out of 112) are below 0.05.

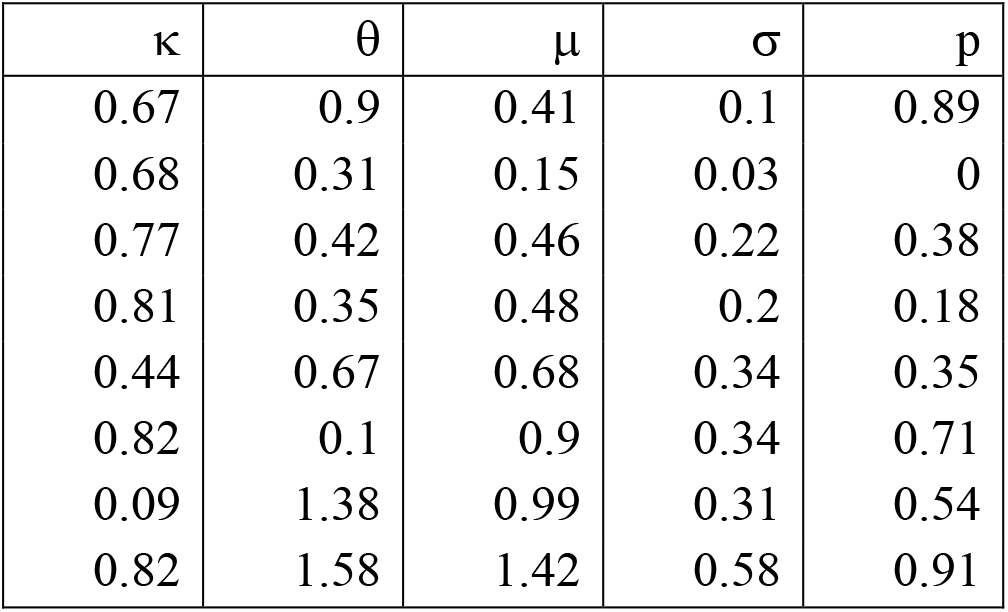
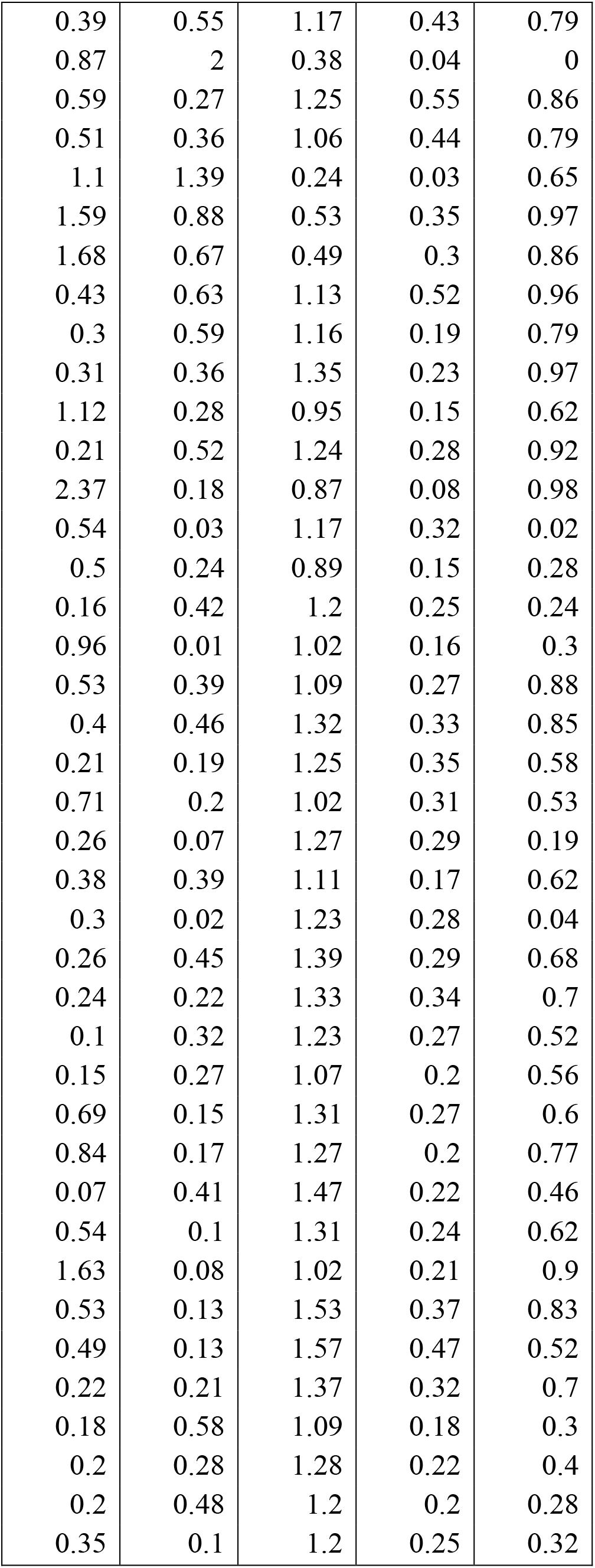
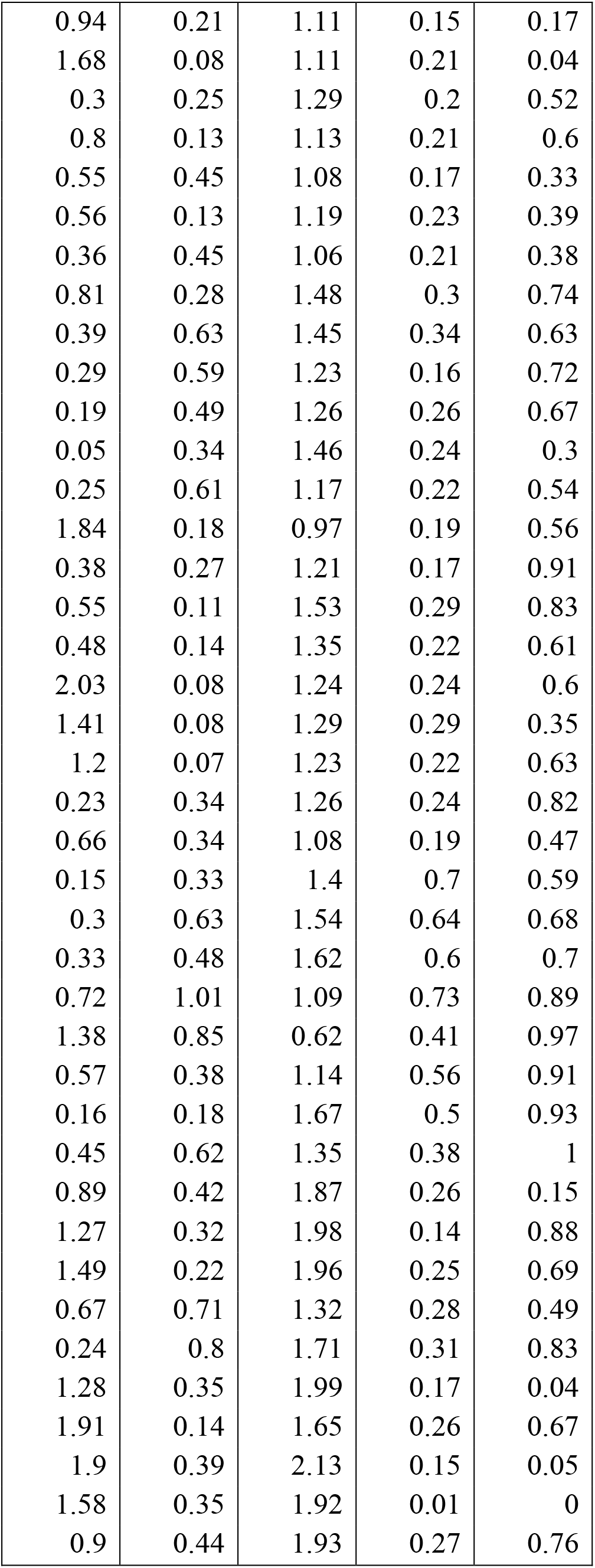
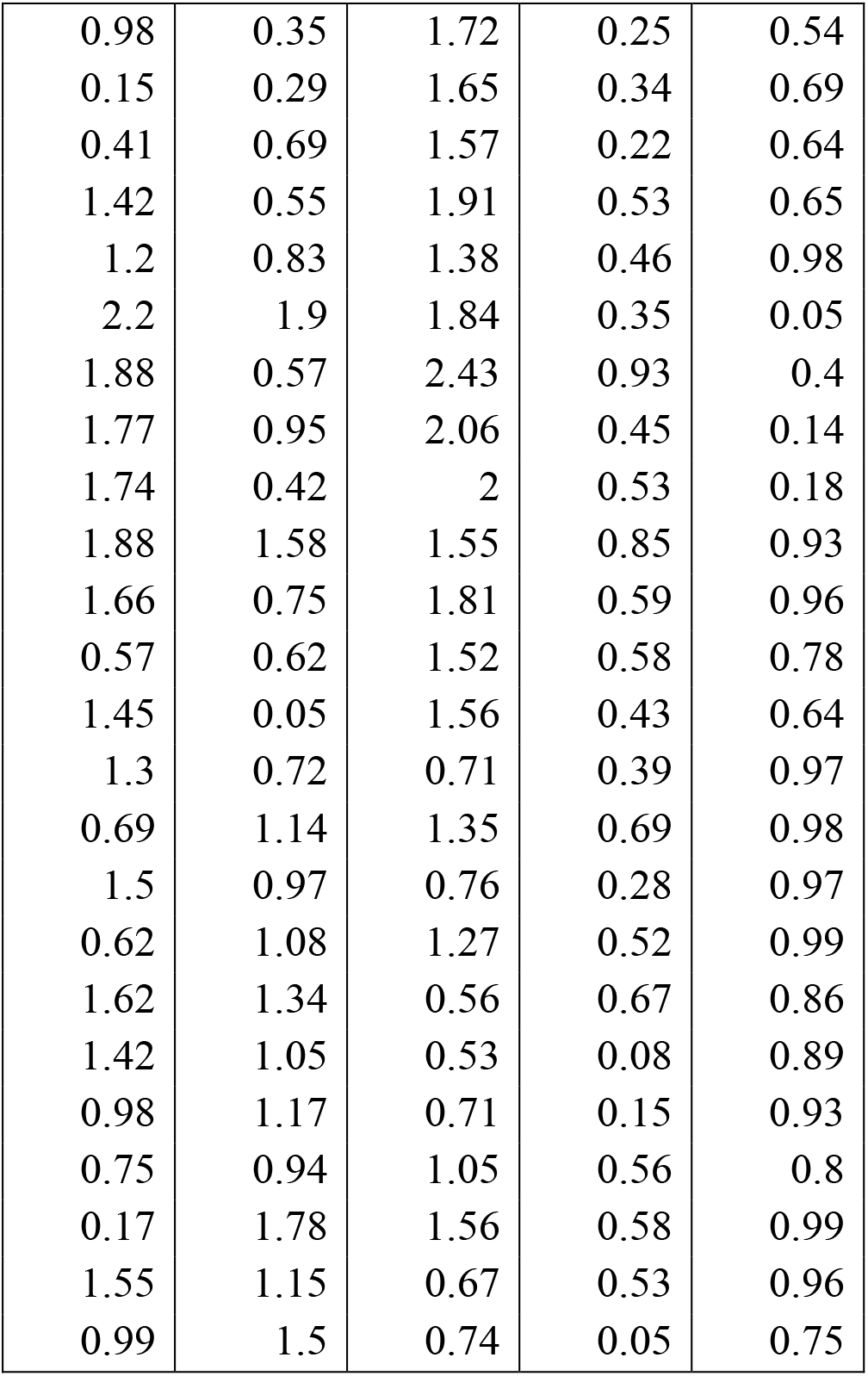

| $\kappa$ | $\theta$ | $\mu$ | $\sigma$ | p |
| --- | --- | --- | --- | --- |
| 0.67 | 0.9 | 0.41 | 0.1 | 0.89 |
| 0.68 | 0.31 | 0.15 | 0.03 | 0 |
| 0.77 | 0.42 | 0.46 | 0.22 | 0.38 |
| 0.81 | 0.35 | 0.48 | 0.2 | 0.18 |
| 0.44 | 0.67 | 0.68 | 0.34 | 0.35 |
| 0.82 | 0.1 | 0.9 | 0.34 | 0.71 |
| 0.09 | 1.38 | 0.99 | 0.31 | 0.54 |
| 0.82 | 1.58 | 1.42 | 0.58 | 0.91 |
| 0.39 | 0.55 | 1.17 | 0.43 | 0.79 |
| 0.87 | 2 | 0.38 | 0.04 | 0 |
| 0.59 | 0.27 | 1.25 | 0.55 | 0.86 |
| 0.51 | 0.36 | 1.06 | 0.44 | 0.79 |
| 1.1 | 1.39 | 0.24 | 0.03 | 0.65 |
| 1.59 | 0.88 | 0.53 | 0.35 | 0.97 |
| 1.68 | 0.67 | 0.49 | 0.3 | 0.86 |
| 0.43 | 0.63 | 1.13 | 0.52 | 0.96 |
| 0.3 | 0.59 | 1.16 | 0.19 | 0.79 |
| 0.31 | 0.36 | 1.35 | 0.23 | 0.97 |
| 1.12 | 0.28 | 0.95 | 0.15 | 0.62 |
| 0.21 | 0.52 | 1.24 | 0.28 | 0.92 |
| 2.37 | 0.18 | 0.87 | 0.08 | 0.98 |
| 0.54 | 0.03 | 1.17 | 0.32 | 0.02 |
| 0.5 | 0.24 | 0.89 | 0.15 | 0.28 |
| 0.16 | 0.42 | 1.2 | 0.25 | 0.24 |
| 0.96 | 0.01 | 1.02 | 0.16 | 0.3 |
| 0.53 | 0.39 | 1.09 | 0.27 | 0.88 |
| 0.4 | 0.46 | 1.32 | 0.33 | 0.85 |
| 0.21 | 0.19 | 1.25 | 0.35 | 0.58 |
| 0.71 | 0.2 | 1.02 | 0.31 | 0.53 |
| 0.26 | 0.07 | 1.27 | 0.29 | 0.19 |
| 0.38 | 0.39 | 1.11 | 0.17 | 0.62 |
| 0.3 | 0.02 | 1.23 | 0.28 | 0.04 |
| 0.26 | 0.45 | 1.39 | 0.29 | 0.68 |
| 0.24 | 0.22 | 1.33 | 0.34 | 0.7 |
| 0.1 | 0.32 | 1.23 | 0.27 | 0.52 |
| 0.15 | 0.27 | 1.07 | 0.2 | 0.56 |
| 0.69 | 0.15 | 1.31 | 0.27 | 0.6 |
| 0.84 | 0.17 | 1.27 | 0.2 | 0.77 |
| 0.07 | 0.41 | 1.47 | 0.22 | 0.46 |
| 0.54 | 0.1 | 1.31 | 0.24 | 0.62 |
| 1.63 | 0.08 | 1.02 | 0.21 | 0.9 |
| 0.53 | 0.13 | 1.53 | 0.37 | 0.83 |
| 0.49 | 0.13 | 1.57 | 0.47 | 0.52 |
| 0.22 | 0.21 | 1.37 | 0.32 | 0.7 |
| 0.18 | 0.58 | 1.09 | 0.18 | 0.3 |
| 0.2 | 0.28 | 1.28 | 0.22 | 0.4 |
| 0.2 | 0.48 | 1.2 | 0.2 | 0.28 |
| 0.35 | 0.1 | 1.2 | 0.25 | 0.32 |
| 0.94 | 0.21 | 1.11 | 0.15 | 0.17 |
| 1.68 | 0.08 | 1.11 | 0.21 | 0.04 |
| 0.3 | 0.25 | 1.29 | 0.2 | 0.52 |
| 0.8 | 0.13 | 1.13 | 0.21 | 0.6 |
| 0.55 | 0.45 | 1.08 | 0.17 | 0.33 |
| 0.56 | 0.13 | 1.19 | 0.23 | 0.39 |
| 0.36 | 0.45 | 1.06 | 0.21 | 0.38 |
| 0.81 | 0.28 | 1.48 | 0.3 | 0.74 |
| 0.39 | 0.63 | 1.45 | 0.34 | 0.63 |
| 0.29 | 0.59 | 1.23 | 0.16 | 0.72 |
| 0.19 | 0.49 | 1.26 | 0.26 | 0.67 |
| 0.05 | 0.34 | 1.46 | 0.24 | 0.3 |
| 0.25 | 0.61 | 1.17 | 0.22 | 0.54 |
| 1.84 | 0.18 | 0.97 | 0.19 | 0.56 |
| 0.38 | 0.27 | 1.21 | 0.17 | 0.91 |
| 0.55 | 0.11 | 1.53 | 0.29 | 0.83 |
| 0.48 | 0.14 | 1.35 | 0.22 | 0.61 |
| 2.03 | 0.08 | 1.24 | 0.24 | 0.6 |
| 1.41 | 0.08 | 1.29 | 0.29 | 0.35 |
| 1.2 | 0.07 | 1.23 | 0.22 | 0.63 |
| 0.23 | 0.34 | 1.26 | 0.24 | 0.82 |
| 0.66 | 0.34 | 1.08 | 0.19 | 0.47 |
| 0.15 | 0.33 | 1.4 | 0.7 | 0.59 |
| 0.3 | 0.63 | 1.54 | 0.64 | 0.68 |
| 0.33 | 0.48 | 1.62 | 0.6 | 0.7 |
| 0.72 | 1.01 | 1.09 | 0.73 | 0.89 |
| 1.38 | 0.85 | 0.62 | 0.41 | 0.97 |
| 0.57 | 0.38 | 1.14 | 0.56 | 0.91 |
| 0.16 | 0.18 | 1.67 | 0.5 | 0.93 |
| 0.45 | 0.62 | 1.35 | 0.38 | 1 |
| 0.89 | 0.42 | 1.87 | 0.26 | 0.15 |
| 1.27 | 0.32 | 1.98 | 0.14 | 0.88 |
| 1.49 | 0.22 | 1.96 | 0.25 | 0.69 |
| 0.67 | 0.71 | 1.32 | 0.28 | 0.49 |
| 0.24 | 0.8 | 1.71 | 0.31 | 0.83 |
| 1.28 | 0.35 | 1.99 | 0.17 | 0.04 |
| 1.91 | 0.14 | 1.65 | 0.26 | 0.67 |
| 1.9 | 0.39 | 2.13 | 0.15 | 0.05 |
| 1.58 | 0.35 | 1.92 | 0.01 | 0 |
| 0.9 | 0.44 | 1.93 | 0.27 | 0.76 |
| 0.98 | 0.35 | 1.72 | 0.25 | 0.54 |
| 0.15 | 0.29 | 1.65 | 0.34 | 0.69 |
| 0.41 | 0.69 | 1.57 | 0.22 | 0.64 |
| 1.42 | 0.55 | 1.91 | 0.53 | 0.65 |
| 1.2 | 0.83 | 1.38 | 0.46 | 0.98 |
| 2.2 | 1.9 | 1.84 | 0.35 | 0.05 |
| 1.88 | 0.57 | 2.43 | 0.93 | 0.4 |
| 1.77 | 0.95 | 2.06 | 0.45 | 0.14 |
| 1.74 | 0.42 | 2 | 0.53 | 0.18 |
| 1.88 | 1.58 | 1.55 | 0.85 | 0.93 |
| 1.66 | 0.75 | 1.81 | 0.59 | 0.96 |
| 0.57 | 0.62 | 1.52 | 0.58 | 0.78 |
| 1.45 | 0.05 | 1.56 | 0.43 | 0.64 |
| 1.3 | 0.72 | 0.71 | 0.39 | 0.97 |
| 0.69 | 1.14 | 1.35 | 0.69 | 0.98 |
| 1.5 | 0.97 | 0.76 | 0.28 | 0.97 |
| 0.62 | 1.08 | 1.27 | 0.52 | 0.99 |
| 1.62 | 1.34 | 0.56 | 0.67 | 0.86 |
| 1.42 | 1.05 | 0.53 | 0.08 | 0.89 |
| 0.98 | 1.17 | 0.71 | 0.15 | 0.93 |
| 0.75 | 0.94 | 1.05 | 0.56 | 0.8 |
| 0.17 | 1.78 | 1.56 | 0.58 | 0.99 |
| 1.55 | 1.15 | 0.67 | 0.53 | 0.96 |
| 0.99 | 1.5 | 0.74 | 0.05 | 0.75 |

**Figure S4.**
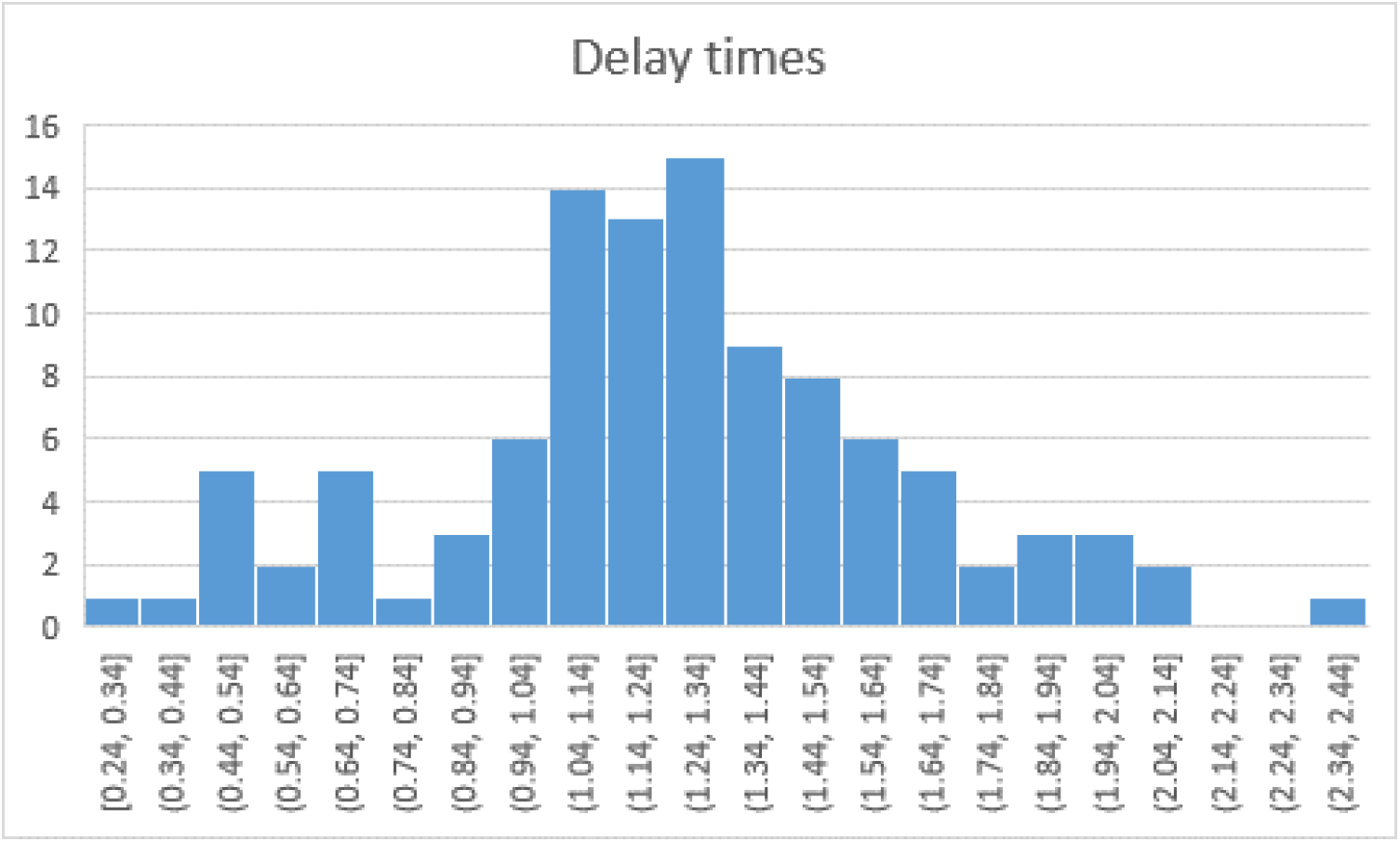
Distribution of the fixed delays. There is a minimal gap of .24 sec (excluding bad fits) and a typical value is of the order of one second. Presumably, this is the time it takes for the system to return to a resting state once excited.

### FitzHugh-Nagumo. Basics

We assume that stable perceptions correspond to stable fixed points in the dynamics of the two-dimensional decision model. We also assume that both variables are noisy, reflecting background neuronal activity not related to the stimulus in question. The illustrative example of such a system is the FitzHugh–Nagumo class of models (FHN, (FitzHugh, 1961; Nagumo et al., 1962; Sherwood, 2014)), which can be described (without noise) by following a set of differential equations

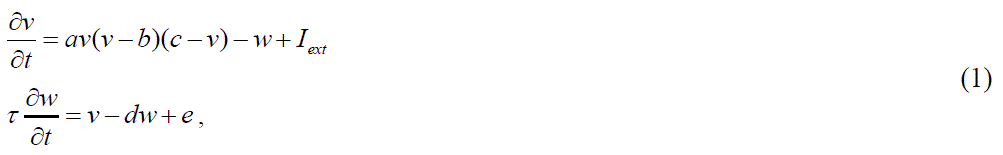

where *v* is a decision variable, *w* is a recovery variable, *I_ext_* is an external input, *τ* is a relative time constant of a recovery variable, and *a*, *b*, *c*, *d* and *e* are some constants defining the behavior of a dynamical system. This dynamical system has a few prototypical behaviors that can be understood by analyzing its phase diagram.

The basic analysis of dynamical systems is carried out by examining conditions where the right side of the differential equation is equal to zero. In this situation, the corresponding variable does not change at a particular moment. All the points where such a condition is satisfied for a single differential equation are called a nullcline. Since there are two equations in System (1), there are two nullclines (Figure S 1). The point where all (two in this case) nulclines intersect (if such a point exists) is a fixed point of a dynamical system, i.e., in the absence of noise, the dynamical system will stay at this point forever if the dynamics start at this point. Nevertheless, in the presence of noise, there are two types of system behavior: a stable fixed point – when any small perturbation of the system due to noise or due to a force will be counteracted by the dynamical system and the system returns to the fixed point or an unstable fixed point – when small perturbations are amplified by the dynamical system and the system will move away from the fixed point – this is referred to as an unstable fixed point. The FHN system may have 1, 2, or 3 fixed points, depending on the parameters.

One can see that the nullcline for the first differential equation is cubic, which in general has one local minimum and one local maximum (the blue curves in Figure S 1). For stability reasons, it is important that on the left side the curve goes to plus infinity, whereas on the right side it goes to minus infinity. The second nullcline is a straight line with positive slope (the red line in Figure S 1). The behavior of the system depends on where the nullclines intersect. If the linear nullcline intersects the cubic nullcline on the left side of the local minimum (Figure S 1a), this intersection point will be a stable fixed point, i.e., the dynamical system will approach this point. In contrast, if the linear nullcline intersects the cubic nullcline in between the local minimum and maximum (Figure S 1b), the fixed point is unstable, and the system (in the presence of noise) will move away from this point, even if visited. This FHN regime leads to oscillatory behavior. The third interesting possibility is when the linear nullcline intersects the cubic nullcline before the minimum, between the minimum and maximum, and after the maximum (Figure S 1c). Here, the system has two stable fixed points. Therefore, in general we have the following cases: No stable fixed point (the linear nullcline intersects the cubic nullcline only between the local minimum and maximum) – the system undergoes oscillations. If there is one stable fixed point on the left side of the minimum or on the right side of the maximum (the system will end up near those points in the presence of small noise). If there are two stable fixed points on the left side of the local minimum and on the right side of the local maximum, the system will end up in one of the minima, depending on the initial conditions. In the presence of noise, FHN with parameters leading to 2 stable fixed points may be used to describe bistable phenomena, whereas FHN with parameters leading to a single fixed point may be used to describe the MIB.

**Figure S1.**
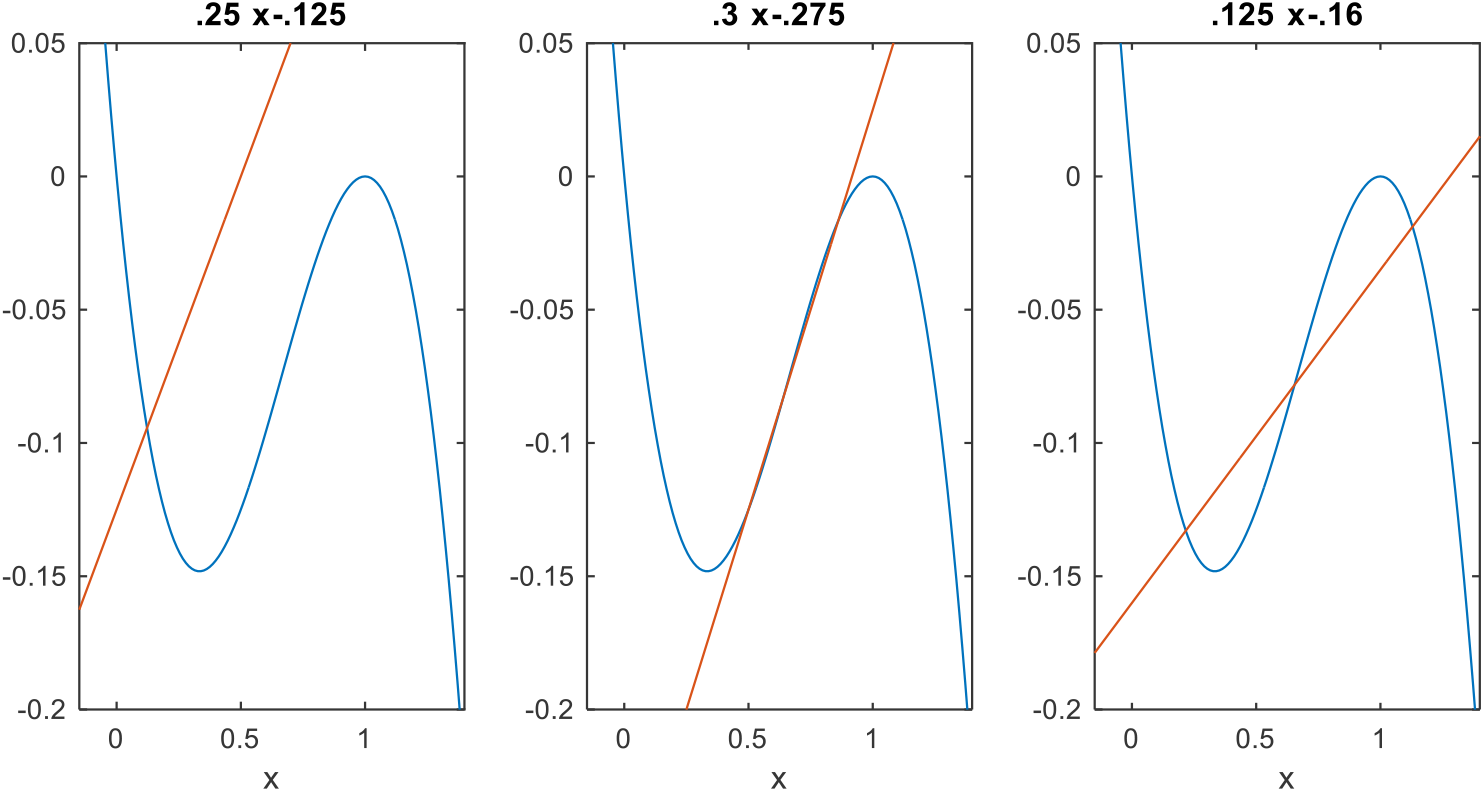
*Nullclines for 3 different regimes in the FitzHugh-Nagumo model*.

To describe MIB as an excitable system, one needs to make an additional assumption that a threshold exists on the fast variable V, so that when the system is on the right side of the threshold the target is perceived as invisible, and when the system is on the left side of the threshold the target is perceived as visible (see Figure S2). The mask can be considered to contribute to external input, which formally moves a cubic in a vertical direction, whereas the linear nullcline stays at the same place. When a cubic nullcline is driven up, the fixed point moves toward a bifurcation point (where the linear nullcline intersects the cubic in a local minimum), after which there is no stable fixed point. The interpretation would be that if there is a strong enough external input, the system will be excited by it and start a large trajectory. In the presence of noise, the closer the fixed point is to the bifurcation, the easier it is for the noise to excite the system. That is what probably happens when the mask bars approach a target – the fixed point is moved closer to the bifurcation point. Since invisibility is not induced every time the mask bar crosses the target, the fixed point does not cross the bifurcation point; moreover, the noise is relatively small in order to induce transition. However, since the invisibility periods are phase locked to the mask, the fixed point is probably moved substantially.

**Figure S2.**
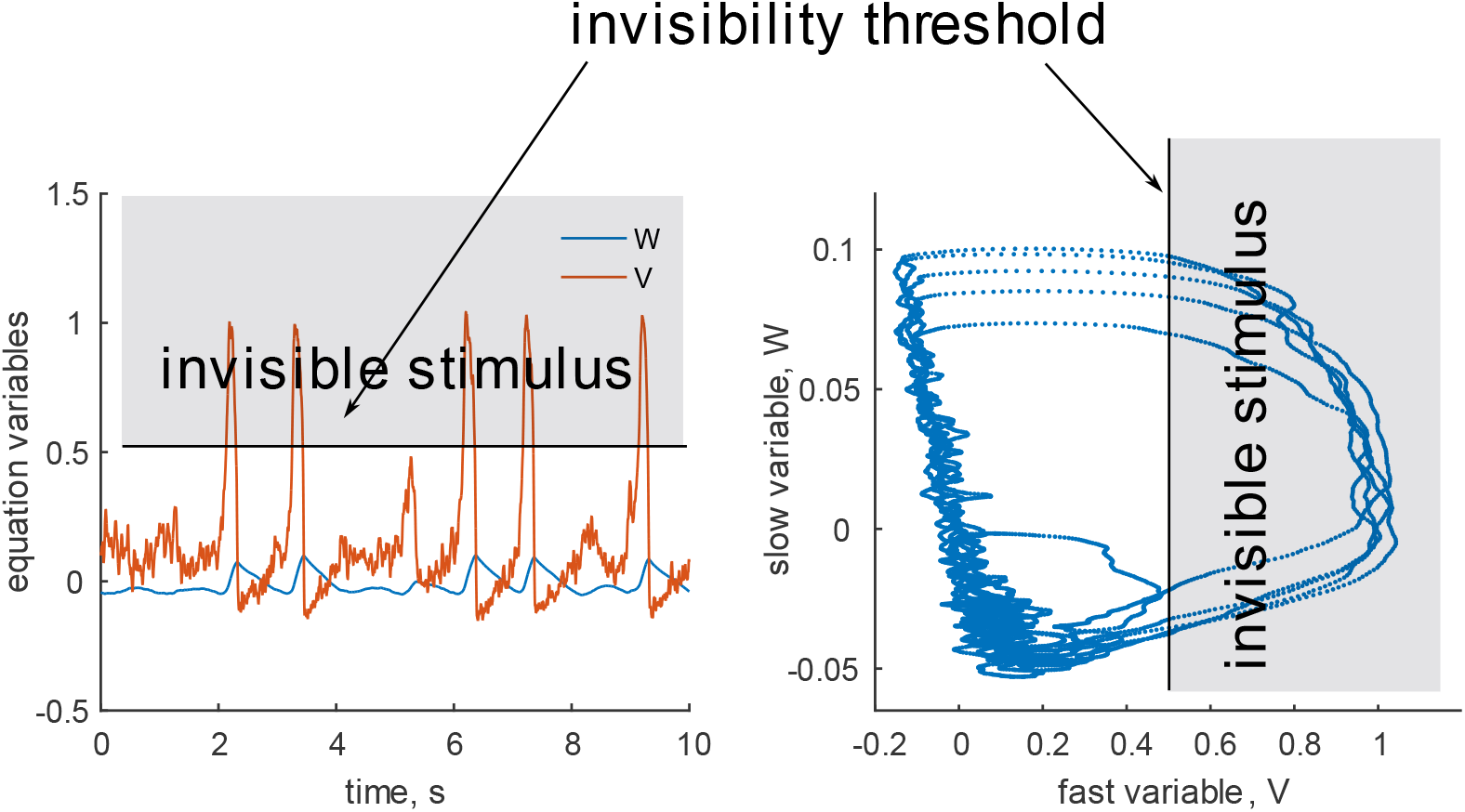
Illustration of the possible dynamics of the FitzHugh-Nagumo model describing the Troxler effect. The left plot shows the temporal dynamics of the fast and slow variables, where we assume that the fast variable is responsible for the perceptual visibility of the target. When the fast variable exceeds the threshold denoted by the black line and enters the gray area, the target is perceived as invisible, whereas when it returns to the unshaded region, the target is perceived as visible. The right plot shows the phase portrait of the dynamics showing the points (V(t), W(t)) along the trajectory. The shaded area represents part of the phase diagram where the target is invisible. It can be seen that most of the time the system is near the fixation point, with noise inducing some large trajectories.

### Stochastic resonance in a bistable FitzHugh-Nagumo system

In the following sections we use extended FitzHugh-Nagumo models with slightly different parameterization.

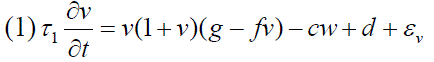

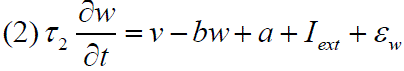

where *ε_v_* and *ε_w_* are Gaussian white noise with intensities parameterized by the same letters.

External stimulation is parameterized by

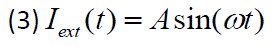

Simulation is performed on GPU using the Euler method with a time step of 10 ms. We will refer to this system as FHN.

In this section we will illustrate the effect of Stochastic resonance in FHN in a bistable regime. Simulation parameters are as follows:

|  |  |
| --- | --- |
| a | 0 |
| b | -0.1685 |
| c | -0.1 |
| d | 0 |
| f | 1/3 |
| g | 0 |
| $\tau_1$ | 0.25 |
| $\tau_2$ | 10.2 |
| A | 0.0001512131525 |
| $\omega$ | $2\pi \cdot 0.1218251826251$ |
| $\varepsilon_v$ | 0 |
| $\varepsilon_w$ | 0-1 (for dt 10 ms) |

All parameters except *ε _w_* were fixed, and *ε _w_* was varied between 0 and 1 in 32 steps. The top graph in Figure S3 represents SNR for different noise amplitudes. One can observe that there is an optimal noise level for which SNR is maximal. This effect is known as stochastic resonance (SR). To illustrate the mechanism underlying SR, we present a phase diagram and trajectory for 3 different levels of noise: top row – below optimal; middle row – close to optimal; bottom row –above optimal. The right column in Figure S3 shows traces for *v* and *w* over time. One can observe that for very low noise levels no transitions occur, leading to a low SNR; for high levels of noise, transitions occur at random times, whereas for near optimal noise levels, switches between states appear more regularly. The phase diagrams in the left column (plot of *v* vs *w*) and nullclines show this expected behavior: for low noise levels, the trajectory remains around a fixed point – the crossing of 2 nullclines near the minimum of a cubic nullcline. For high noise levels, there is a large diversity of trajectories, whereas near the optimal noise level the trajectories follow a very prototypical trajectory. However, once the system is driven outside of the basin of attraction, it moves along a horizontal line in phase space due to the large separation of time; then it follows a cubic nullcline and stays for some time in another stable state until noise drives the system out of basin of attraction (observe noisy horizontal lines in the trajectory for the *v* variable after positive or negative peaks).

**Figure S3.**
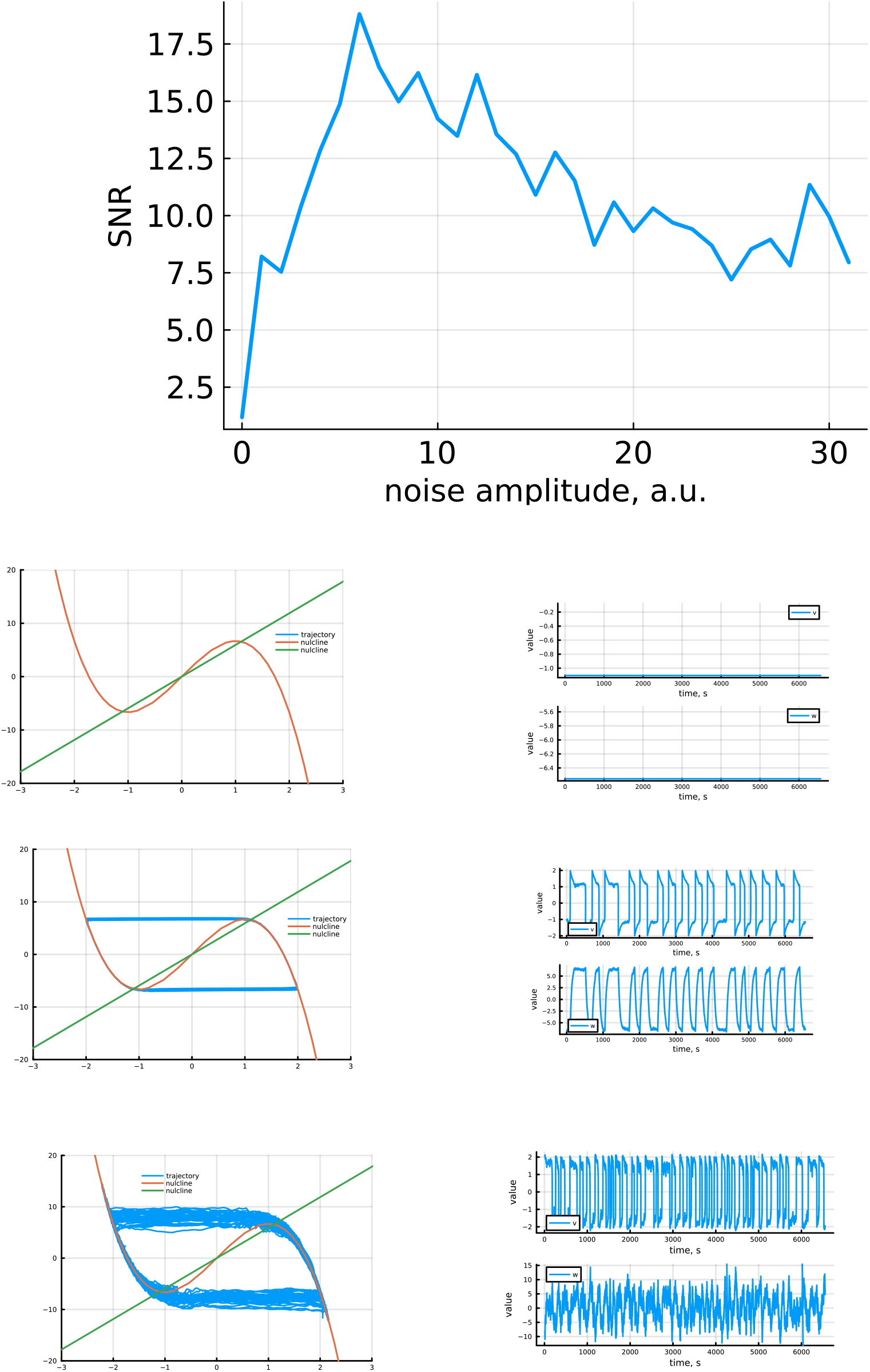

### Stochastic resonance in an excitable FitzHugh-Nagumo system

All simulations were performed similarly to the previous section, except the parameters of the model, which were mostly adapted from (Volkov, Ullner, Zaikin, & Kurths, 2003)):

|  |  |
| --- | --- |
| a | 1.01 |
| b | -0.001 |
| c | -0.1 |
| d | 0 |
| f | 1/3 |
| g | 0 |
| $\tau_1$ | 0.5 |
| $\tau_2$ | 5.0 |
| A | 0.0002 |
| $\omega$ | $2 \pi 0.31415927$ |
| $\varepsilon_v$ | 0 |
| $\varepsilon_w$ | 0-0.6 (for dt 10 ms) |

All parameters except *ε _w_* were fixed, and *ε _w_* was varied between 0 and 0.6 in 32 steps.

**Figure S4.**
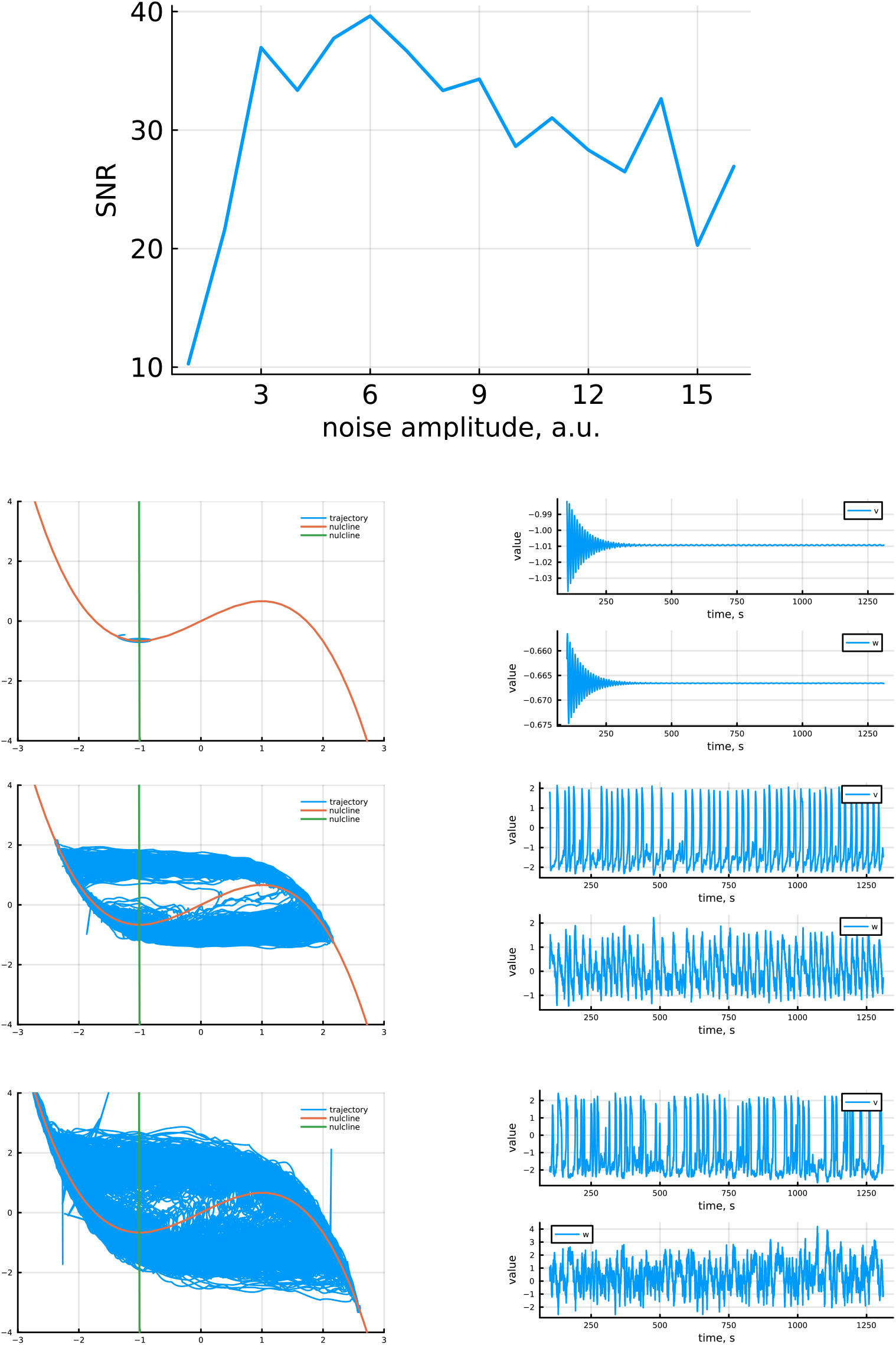

The top graph in Figure S4 represents SNR for different noise amplitudes. One can see that there is an optimal noise level for which SNR is maximal; therefore, SR is present in excitable systems. To illustrate the mechanism underlying the SR, we present phase diagrams and trajectories for three different noise levels: top row – below optimal; middle row – close to optimal; bottom row – above optimal. The right column depicts traces for *v* and *w* over time. One can observe that no transitions occur for the very low noise level; for the high noise level, transitions occur at random times, whereas for a near optimal noise level, switches between states appear more regularly. Phase diagrams in the left column (the plot of *v* vs *w*) and nullclines show the expected behavior: for the low noise level the trajectory stays around the fixed point – crossing 2 nullclines near the minimum of the cubic nullcline. For high noise levels, there is a large diversity of trajectories, whereas near the optimal noise level the trajectories follow a very prototypical trajectory; once the system is driven outside the basin of attraction, it moves along a horizontal line in phase space due to the large separation of time; then it follows the cubic nullcline. In contrast to bistable systems, the trajectories in the optimal case are noisier and the time spent in the excited state mainly depends on the internal dynamics with no stop for positive values, whereas in a bistable system there was a fixed point.

### Frequency dependence of stochastic resonance in an excitable FitzHugh-Nagumo system

A detailed analysis is provided in (Longtin & Chialvo, 1998; Volkov et al., 2003). Here, in Figures S5 and S6, we just illustrate the non-monotonic dependence of SNR on frequency for the same system as in the previous section. In the simulations we use the same parameters, except we fix *ε_w_* to 0.015, A to 0.01, and vary *ω*.

**Figure S5.**
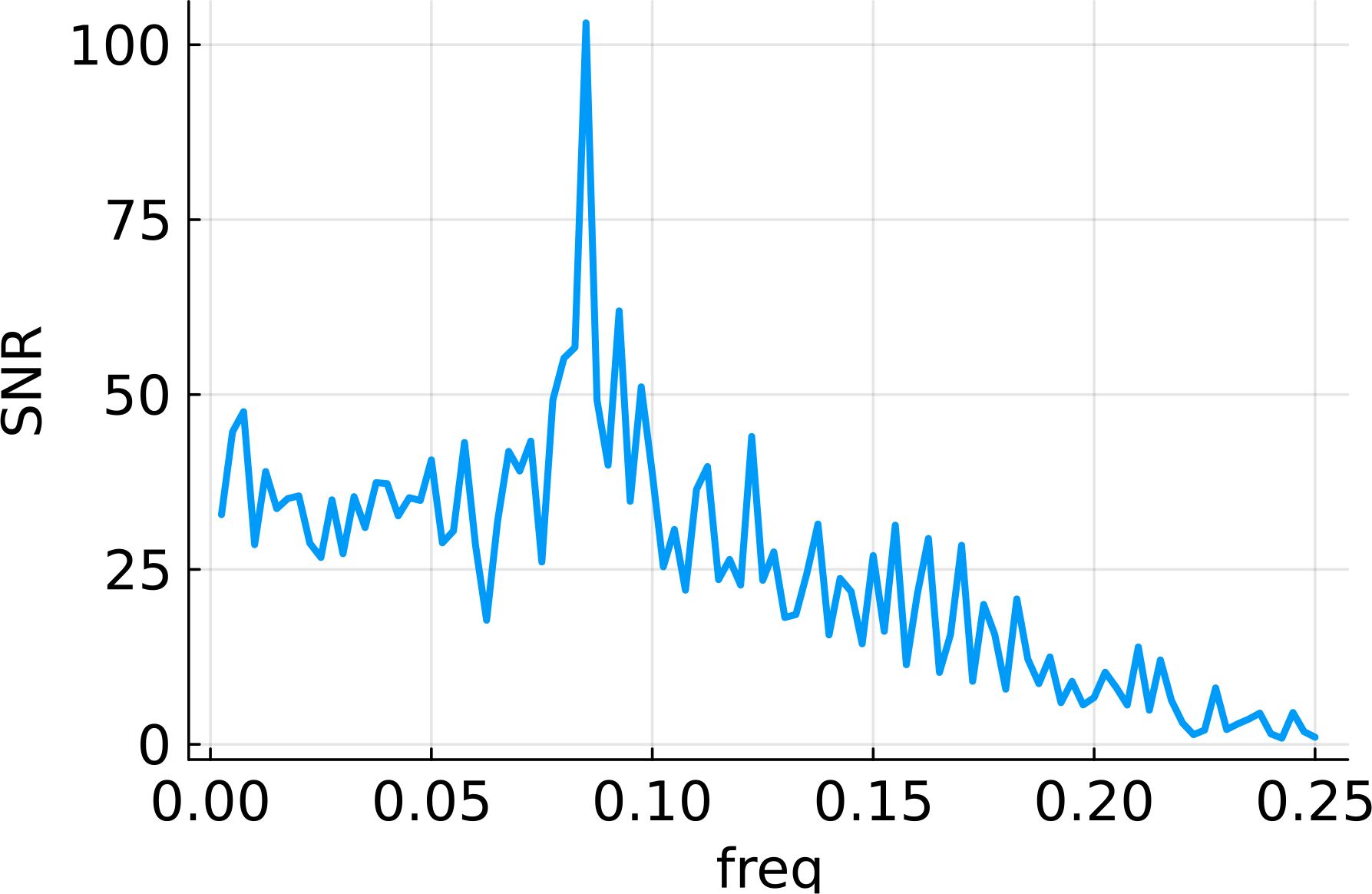

**Figure S6.**
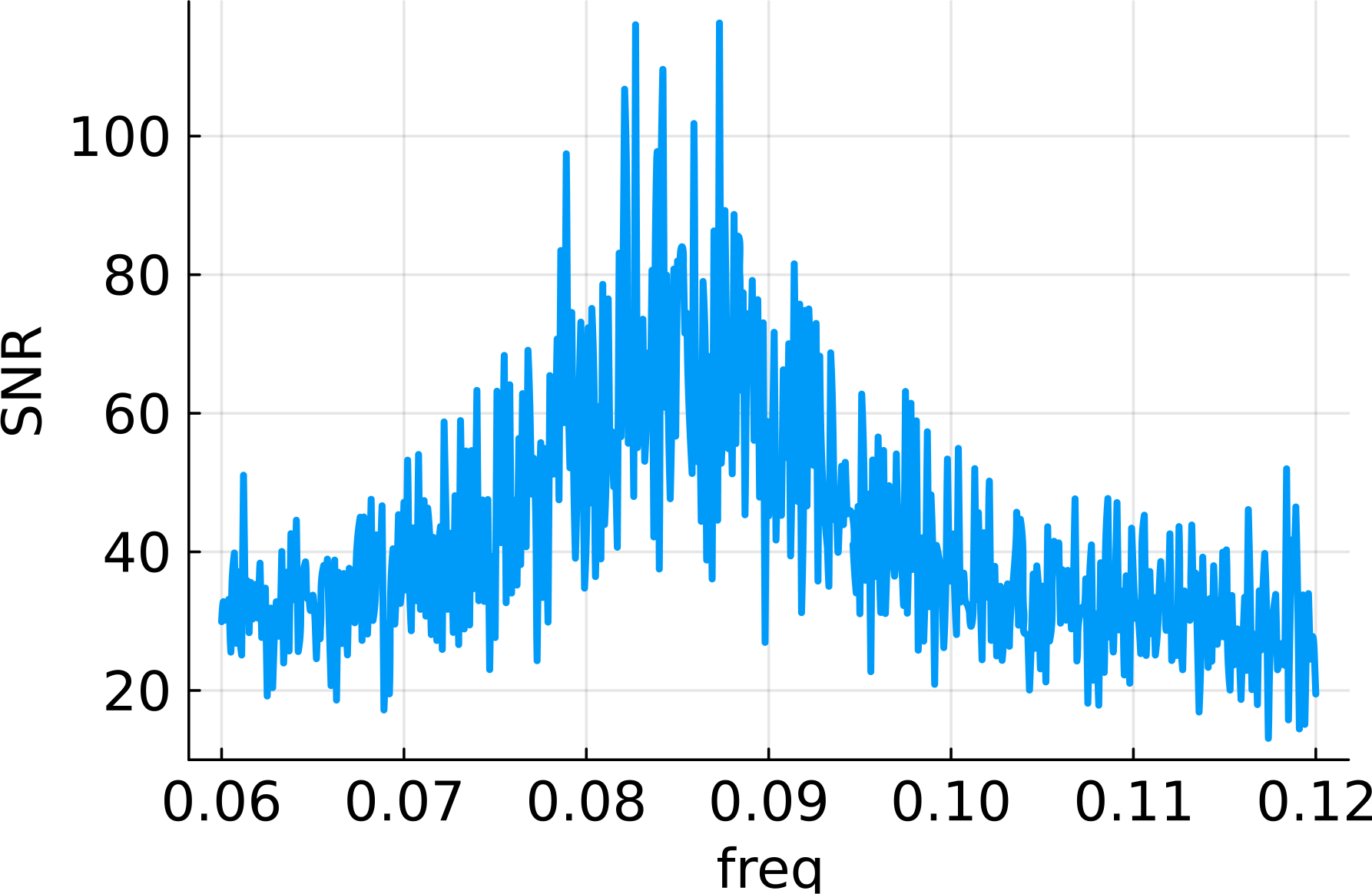

### Modelling experimental results with the FitzHugh-Nagumo system

Although we treat FHN as a toy model, it can be illustrative to show how well, and what aspects of data it can model.

We used equations (1),(2), and (3) to simulate the visibility of the target. When *v* was smaller than parameter *ϑ*, the target was considered visible and was considered invisible otherwise. We used Julia language to implement the Euler method with a time constant of 10 ms, and 1792 cuda threads. Each thread simulated 1638.4sec of time evolution. The switching statistics (visible and invisible periods) were collected from all threads and an empirical cumulative distribution function was constructed. We optimized the squared difference between the experimental and simulated empirical cumulative functions, which were evaluated at the same time points by linear interpolation. After extensive optimization using simulated annealing, there is still no guarantee that the set of parameters are optimal. It appears, based on simulation experience, that there are domains of parameters that define a qualitatively different behavior of the model and a phase diagram of these domains is not simple. For example, the FHN model can produce both a single mode distribution of visibility intervals and a multimodal distribution (compare Observer 1’s conditions T=1sec and T=4sec). It can also produce single and double mode distributions of invisible periods (compare, for example, for Observer 3 any moving mask condition with Observer 2’s T=1sec condition). Overall, the model can reproduce the gist of the data; however, it fails to reproduce fine-grained details.

Next, the simulation results are presented for each observer and for each condition. The graph for each condition consists of 5 panels. The left panel shows the graphical representation of the model – 2 nullclines and the location of a threshold. The two panels in the middle represent empirical cumulative functions for a duration of visibility periods (top) and for a duration of invisibility periods (bottom). The right panel represents the histogram of the durations of the visibility periods (top) and the invisibility periods (bottom). Since the FHN system is presented here as a toy model, only qualitative fits are expected and therefore we do not present any statistics quantifying the goodness of fit (although there are some very good ones).

Observer 1:

| Condition | Troxler | Static mask | T= 1 sec | T= 2 sec | T= 4 sec | T= 8 sec |
| --- | --- | --- | --- | --- | --- | --- |
| a | 2.126949 | 2.537207 | 0.401874 | 0.282345 | 1.031162 | 1.152413 |
| b | -1.06559 | 0.075527 | -2.00085 | -2.04071 | -1.48359 | -1.90866 |
| c | -0.94236 | -0.6744 | -1.34693 | -1.85308 | -1.22197 | -1.39463 |
| d | -0.70606 | -0.96372 | 0.007982 | -0.27136 | 0.239158 | -0.28279 |
| f | 0.108981 | 0.311872 | 0.215763 | 0.14434 | 0.703961 | 0.176807 |
| g | -0.16937 | -0.42574 | -0.27444 | -0.38791 | 0.572244 | 0.317861 |
| $\tau_1$ | 0.123897 | 0.31495 | 0.232296 | 0.410983 | 0.334035 | 0.200963 |
| $\tau_2$ | 1.485418 | 1.305269 | 1.381106 | 1.484369 | 2.162105 | 0.931568 |
| $\mathcal{G}$ | 0.663432 | 0.854217 | -0.35624 | -0.47543 | 0.872134 | 0.915573 |
| A | 0.156968 | 0.444717 | 0.282998 | 0.614291 | 0.321997 | 0.579956 |
| $\omega$ | 0 | 0 | 6.283185 | 3.141593 | 1.570796 | 0.785398 |
| $\varepsilon_v$ | 0.01732 | 0.03471 | 0.043101 | 0.01815 | 0.00443 | 0.03354 |
| $\varepsilon_w$ | 1.128973 | 0.666719 | 0.738909 | 0.945618 | 0.316718 | 0.772264 |

**Figure S7 1.**
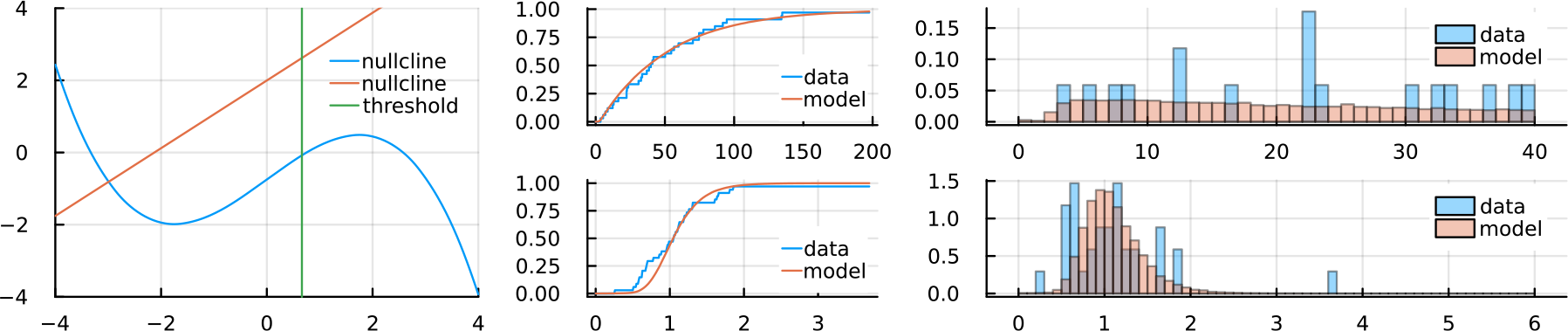
Observer 1, Troxler condition

**Figure S7 2.**
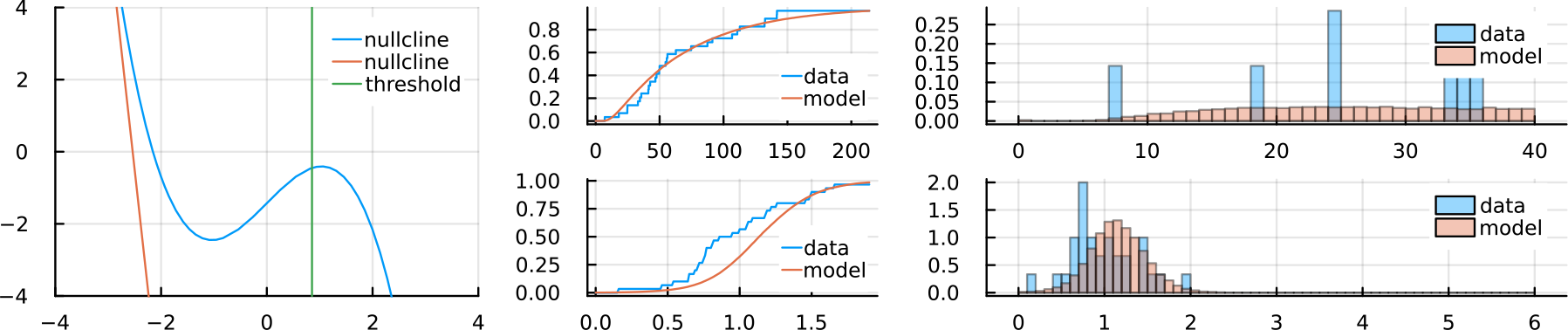
Observer 1, fixed mask condition.

**Figure S7 3.**
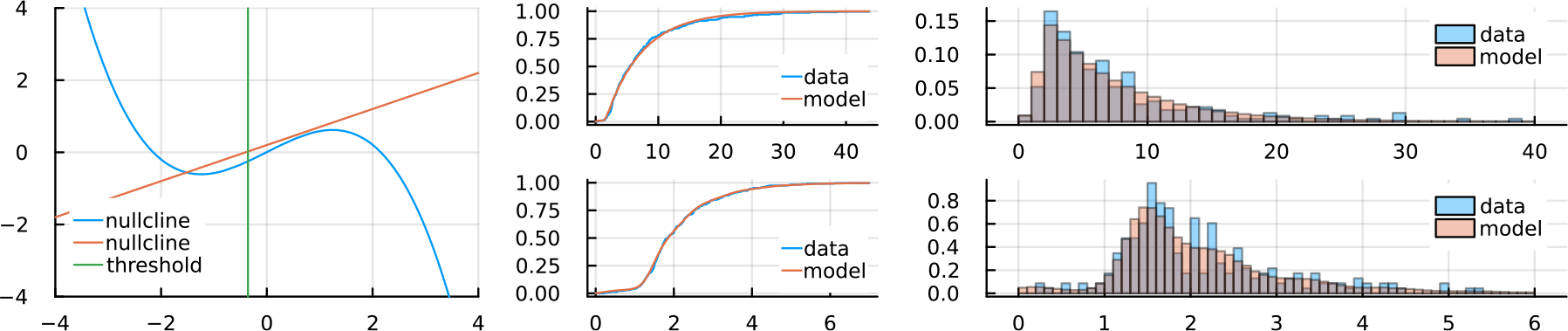
Observer 1, mask period 1 sec.

**Figure S7 4.**
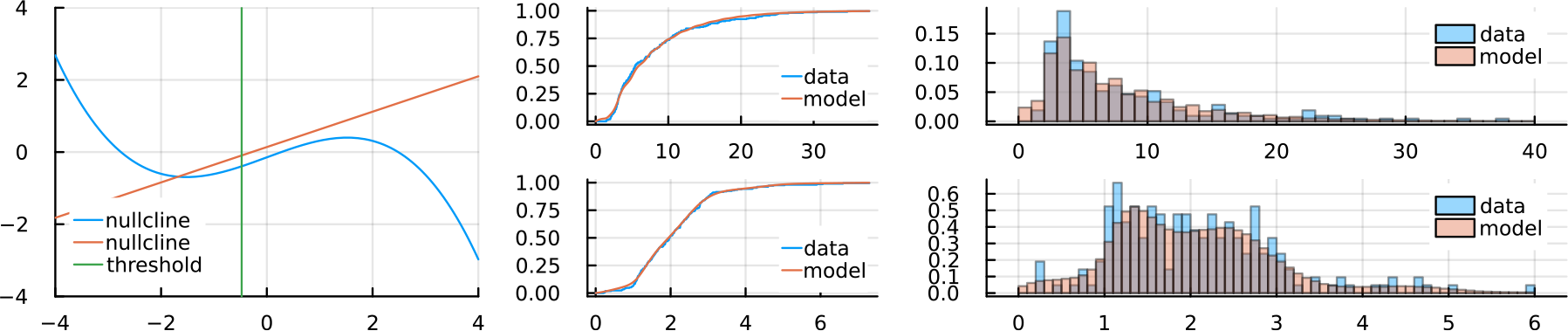
Observer 1, mask period 2 sec

**Figure S7 5.**
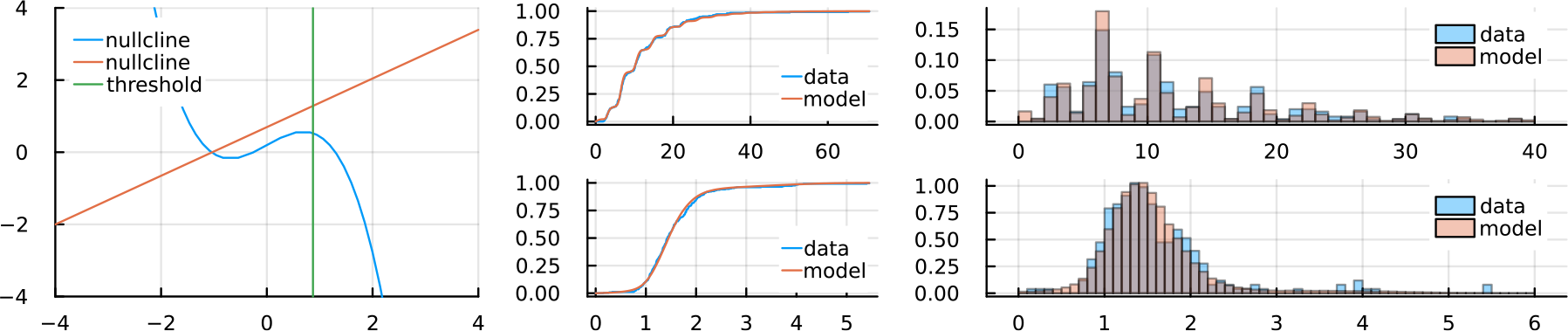
*Observer 1, mask period 4 sec*

**Figure S7 6.**
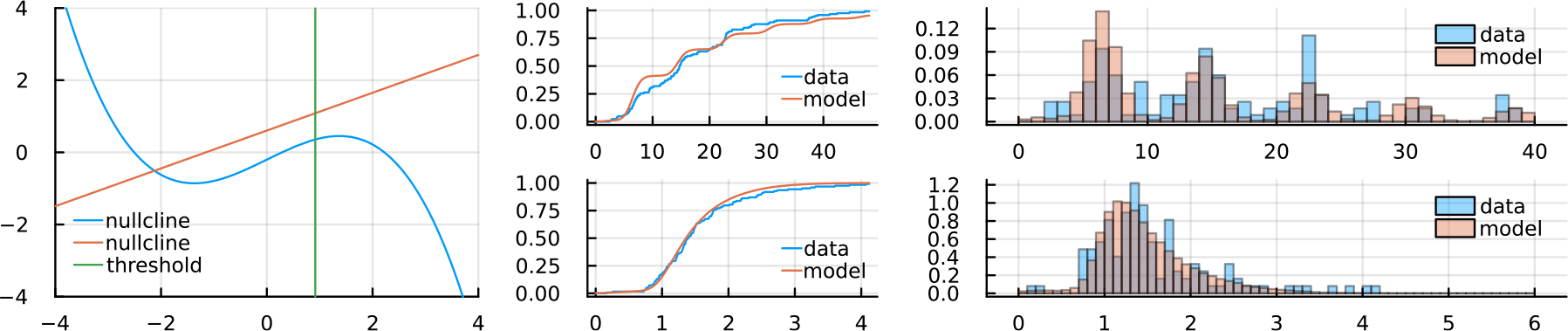
*Observer 1, mask period 8 sec*

Observer 2:

Observer 2
| Condition | Troxler | Static mask | T= 1 sec | T= 2 sec | T= 4 sec | T= 8 sec |
| --- | --- | --- | --- | --- | --- | --- |
| a | 1.281363 | 1.73984 | 0.47692 | 1.386709 | 1.764775 | 1.026208 |
| b | -1.60968 | -2.13624 | -2.01798 | -1.37222 | -1.67457 | -1.62193 |
| c | -1.18784 | -1.28864 | -1.0043 | -1.94192 | -2.11952 | -1.09593 |
| d | -0.25777 | -0.44118 | -0.2692 | -0.03512 | -1.62677 | -0.50866 |
| f | 0.312004 | 0.521641 | 0.693827 | 0.265933 | 0.400128 | 0.781429 |
| g | -0.69694 | -0.16719 | -0.52065 | 0.024041 | 0.244929 | -0.48591 |
| $\tau_1$ | 0.43201 | 0.668503 | 0.066665 | 0.655951 | 0.388831 | 0.133596 |
| $\tau_2$ | 0.519698 | 0.467045 | 1.944515 | 1.497732 | 2.29112 | 1.79797 |
| $\mathcal{G}$ | 0.391573 | 0.325812 | 0.188097 | 0.229121 | 0.920289 | 0.301037 |
| A | 0.23794 | 0.28797 | 0.365738 | 1.431454 | 1.02518 | 0.302424 |
| $\omega$ | 0 | 0 | 6.283185 | 3.141593 | 1.570796 | 0.785398 |
| $\mathcal{E}_v$ | 0.02476 | 0.12558 | 0.146495 | 0.07157 | 0.013362 | 0.0487 |
| $\mathcal{E}_w$ | 3.585802 | 3.69517 | 0.936421 | 0.993511 | 1.583296 | 1.035695 |

**Figure S7 7.**
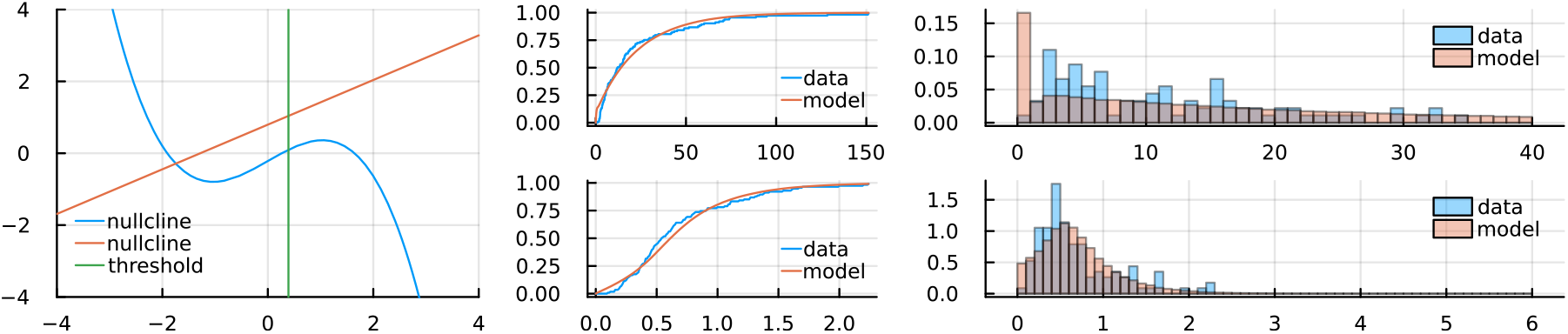
*Observer 2, Troxler condition*

**Figure S7 8.**
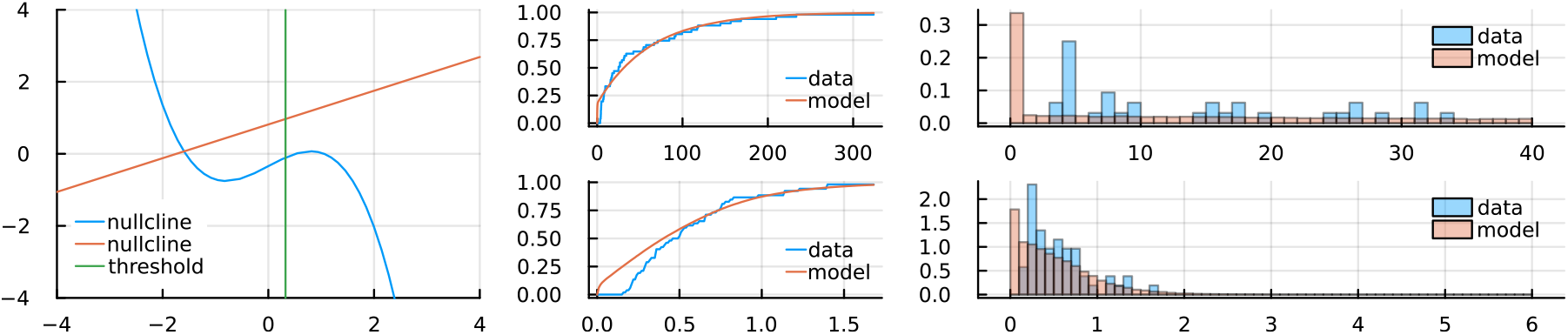
*Observer 2, static mask*

**Figure S7 9.**
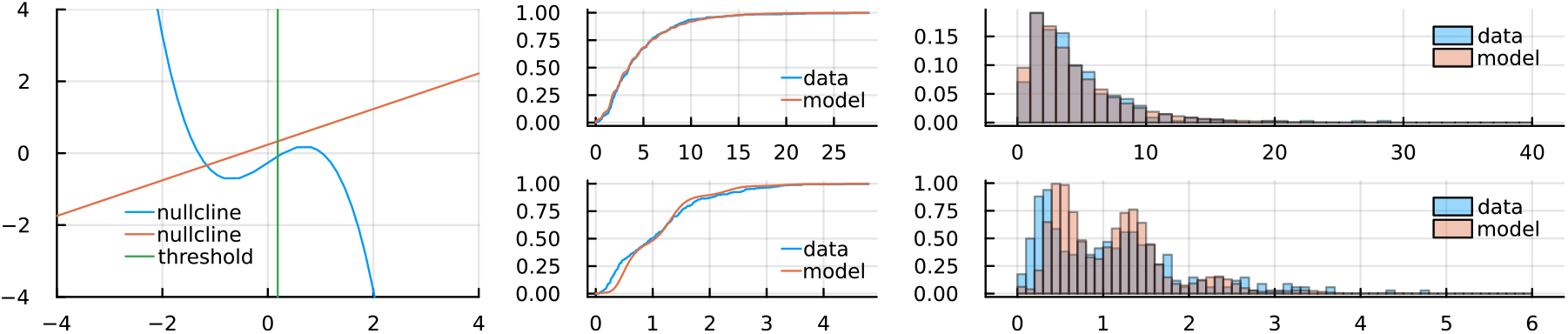
*Observer 2, mask period 1 sec*

**Figure S7 10.**
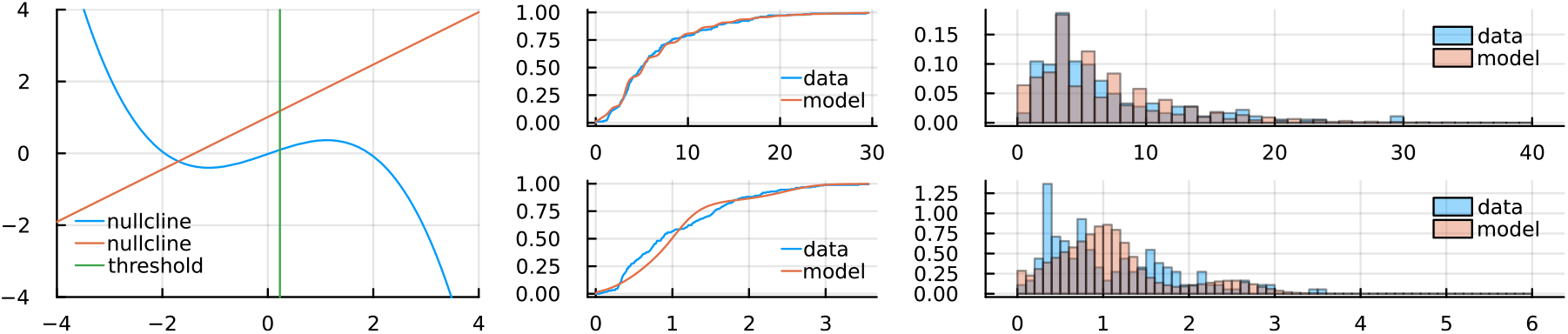
*Observer 2, mask period 2 sec*

**Figure S7 11.**
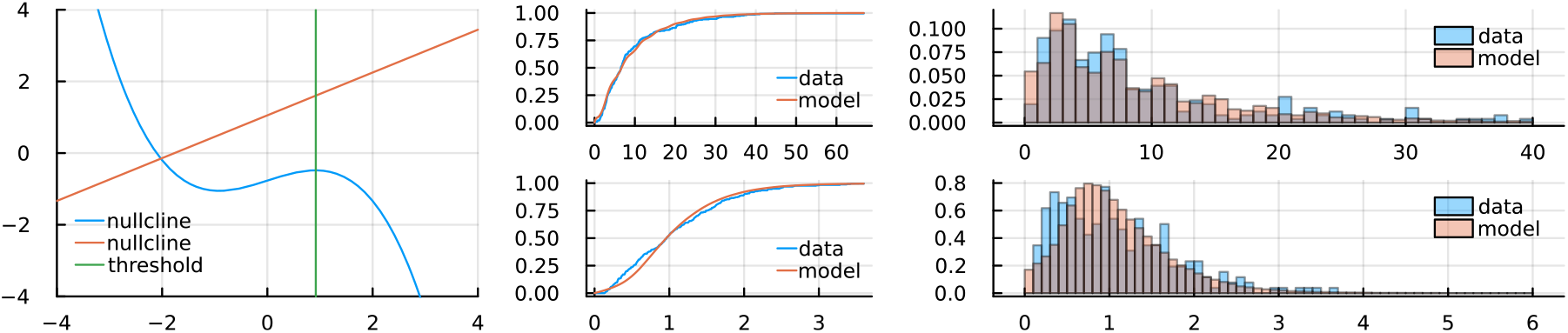
*Observer 2, mask period 4 sec*

**Figure S7 12.**
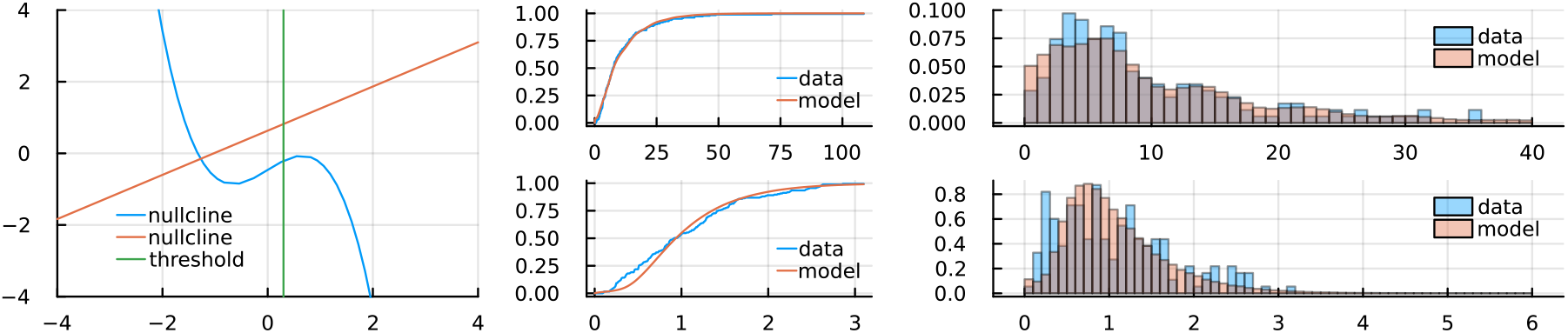
*Observer 2, mask period 8 sec*

Observer 3:

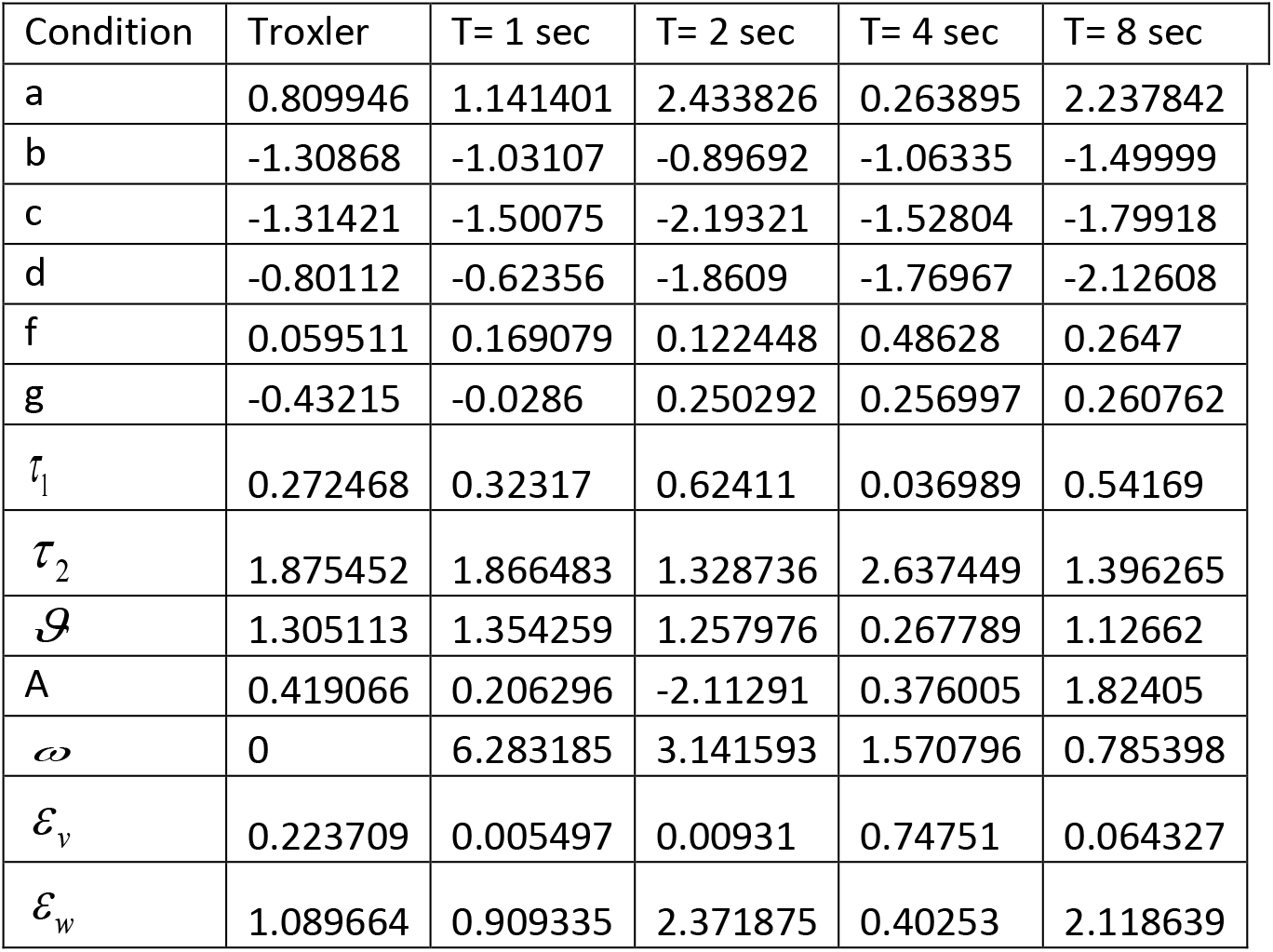

**Figure S7 13.**
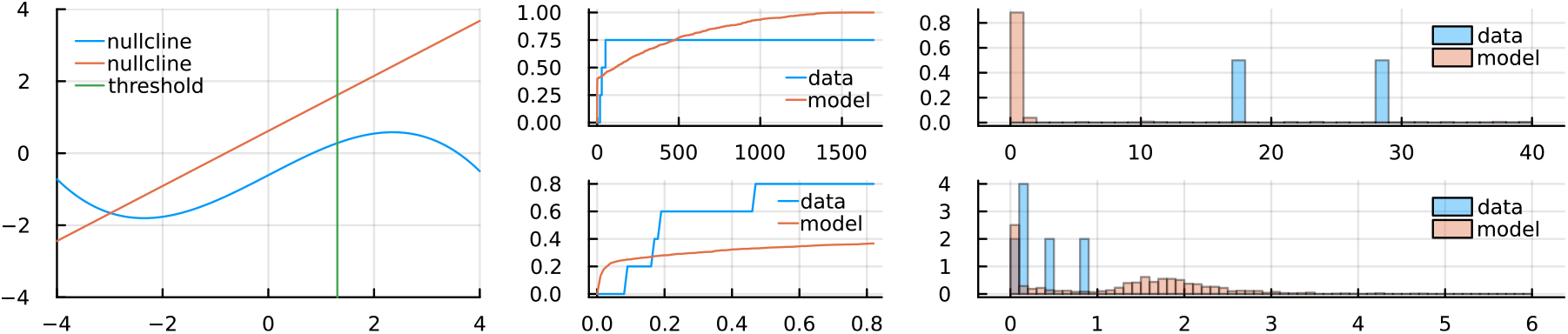
*Observer 3, Troxler condition*

**Figure S7 14.**
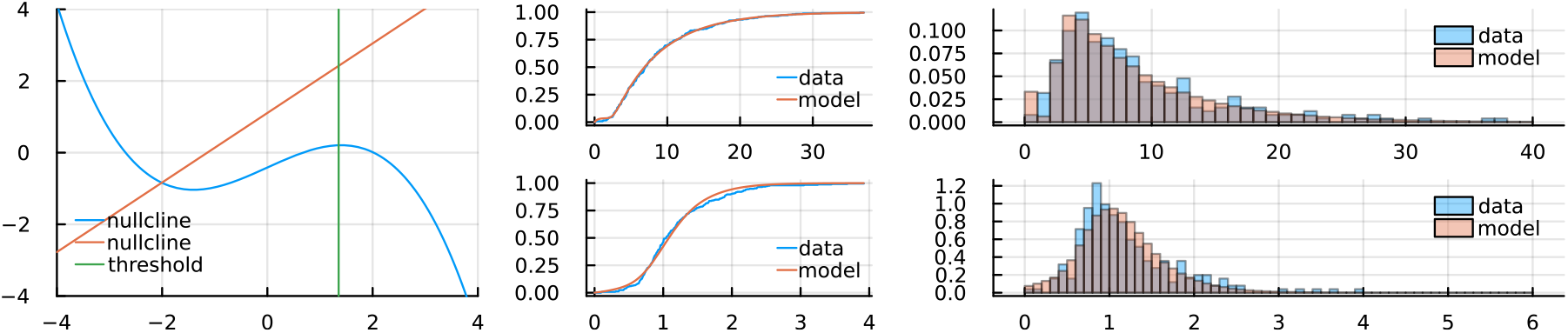
*Observer 3, mask period 1 sec*

**Figure S7 15.**
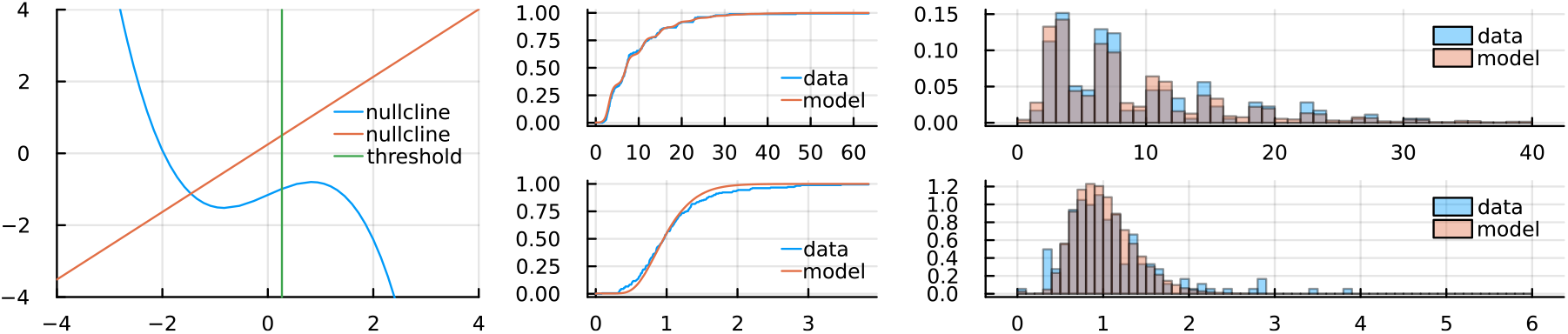
*Observer 3, mask period 2 sec*

**Figure S7 16.**
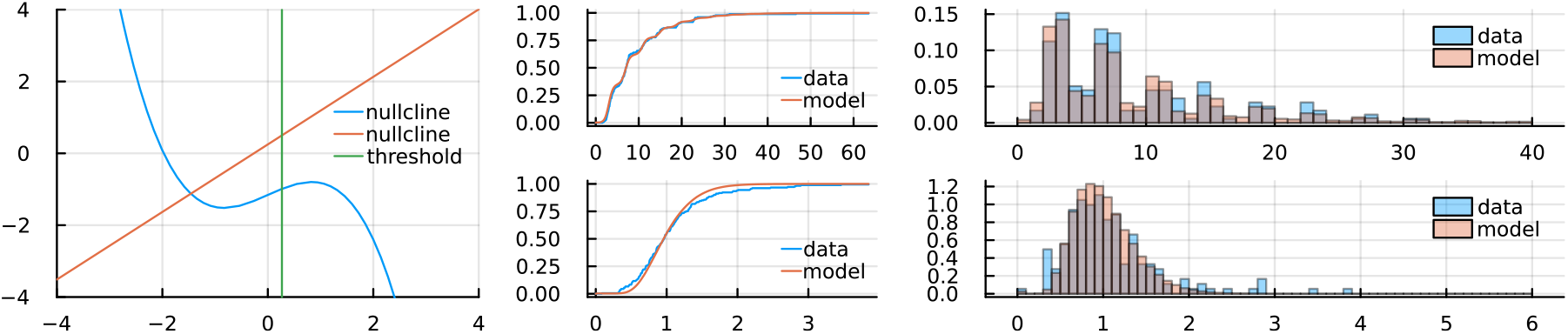
*Observer 3, mask period 4 sec*

**Figure S7 17.**
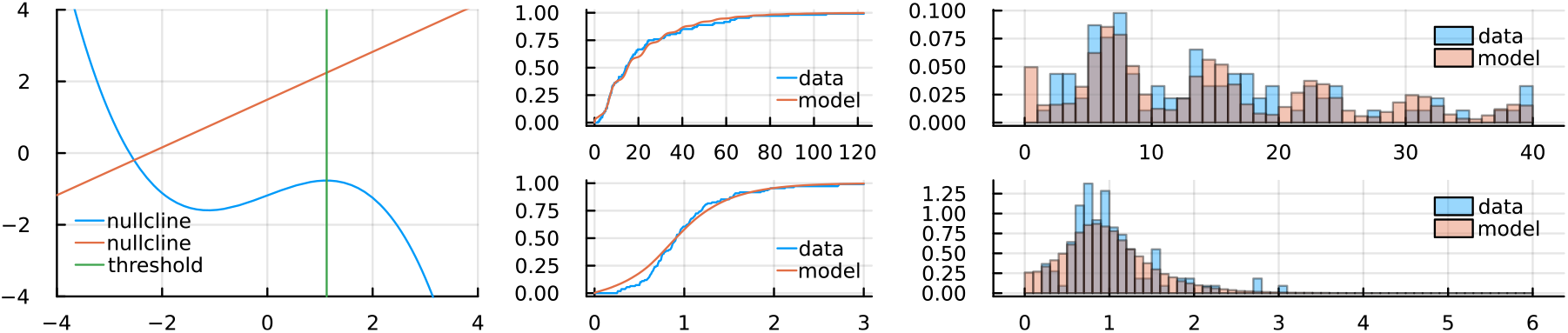
*Observer 3, mask period 8 sec*

Observer 4:

Observer 4
| Condition | Troxler | Static mask | T= 1 sec | T= 2 sec | T= 4 sec | T= 8 sec |
| --- | --- | --- | --- | --- | --- | --- |
| a | 1.026908 | 1.106662 | 1.606869 | 0.633064 | 1.196692 | 1.301805 |
| b | -0.48727 | -0.12017 | -0.22013 | -0.73089 | -0.45256 | -0.39631 |
| c | -0.97979 | -0.91179 | -1.90484 | -0.98734 | -1.10936 | -1.00158 |
| d | -0.05874 | -0.21817 | -0.8302 | -0.83492 | -1.00466 | -0.157 |
| f | 0.472557 | 0.556036 | 0.513884 | 0.479757 | 0.527935 | 0.368404 |
| g | -0.26817 | 0.602486 | -0.75682 | -0.06794 | -0.5626 | 0.039258 |
| $\tau_1$ | 0.50526 | 0.333002 | 0.055508 | 0.042189 | 0.033833 | 0.054194 |
| $\tau_2$ | 0.453373 | 0.605014 | 1.947667 | 1.520491 | 1.372959 | 1.389843 |
| $\mathcal{G}$ | 0.909866 | 1.380892 | -0.07187 | -0.16036 | -0.03143 | -0.38733 |
| A | 0.010446 | 0.659827 | 0.09758 | 0.317599 | 0.527176 | -0.27219 |
| $\omega$ | 0 | 0 | 6.283185 | 3.141593 | 1.570796 | 0.785398 |
| $\mathcal{E}_v$ | 0.00169 | 0.006491 | 0.96616 | 0.6958 | 0.585388 | 0.413851 |
| $\mathcal{E}_w$ | 2.112738 | 2.172679 | 0.99561 | 1.094958 | 1.296876 | 1.287108 |

**Figure S7 18.**
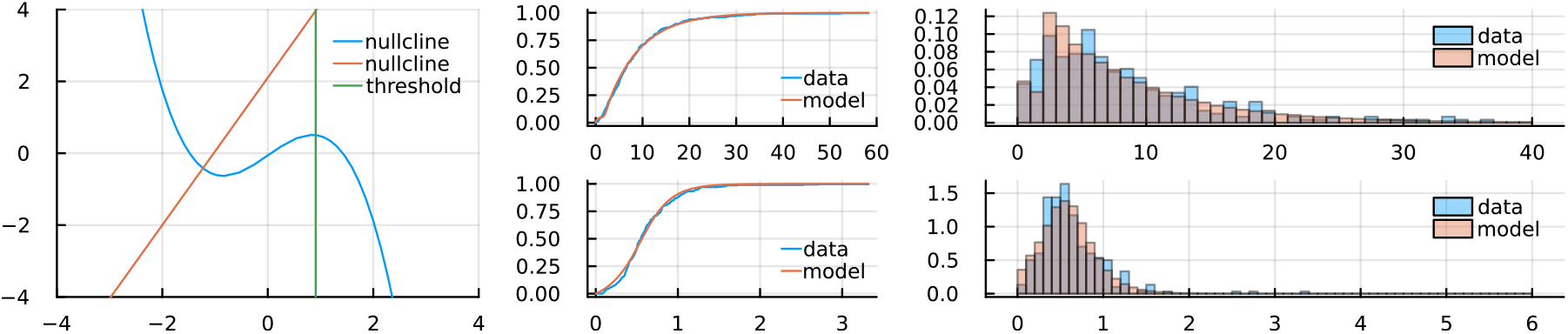
*Observer 4, Troxler condition*

**Figure S7 19.**
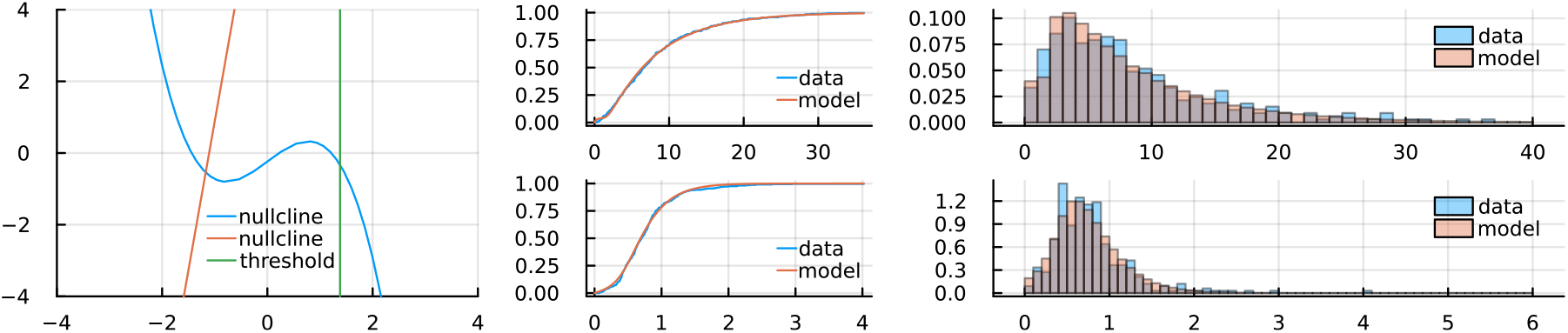
*Observer 4, static mask*

**Figure S7 20.**
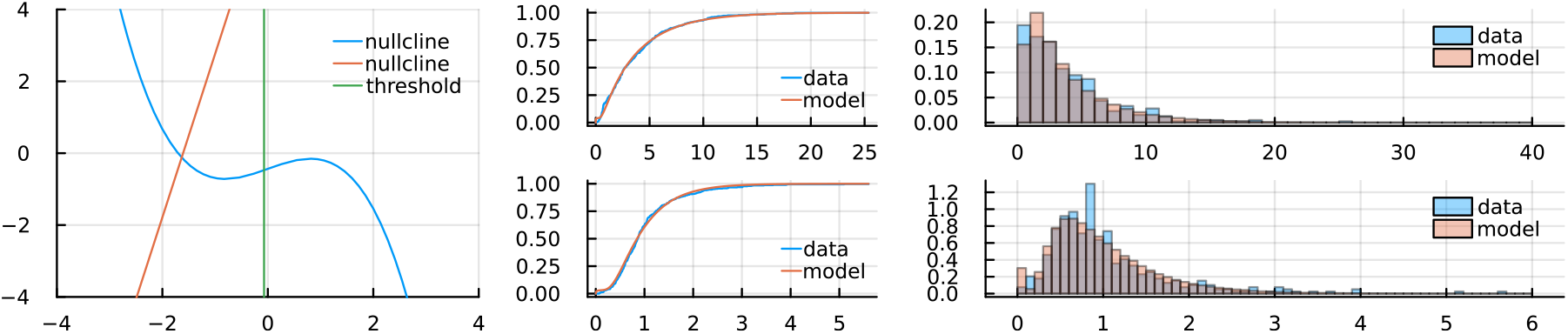
*Observer 4, mask period 1 sec*

**Figure S7 21.**
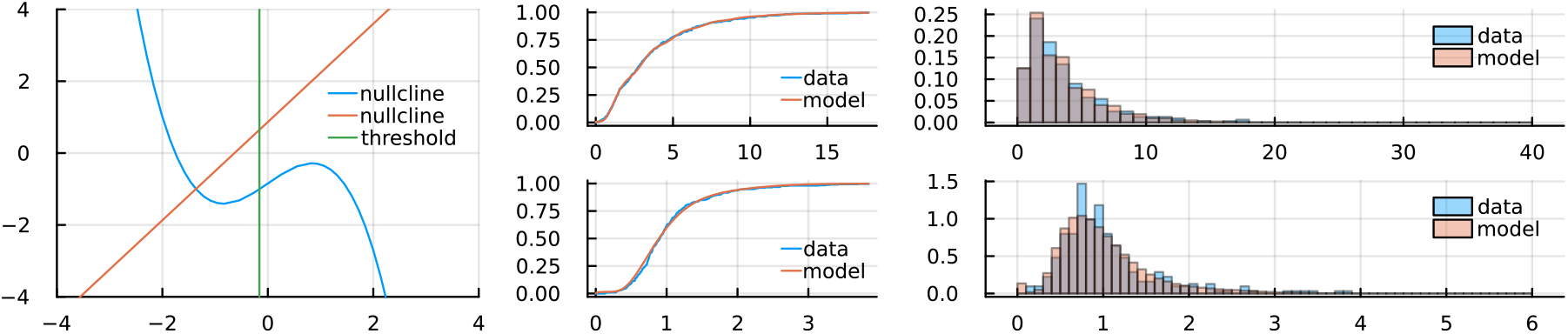
*Observer 4, mask period 2 sec*

**Figure S7 22.**
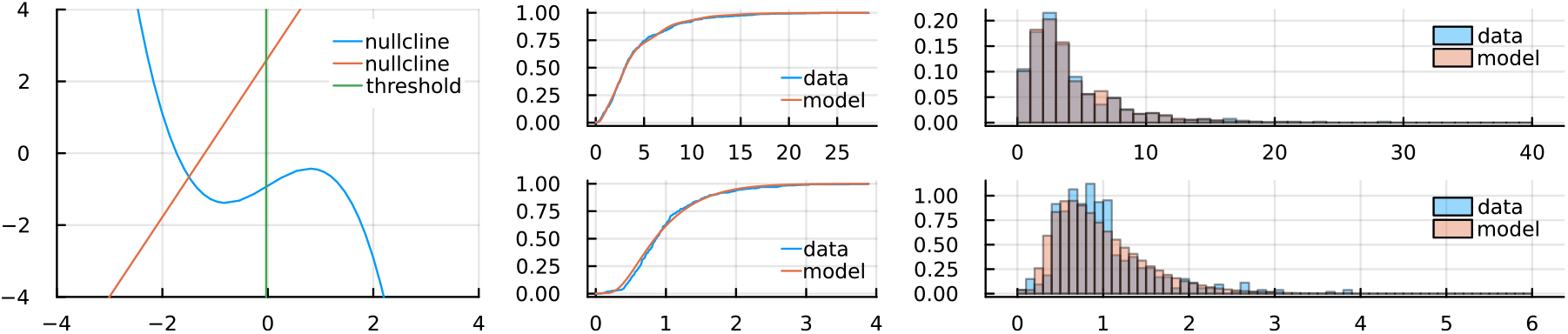
*Observer 4, mask period 4 sec*

**Figure S7 23.**
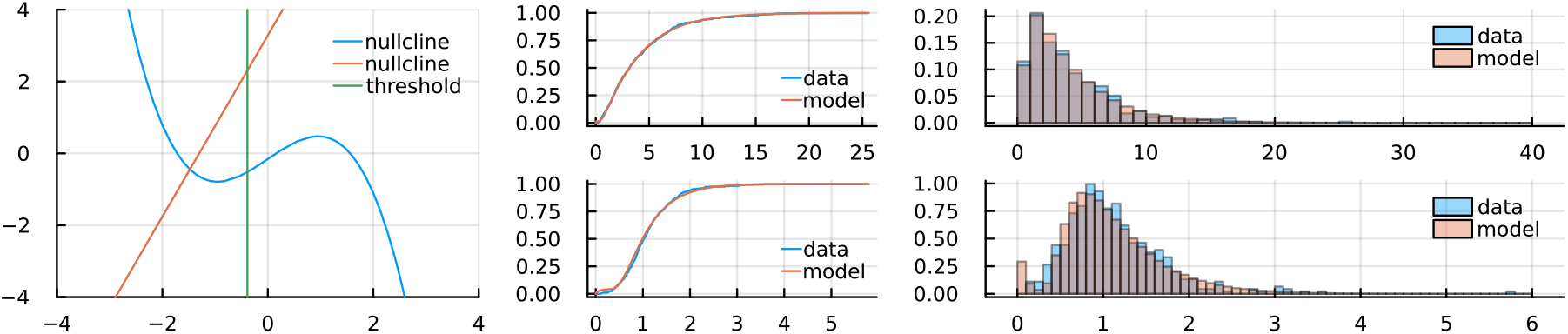
*Observer 4, mask period 8 sec*

Observer 5:

| Condition | Troxler | Static mask | T= 1 sec | T= 2 sec | T= 4 sec | T= 8 sec |
| --- | --- | --- | --- | --- | --- | --- |
| a | 1.196467 | 1.720968 | 1.443092 | 0.085195 | 1.278365 | 1.13296 |
| b | -1.39322 | 0.066738 | -1.10594 | -1.82323 | -1.55184 | -1.41153 |
| c | -1.15404 | -0.53324 | -1.03932 | -2.02145 | -1.01456 | -1.58057 |
| d | -0.23163 | -0.17772 | 0.504252 | -0.65488 | 0.242235 | -0.38216 |
| f | 0.034837 | 0.054514 | 0.67193 | 0.253125 | 0.461904 | 0.324195 |
| g | -0.32562 | 0.610691 | -0.58135 | -0.58692 | -0.75821 | 0.467891 |
| $\tau_1$ | 0.034011 | 0.246963 | 0.113909 | 0.12637 | 0.419616 | 0.120938 |
| $\tau_2$ | 0.828991 | 0.212921 | 1.309333 | 2.17649 | 1.633262 | 1.849463 |
| $\mathcal{G}$ | 0.053378 | 0.42426 | 0.056509 | 0.508063 | -0.14822 | 0.745169 |
| A | 0.33543 | 0.45485 | 0.058331 | 0.352975 | 1.345371 | 0.520362 |
| $\omega$ | 0 | 0 | 6.283185 | 3.141593 | 1.570796 | 0.785398 |
| $\mathcal{E}_v$ | 0.03528 | 0.08602 | 0.039454 | 0.0002 | 0.06644 | 0.168476 |
| $\mathcal{E}_w$ | 3.443979 | 3.058513 | 0.787748 | 0.116877 | 0.299801 | 0.465378 |

**Figure S7 24.**
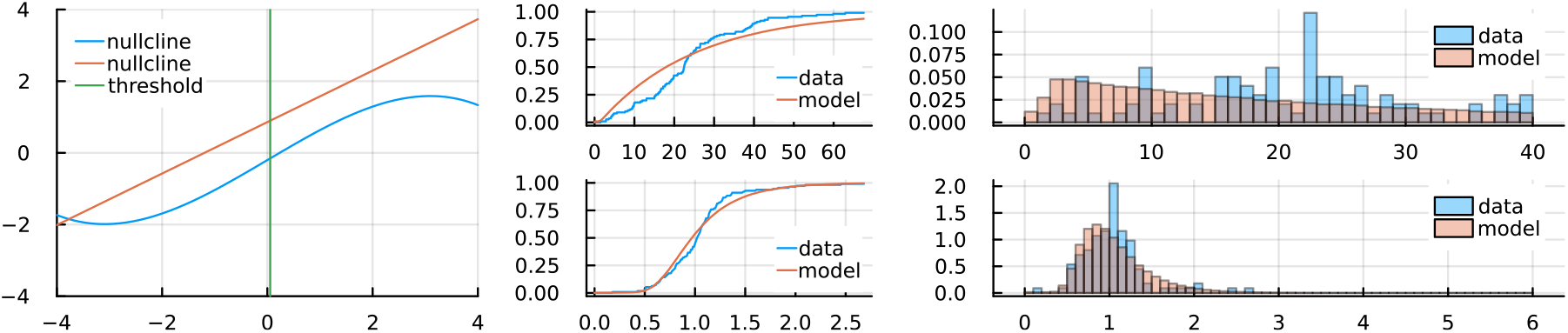
*Observer 5, Troxler condition*

**Figure S7 25.**
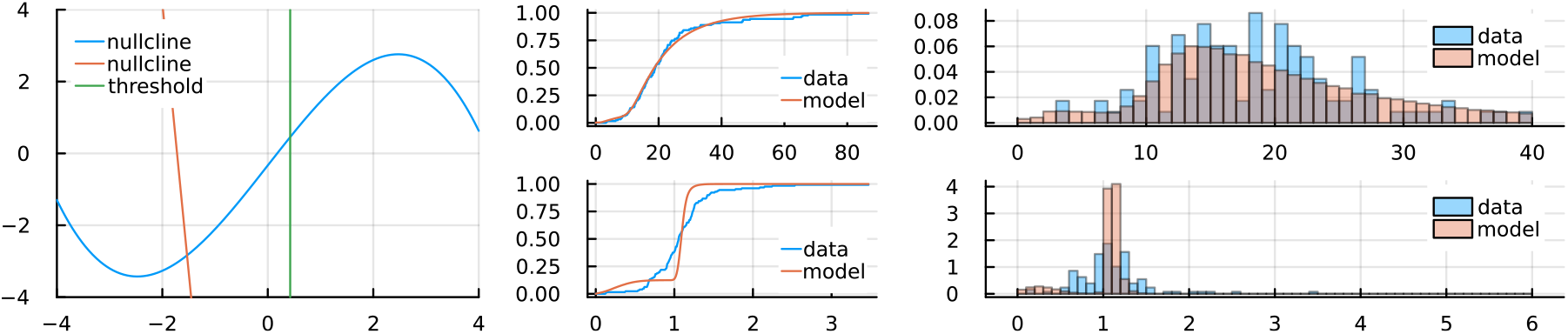
*Observer 5, static mask*

**Figure S7 26.**
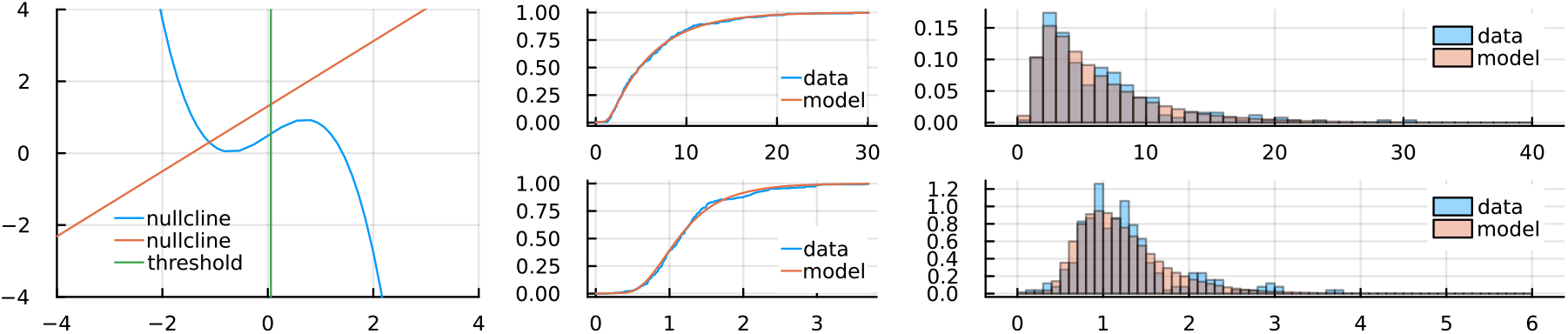
*Observer 5, mask period 1 sec*

**Figure S7 27.**
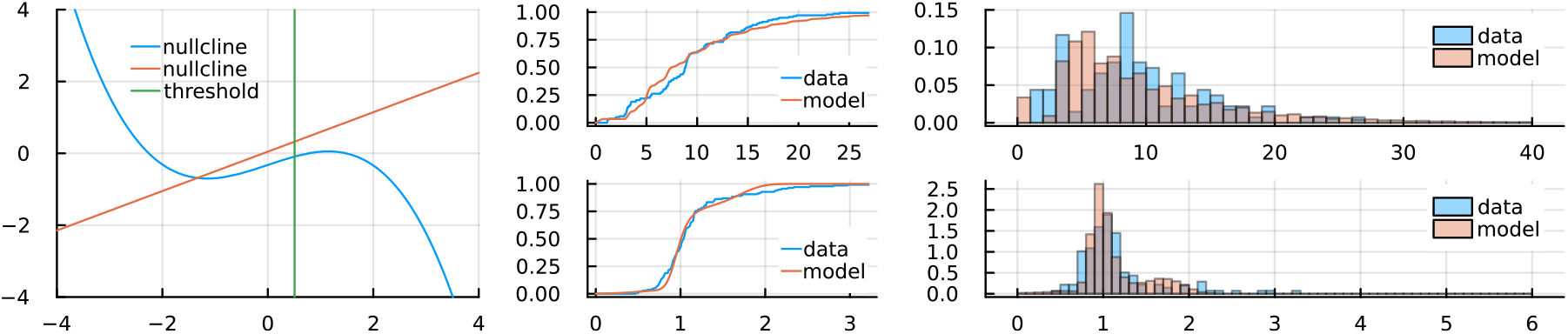
*Observer 5, mask period 2 sec*

**Figure S7 28.**
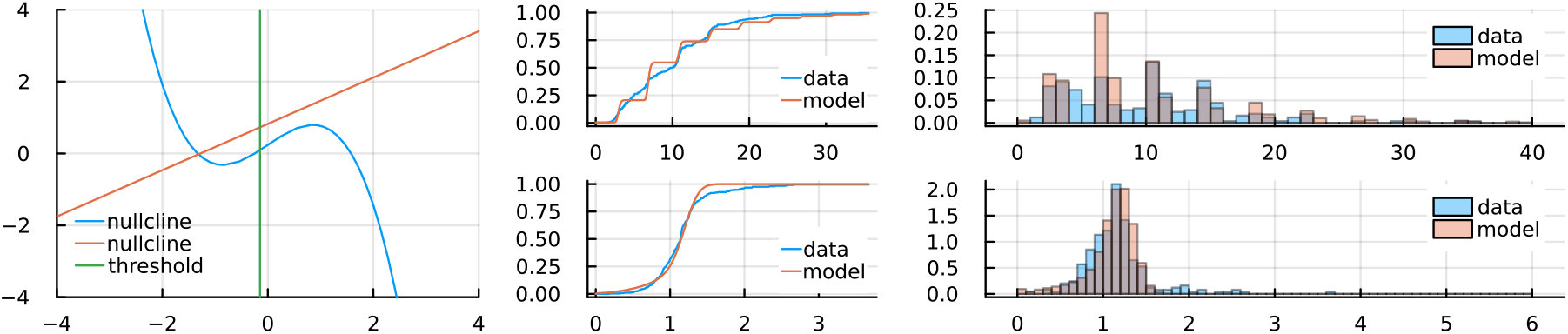
*Observer 5, mask period 4 sec*

**Figure S7 29.**
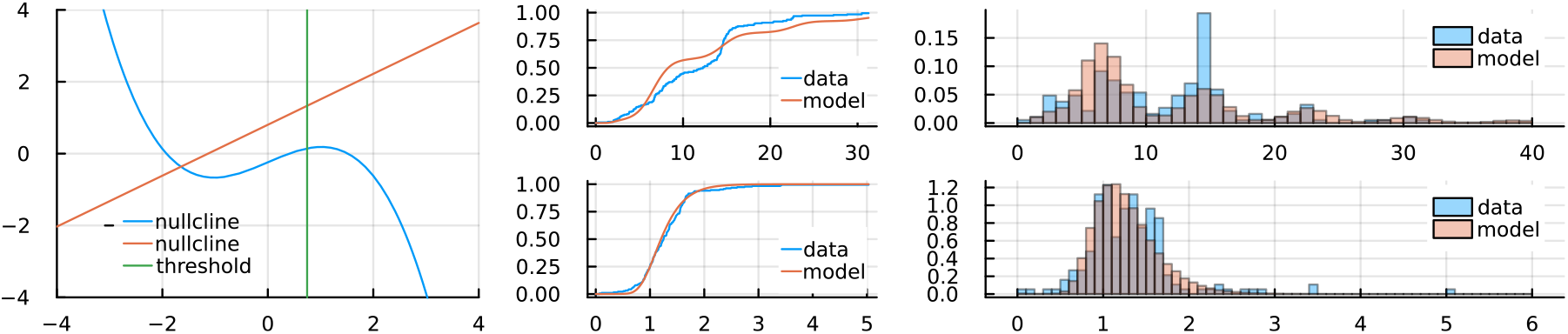
*Observer 5, mask period 8 sec*

Observer 6:

| Condition | Troxler | Static mask | T= 1 sec | T= 2 sec | T= 4 sec | T= 8 sec |
| --- | --- | --- | --- | --- | --- | --- |
| a | 1.38979 | 1.28393 | 0.559328 | 1.367742 | 1.574264 | 1.371861 |
| b | -1.53582 | -1.33379 | 0.22013 | -0.40545 | 0.104991 | -0.40351 |
| c | -0.92482 | -0.72756 | -2.62585 | -1.56167 | -1.37766 | -1.56023 |
| d | -0.19018 | -0.06896 | -0.2486 | -0.25519 | -0.3875 | -0.25863 |
| f | 0.117392 | 0.309216 | 0.18092 | 0.33483 | 0.75969 | 0.336832 |
| g | -0.5483 | -0.6866 | 0.624168 | -0.08291 | -0.72322 | -0.08189 |
| $\tau_1$ | 0.028988 | 0.151263 | 0.082068 | 0.24242 | 0.22669 | 0.249326 |
| $\tau_2$ | 0.667417 | 0.394121 | 1.430578 | 1.905523 | 1.74512 | 1.906353 |
| $\mathcal{G}$ | 0.205324 | 0.45658 | 1.466674 | 0.934116 | 0.998277 | 0.935328 |
| A | 0.24777 | 0.03333 | 0.55002 | 2.130714 | 2.139876 | 2.129533 |
| $\omega$ | 0 | 0 | 6.283185 | 3.141593 | 1.570796 | 0.785398 |
| $\mathcal{E}_v$ | 0.10146 | -0.18535 | 0.253672 | 0.014673 | 0.01997 | 0.01296 |
| $\mathcal{E}_w$ | 3.414567 | 3.38991 | 2.10691 | 1.692612 | 1.088045 | 1.689976 |

**Figure S7 30.**
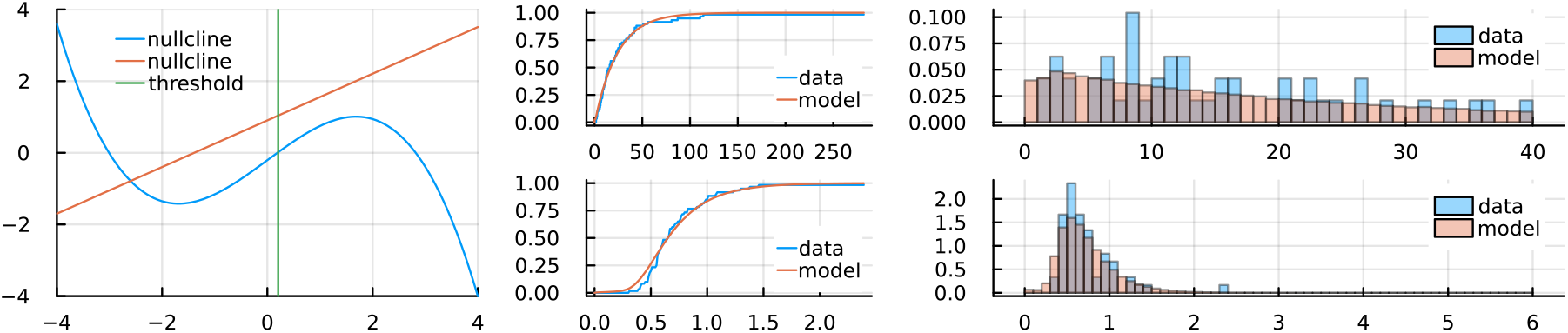
*Observer 6, Troxler condition*

**Figure S7 31.**
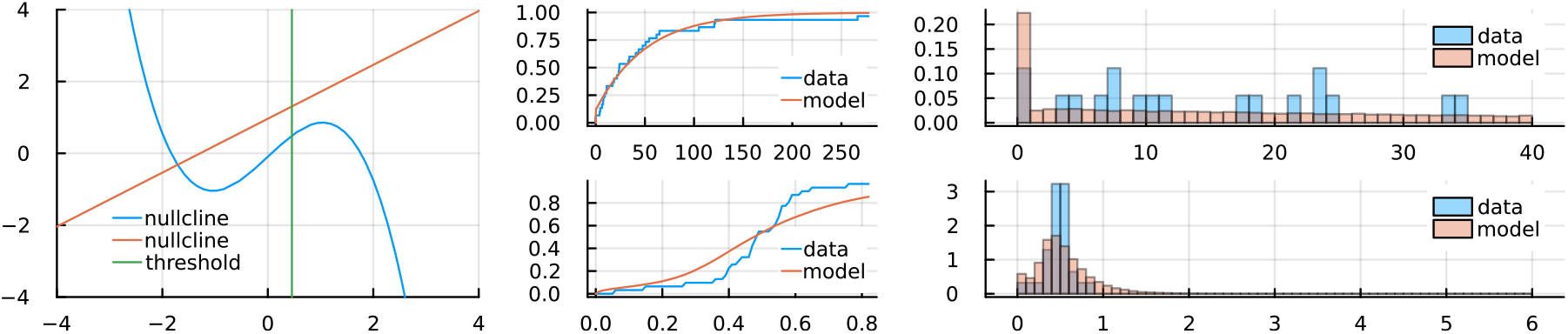
*Observer 6, static mask*

**Figure S7 32.**
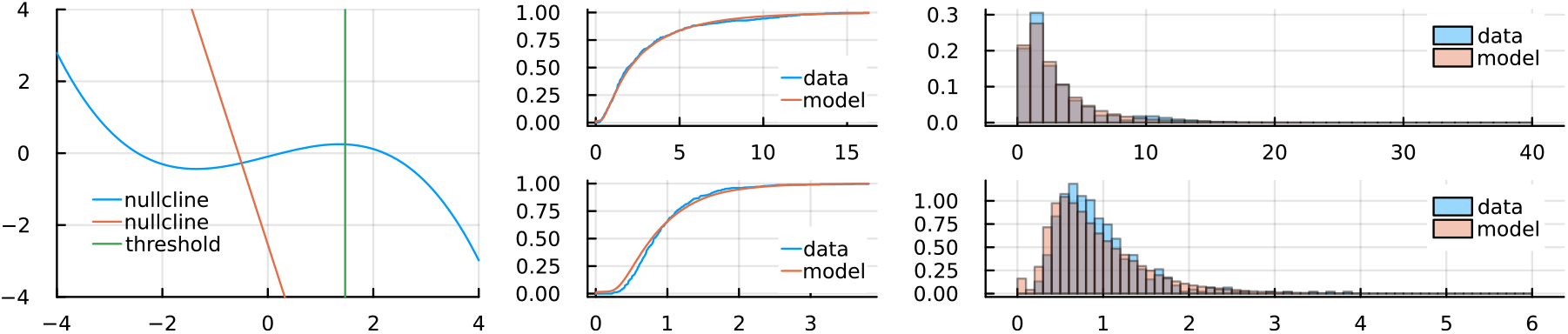
*Observer 6, mask period 1 sec*

**Figure S7 33.**
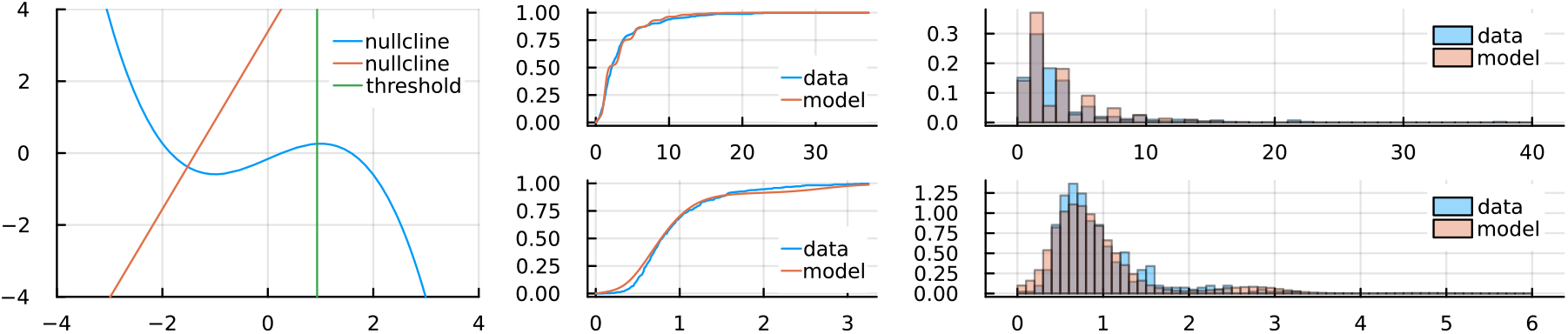
*Observer 6, mask period 2 sec*

**Figure S7 34.**
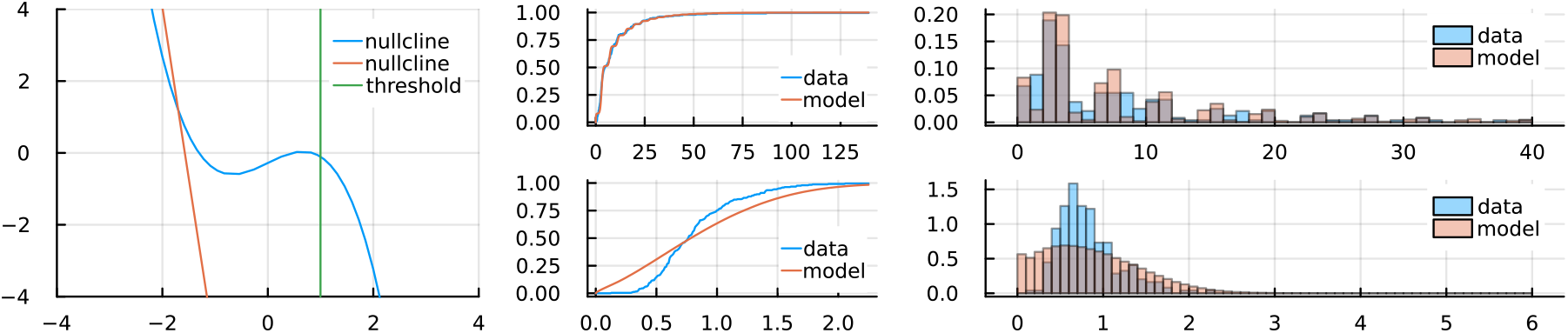
*Observer 6, mask period 4 sec*

**Figure S7 35.**
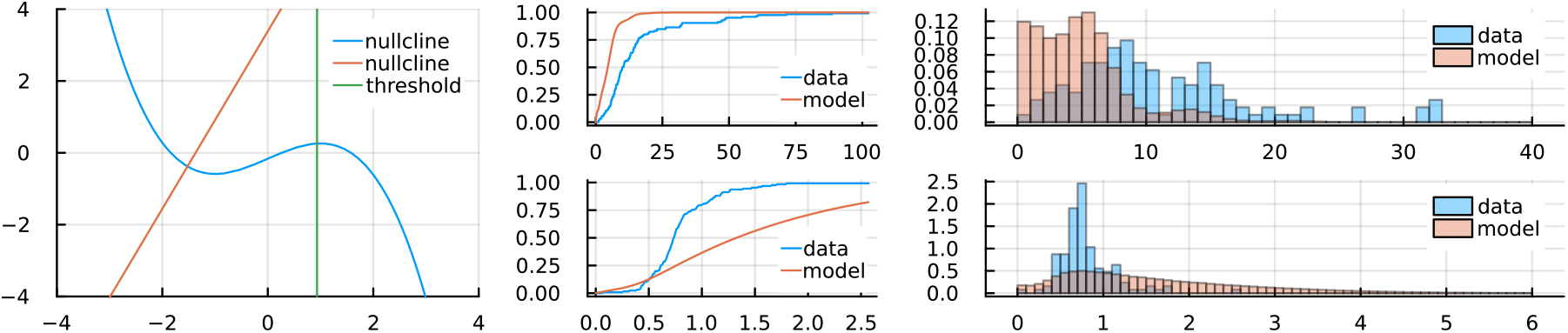
*Observer 6, mask period 8 sec*

